# Theoretical Principles of Enhancer-Promoter Communication in Transcriptional Bursting

**DOI:** 10.1101/2022.01.24.477520

**Authors:** Zihao Wang, Zhenquan Zhang, Songhao Luo, Tianshou Zhou, Jiajun Zhang

**Affiliations:** Guangdong Province Key Laboratory of Computational Science School of Mathematics, Sun Yat-sen University, Guangzhou 510275, P. R. China; School of Mathematics, Sun Yat-Sen University, Guangzhou 510275, P. R. China

## Abstract

Transcriptional regulation occurs through genomic contacts between enhancers and their cognate promoters, and most genes are transcribed in a bursty fashion. To understand the relationship between these two phenomena, we develop a general modeling framework in terms of the information transmission from upstream genomic organization to downstream transcriptional bursting. Importantly, we uncover fundamental theoretical principles of enhancer-promoter (E-P) spatial communication in the modulation of transcriptional burst size (BS) and burst frequency (BF). First, BS and BF obey their respective power-law dependences on the E-P communication strength and distinct scaling exponents. Second, the E-P spatial distance follows a Maxwell-Boltzmann distribution rather than the previously assumed Gauss distribution. Third, the E-P genomic distance affects transcriptional outcomes biphasically (i.e., in an exponential decay for small E-P genomic distances but insensitively to large E-P genomic distances). Fourth, the E-P communication mainly modulates BF rather than BS. Finally, the mutual information between BS (or BF) and E-P spatial distance further reveals essential characteristics of the information transfer from the upstream to the downstream. Our predictions are experimentally verifiable, e.g., confirmed by experimental data on *Drosophila*. The overall analysis provides insights into the role of the E-P communication in the control of transcriptional bursting.

**Significance:** Measurement technologies of chromatin conformations and genome-wide occupancy data of architectural proteins have revealed that genome topology is tightly intertwined with gene transcription. However, a long-standing question in transcriptional regulation is how the enhancer-promoter (E-P) spatial communication impacts transcriptional bursting kinetics. To address this issue, we develop a multiscale model that couples upstream chromatin dynamics to downstream transcriptional bursting. This model not only reveals fundamental principles of E-P communication in transcriptional bursting kinetics (e.g., burst size and frequency follow their own power-law behaviors) but also provides a general modeling framework toward the 4D nucleome project.

## Introduction

Gene transcription that is tightly related to three-dimensional (3D) genomic organization is a highly complex and regulated process that exhibits a discontinuous episodic bursting behavior (1–8). As two cardinal regulatory elements, the promoter and the enhancer are responsible for the accurate spatiotemporal gene expression ensuring reliable cell functioning and cellular decision-making (9–14). Many experimental studies have been invested in understanding the roles of distal enhancers in regulating transcriptional bursting kinetics (15–19). However, the mechanism of how 3D chromatin organization (in particular 3D enhancer-promoter (E-P) communication) shapes transcriptional bursting patterns still remains elusive.

Hierarchic genomic structures captured by chromosome conformation capture and fluorescence *in situ* hybridization (FISH) as well as other experimental technologies have provided evidences for supporting various possible E-P topologies and linking upstream distinctive E-P communications to downstream gene transcription (20–28). Measurable sustained physical proximity of E-P communication, which is believed to increase the local concentrations of the coactivators and transcription factors (TFs), seems necessary for transcription in living *Drosophila* embryos (18, 29). This proximity is also needed for cells to execute correct gene expression programs, though the ways of E-P communication such as E-P loop and hub hypothesis are still a matter of debate (30, 31). To understand the mechanism of transcriptional bursting theoretically, many models of gene expression have been proposed, including simple models (e.g., the common ON-OFF model) that are mainly based on distinctive transient chromatin behaviors (32–37), and multistate models that seem a good candidate for mimicking complex promoter dynamics in mammalian cells (8, 19, 38–45). However, these models ignored dynamic transcriptional regulation by spatial chromosome topology (46–50). To the best of our knowledge, a comprehensive theory of the information transmission from upstream chromatin organization to downstream transcriptional bursting is still lacking. And important yet fundamental questions such as how E-P communication shapes the observed patterns of mRNA expression and what is the role of the “range of action” for E-P proximity in the control of bursting kinetics remain unsolved. Understanding and revealing transcriptional bursting kinetics characterized by burst size (BS) and burst frequency (BF) require integrative models that consider both 3D chromatin motion (including E-P spatial communication) and upstream-to-downstream regulation.

A collection of experimental evidences has firmly established that the E-P communication strength can significantly raise transcriptional levels (16, 17, 51, 52), e.g., the *sna* distal shadow enhancer generates more bursts than the primary enhancer in *Drosophila* embryos (16). Recent live-imaging measurements have provided the clear evidence that E-P genomic distance can effectively control gene activities and thus affect transcriptional bursting kinetics (16, 53, 54). These experimental observations indicate that the E-P genomic distance arranged by chromosomal rearrangements and the E-P communication strength limited by the E-P spatial distance as well as this distance itself are important parameters impacting the formation of transcriptional bursting patterns. Currently, a new trend is that genomic structures are stochastic at almost every level of organization and this stochasticity is suggestively linked to gene transcription and finally affects transcriptional outcomes (26, 27, 46). Given the important impacts of these factors on transcriptional bursting, an unsolved and even theoretically unexplored issue is what principles govern the formation of transcriptional bursting patterns. How E-P spatial and genomic distances differently influence bursting patterns is unclear, either. As a matter of fact, biological systems are by nature multiscale. To date, many experimental studies have shown distinct timescale differences between upstream chromatin dynamics and downstream bursting kinetics (55, 56). For instance, E-P communication occurs on a timescale of seconds to minutes (18, 29, 57), whereas half-lives of Pol II pause during transcription are on a timescale of minutes to hours (58–60). This temporal disconnection between the upstream and the downstream as well as the stochasticity of genomic organization and transcriptional bursting lies at the heart of a broad challenge in physical biology of forecasting transcriptional outcomes from the dynamics of underlying molecular processes.

Building upon experimental phenomena and acquired data, we develop a general theoretical framework to investigate how E-P communication shapes transcriptional bursting patterns, focusing on the uncovering of underlying molecular processes and dynamical mechanisms. This framework considers 3D upstream chromatin motion on a fast timescale and downstream mRNA bursty production on a slow timescale as well as the connection between the upstream and the downstream. We employ the classical differential Chapman-Kolmogorov Equation (dCKE, (61)) in stochastic process theory to analyze dynamic behaviors of transcriptional bursting across space and time, i.e., 4D transcriptional bursting kinetics (62, 63). Both model analysis and numerical simulations reveal fundamental principles of the E-P communication in modulating transcriptional bursting, e.g., BS and BF obey their own power laws, E-P spatial distance follows Maxwell-Boltzmann distribution instead of previously assumed Gauss distribution, and E-P communication mainly modulates BF but not BS. All these uncovered principles are verified by experimental results and seem universal. We emphasize that our model, which exhibits scalability by interpreting many experimental phenomena reported in the existing literatures (16, 17, 51, 53), provides a general modeling framework for investigating how chromatin dynamics affect transcriptional bursting kinetics in more realistic cases.

## Results

### The First Principle Framework of Gene Transcriptional Bursting

Transcription involving multiple regulatory elements is driven mainly by E-P communication (Fig. 1A), often exhibiting a burst-like pattern (Fig. 1B). Our goal is to develop an analysis framework that can reveal the essential mechanism of transcriptional bursting and predict possible dynamic behaviors compatible with experimental observations. This framework considers that upstream chromatin dynamics play on a fast timescale and downstream transcriptional bursting operates on a relatively slow timescale, but keeps the upstream and the downstream linked via a biologically reasonable way (Fig. 1D).

**Fig. 1.**
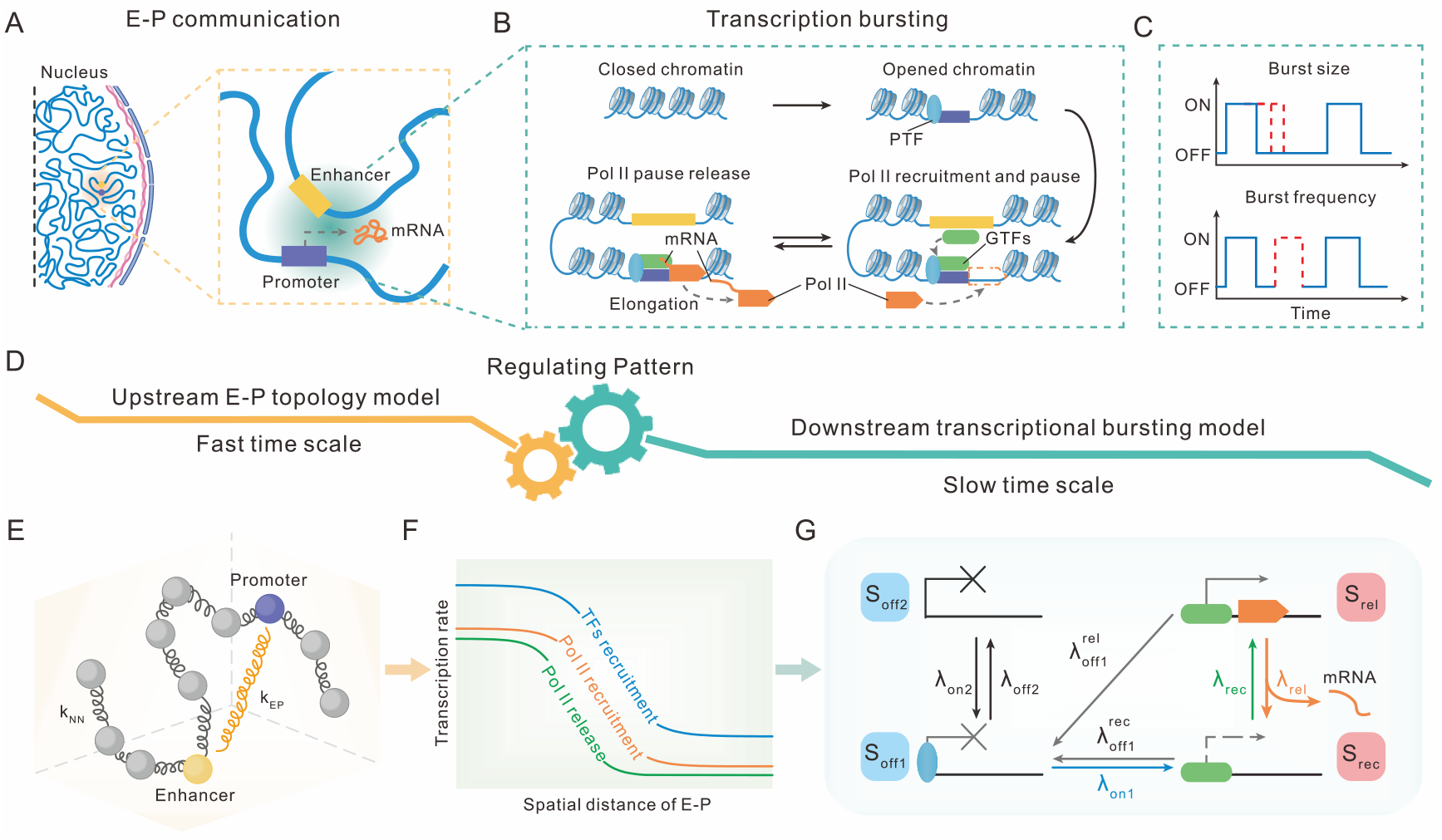
A framework for studying how upstream E-P communication shapes downstream mRNA bursting patterns. (*A*) E-P communication plays a key role in regulating transcriptional bursting in cell nucleus. (*B*) The downstream transcription cycle is a multistep process guided by upstream E-P communication, where only the main steps are depicted. (*C*) Burst size and burst frequency as two quantitative indices to measure the entire bursting system. (*D*) An information flow viewpoint for bridging the temporal disconnection between upstream E-P topology and downstream transcriptional bursting. (*E*) A generalized Rouse model with additional E-P communication (indicated by the yellow spring with coefficient *k*_EP_) for simulating chromatin dynamics. (*F*) A schematic for input-output functions, where the upstream E-P spatial distance is taken as input whereas gene state switching rates as outputs. (*G*) A four-state model for imitating transcriptional bursting. Here *λ*_on 2_ and *λ*_off 2_ are switching rates between a deep inactive state (S_off 2_) and a primed but inactive state (S_off1_); 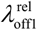 is a transition rate from S_rel_ (representing Pol II pause release state) to S_off1_; 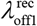 and *λ*_on1_ are transition rates from S_rec_ (representing Pol II recruitment state) to S_off1_ and from S_off1_ to S_rec_, respectively; *λ*_rel_ and *λ*_rec_ are switching rates between S_rec_ and S_rel_.

Our modeling framework is a general dCKE (61), whose form is as follows (see *SI Appendix* B1 for derivation)

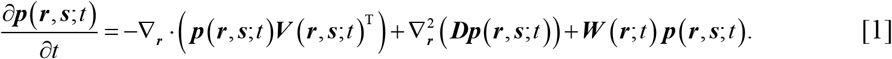

Here the column vector ***p***(***r***, ***s***;*t*) represents the joint probabilities that the positions of *N* nucleosomes are ***r*** and the gene states are ***s*** at time *t*; ∇_*r*_ and 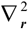 are the gradient operator and the Laplace operator respectively; ***V***(***r***, ***s***; *t*)^T^ represents the deterministic velocity field of chromatin dynamics with T standing for transpose; ***D*** is a diffusion matrix determining the additional noise strength of chromatin motion under isotropic diffusion with diffusion coefficient *D*; and ***W***(***r***;*t*) is the nucleosome position-dependent transition matrix for all gene states. The first term on the right-hand side of Eq. [1] represents a deterministic component modeling upstream chromatin motion and the second term represents a stochastic component accounting for random fluctuations, both altogether describing the upstream chromatin’s spatiotemporal diffusion process. The last term captures downstream gene states’ randomly switching process driven by molecular events such as chromatin remodeling, TFs binding, and Pol II cluster. We point out that Eq. [1] is the first theoretical model that comprehensively considers upstream and downstream molecular processes and the connection between the upstream and the downstream, and can be taken as a good starting point for analyzing how chromatin motion affects transcriptional bursting in complex cases.

From a physical viewpoint, upstream chromatin can be modeled as a polymer discretized into a collection of successive monomers connected by harmonic spring (64–67). Then, chromatin motion evolves according to the overdamped equation 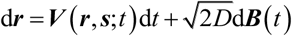, where ***B***(*t*) is a vector of independent Brownian motions (see *SI Appendix* C1). We assume that changes in gene state do not affect chromatin motion although transcriptional outcomes may impact chromatin configurations (68, 69). Therefore, ***V***(***r***, ***s***; *t*) can be approximated as ***V***(***r***; *t*) = −∇_*r*_*U*(***r***;*t*)/*γ*, where *U*(***r***; *t*) is the total potential of chromatin conformation and *γ* is a friction coefficient. Note that *U*(***r***;*t*) can be decomposed as *U*(***r***;*t*) = *U*_NN_ (***r***;*t*) + *U*_EP_ (***r***;*t*), where the potential *U*_NN_ (***r***; *t*) for the chain connection of *N* monomers is set as 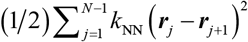 with *k*_NN_ being the spring coefficient and the potential *U*_EP_ (***r***; *t*) for the E-P communication is set as (1/2)*k*_EP_ (***r***_E_ – ***r***_P_)^2^ with E, P ∈{1, ⋯, *N*} representing the enhancer and the promoter respectively. Here *k*_EP_ represents the E-P communication strength and will be taken as a key parameter of our model (Fig. 1E). It should be pointed out that the above model is simplified since realistic molecular processes underlying chromatin motion would be complex, e.g., multiple promoters may interact with one enhancer (70), and chromatin’s spatiotemporal diffusion may be anomalous (71, 72). However, the fact that model predictions are in good agreement with experimental measurements indicates the reasonability of our model assumptions (*SI Appendix*, Fig. S10B).

Downstream transcription apparatus can be characterized by promoter-state switching (Fig. 1B). In line with previous propositions (52, 73), we introduce a four-state regulatory architecture with each state transition corresponding to a biochemical process (Fig. 1G, see *SI Appendix* C2). Specifically, a deep inactive state (S_off2_) and a primed but inactive state (S_off1_), integrated as the OFF state, are used to explain chromatin remodeling and TF binding. Pol II recruitment state (S_rec_, (74)) and Pol II pause release state (S_rel_, (75)), two critical processes involved in transcriptional bursting, are taken as the ON state, and mRNA generation is accompanied by the transition from S_rel_ to S_rec_. If states S_rel_ and S_rec_ switch multiple times during one ON period, transcription will occur in a burst-like manner. Switching rates between these states are depicted in Fig. 1G and the elements of state transition matrix ***W*** in Eq. [1] are related to gene state switching rates (see *SI Appendix* C2).

After identifying the above formulations and settings, we build a link for the information flow from upstream chromatin conformations to downstream biochemical reactions (Fig. 1F). E-P communication carries the upstream information to orchestrate bursting patterns, and downstream transition rates change accordingly. We introduce input-output functions in which E-P spatial distance *d*_S_ is taken as input and *d*_S_-dependent gene state switching rates are treated as outputs, thus bridging the upstream and the downstream (see *SI Appendix* C3). Although how E-P communication is implemented is arguing (30, 31), here we adopt a hub hypothesis that each of switching rates *λ*_on1_, *λ*_rec_ and *λ*_rel_ is a piecewise Hill-like function of *d*_S_, i.e. *λ*_S_(*d*_S_), *c* ∈ {on1,rec,rel} (Fig. 1F). This assumption would be idealistic compared to elaborate organisms, but all the above settings constitute an analysis framework that can be used in the study of complex transcriptional regulation.

It is worth mentioning that we have developed both an effective analytical approach and a numerical simulation algorithm for solving the above entire system (*SI Appendix* B2 gives details of the algorithm implementation and *SI Appendix* E presents analytical details). These methods provide a systematic approach for tracing the respective contributions of the system’s key parameters (e.g., E-P communication strength, E-P genomic distances) to the experimentally observable patterns of transcriptional bursting and further for revealing the essential mechanism of bursting kinetics.

### Distribution characteristics of transcriptional bursting

BS (defined as the number of mRNA molecules produced per burst) and BF (defined as the number of bursts occurred per unit time) are two important quantities characterizing bursting kinetics but are also two random variables (Fig. 1C). In order to reveal the characteristics of BS and BF, we try to find the BS distribution *p_BS_*(*m*) and the cycle time (CT) distribution *p_CT_*(*t*), where CT is defined as the total time that the gene dwells at OFF and ON states, and BF as the reciprocal of CT. In the following, *p_BS_*(*m*) and *p_CT_*(*t*) are denoted uniformly by *p_X_* (*x*), where *X* ∈ {*BS, CT*}.

Note that the entire transcriptional system involves two kinds of timescales – the fast motion of upstream chromatin conformation and the slow occurrence of downstream transcription reactions. To derive distribution *p_X_* (*x*) directly from this multiscale system is a particularly challenge task. Therefore, we resort to a timescale separation method that corresponds to the two limits in which the upstream chromatin motion is either much faster or much slower than the downstream transcription reactions. This method has been successfully used in the analysis of gene regulatory networks (76, 77) and the mutual information that disentangles interactions from changing environments (78).

Specifically, we introduce a scaling parameter *ω*, defined as the ratio of the minimum variable transition rate and the maximum chromatin motion velocity, to represent the maximum achievable ratio of upstream and downstream timescales. If the timescale of the upstream motion is much faster than that of the downstream reactions (*ω* ≪ 1), the quasi-stationary upstream can be well separated from the downstream transcription. Then, *p_X_* (*x*) can be analytically calculated through 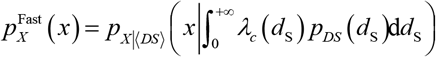 (*SI Appendix*, Fig. S12C), where *p_DS_* (*d*_S_) is upstream E-P spatial distance distribution and 〈*DS*〉 is the expectation of E-P spatial distance. Oppositely, if the timescale of the upstream motion is much slower than that of the downstream reactions (*ω*>> 1), then according to total probability principle, *p_X_* (*x*) can be calculated through 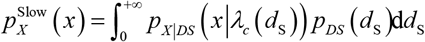 (*SI Appendix*, Fig. S12E). For the intermediate range of *ω* where there is no timescale separation, we find that *p_X_* (*x*) can well fit with the following interpolation formula (again see *SI Appendix* E1)

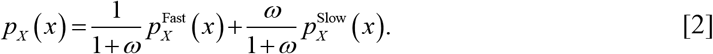

It should be pointed out that this interpolation formula needs to know E-P spatial distance distribution *p_DS_*(*d*_S_) and conditional distribution *p*_*X*|*DS*_ (*x*|*λ_c_*(*d*_S_)) given E-P spatial distance *d*_S_.

By complex calculations, we find that E-P spatial distance *d*_S_ obeys the following exact Maxwell-Boltzmann distribution (see *SI Appendix* E2, Fig. S10A) but not previously assumed Gauss distribution (18)

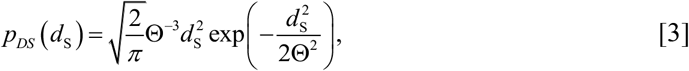

where 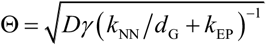, a compound parameter depending on the genomic property, can determine how changes to any of the concerned parameters such as E-P communication strength *k*_EP_ or E-P genomic distance *d*_G_ (different from E-P spatial distance *d*_S_) alter the shape of the *d*_S_ distribution (*SI Appendix*, Fig. S10C). Note that the sum *k*_NN_/*d*_G_ + *k*_EP_ can represent the integrative effect of two paralleling springs, possibly hinting the principle of engineering communication between regulatory elements. Quantitative measurements of the distance between Sox2 and its essential enhancer in living mouse embryonic stem cells have indicated the validity of this Maxwell-Boltzmann distribution (see *SI Appendix* J, Fig. S10B, (79)). It is worth pointing out that Eq. [3] also uncovers a power-law behavior between encounter probabilities and E-P spatial (or genomic) distance, which is in accordance with experimental data for consecutive and nonconsecutive TAD borders in *Drosophila* (see *SI Appendix* J, Fig. S10E, (80)). In addition, the combination of *k*_NN_ equaling 0.1 and *d*_G_ taking 50 shows that the mean E-P spatial distance is also consistent with that of experimental data (see *SI Appendix* J).

Note that *p_X|DS_*(*x*|*λ_c_*(*d*_S_)) is merely related to given transition rates and can be then given analytically (see *SI Appendix* E3). Furthermore, we can analytically show that the BS follows a geometric distribution *Geo*(*θ*) where parameter *θ* represents the success probability of burst termination; the duration of OFF period follows a bi-exponential distribution, supporting that gene inactivation to activation is not necessarily a one-step process following an exponential distribution (8, 43, 81, 82); and the CT distribution is expressed as the weighted combination of multiple exponential distributions, which demonstrates a non-origin peak (*SI Appendix*, Fig. S11).

Finally in this subsection, we point out that Eq. [2] provides high-accuracy approximations of downstream transcriptional outcomes (*SI Appendix*, Fig. S13) and is a useful and simple formula for predicting the dynamics of mRNA over a broad range of timescale separation. By the combination of Eq. [2] and Eq. [3], we can further forecast how upstream chromatin random organization impacts downstream transcriptional outcomes.

### Transcriptional bursting kinetics follow power laws

With the above arsenal of theoretical analysis, we further explore how E-P communication strength *k*_EP_ qualitatively affects transcriptional bursting kinetics (mainly, BS and BF). As one of the most common modular organisms, *Drosophila* has been extensively studied in terms of bursting (83, 84). In our analysis, we take the inherent communication strength between distinctive enhancers such as *sna* shadow and *sna* primary enhancer in *Drosophila* (16, 85, 86) as the value of *k*_EP_, but make use of the fact that effective experimental means such as external hormone (87) or heavy metal stimulations (88) can alter *k*_EP_. In order to reveal the distinct effects of different *k*_EP_ on bursting kinetics, we keep the E-P genomic distance fixed (Fig. 2A).

**Fig. 2.**
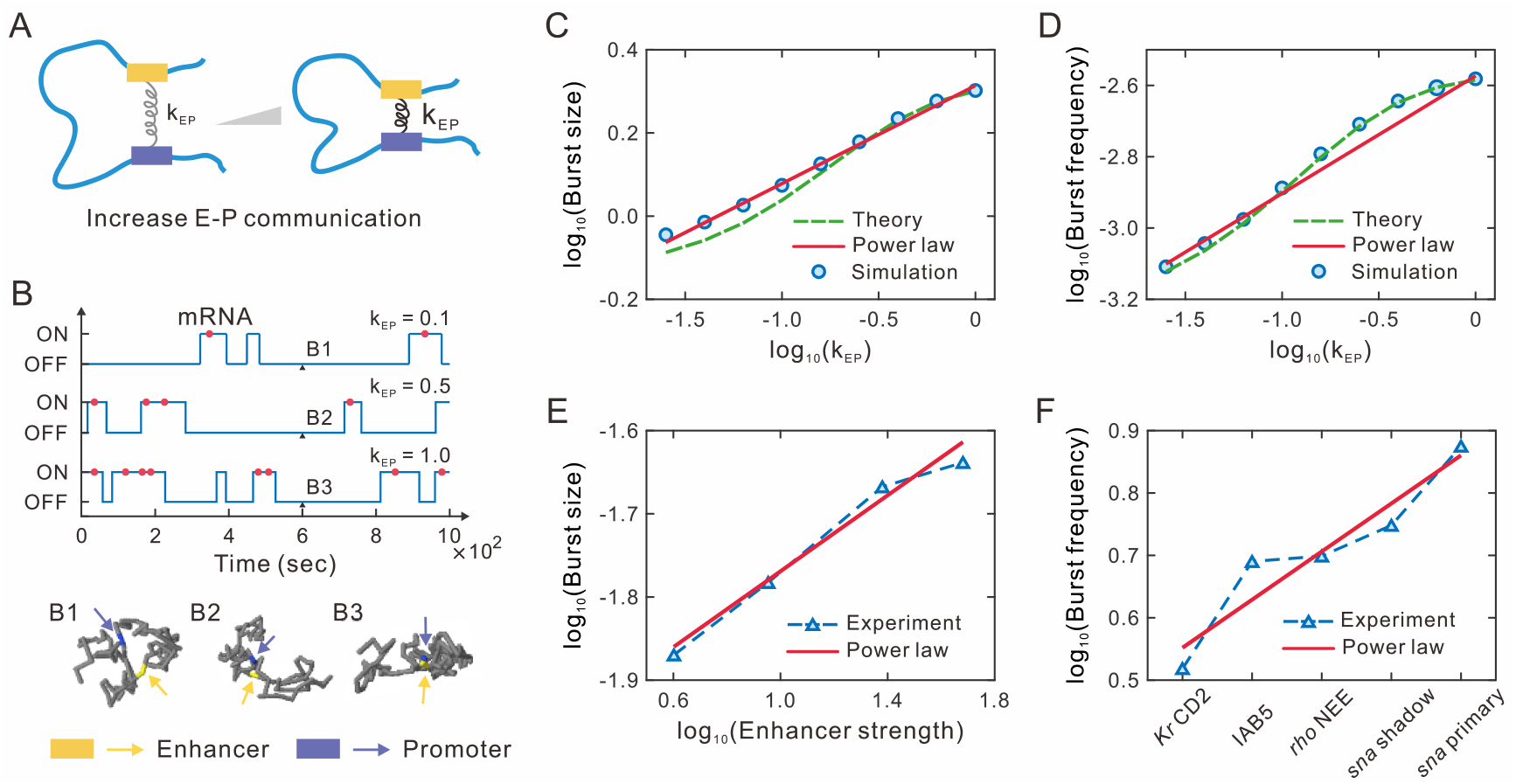
Effect of the E-P communication strength on transcriptional bursting kinetics. (*A*) Schematic illustration of the augment of E-P communication strength *k*_EP_. (*B*) Example traces of binarized transcriptional activity for three different values of *k*_EP_, where red points represent the mRNA generation events and the corresponding 3D chromatin structures are shown on the bottom. (*C*) Log-log plot for the relationship between mean burst size and strength *k*_EP_. The green dashed line represents the theoretical analysis result (Eq. [2]), whereas blue circles show numerical simulations. The red solid line indicates a power-law approximation (Eq. [4]) with scaling exponent *S*_BS_ = 0.23. (*D*) Log-log plot for the relationship between burst frequency and *k*_EP_. Meanings of symbols are the same as those in (*C*), but the scaling exponent is *S*_BF_ = 0.33. (*E*) Log-log plot of experimental data from (17). (*F*) Plot of experimental data from (16). The labels of the *x*-axis represent the *sna* shadow, *sna* primary, *rho* NEE, IAB5, and *Kr* CD2 enhancers, respectively.

Fig. 2B illustrates how changes in *k*_EP_ alter chromatin conformations and further affect bursting profiles. Theoretically, a larger (smaller) *k*_EP_ corresponds to a shorter (longer) E-P spatial distance, and the increasing *k*_EP_ can boost both BS and BF (*SI Appendix*, Figs. S10C, S13A-B). These features have been qualitatively supported by many experimental evidences from different organisms (16, 17, 52, 89). Notably, since ON state dwell-time remains fundamentally the same, the amplified BF can be achieved by progressively reducing OFF state dwell-time (*SI Appendix*, Figs. S13C-D, (19, 89, 90)). This implies that the gene state switching rate *λ*_on1_, associated with TF recruitment in biology, is a principal parameter affecting BF, in agreement with the experimental finding that a higher TF level leads to a higher BF on the c-Fos gene (51).

To reveal the qualitative relevance of bursting kinetics to E-P communication strength, we further analyze two logarithmic gains *∂*log_10_BS/*∂*log_10_ *k*_EP_ and *∂*log_10_BF/*∂*log_10_ *k*_EP_, which measure how BS and BF are affected by *k*_EP_. Binary approximations of gene state switching rates provide a good strategy for simplifying analysis including estimating BS and BF (see *SI Appendix* F1, Figs. S14D-E). Our theoretical analysis shows that the logarithmic gains are about zero when *k*_EP_ is too small (*k*_EP_ < 0.01) or too large (*k_EP_* > 1) (*SI Appendix*, Figs. S14A-C), implying that transcriptional burst response is insensitive to *k*_EP_ in this case. Then we focus on a reasonable biological range of *k*_EP_ (0.01 < *k_EP_* < 1). The logarithmic gains are almost constant and BS and BF can be thus approximated with linear functions of *k*_EP_ in logarithmic scale (Fig. 2C-D, *SI Appendix*, Figs. S14D-E). Concretely, BS and BF approximately obey the following power-law behaviors

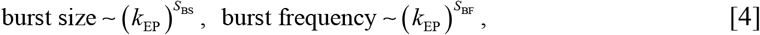

where *S*_BS_ and *S*_BF_ are two positive scaling exponents that can be theoretically estimated (see *SI Appendix* F2). Note that these two different scaling exponents, e.g., *S*_BS_ = 0.23 in Fig. 2C and *S*_BF_ = 0.30 in Fig. 2D, are critical indices since they reflect the ability of the BS and BF responses to E-P communication, which will be in details analyzed in the next section. To ascertain whether the power-law behaviors shown in Eq. [4] always hold, we let key parameters change in broad ranges. For all the possible cases of parameter values, we find that log-log plots between BS (or BF) and *k*_EP_ still demonstrate an approximately linear relationship, implying that the growing tendency can be featured by power-law relationships (*SI Appendix*, Fig. S15).

The above theoretical predictions have been verified by experimental data. For example, the addition of estradiol that triggers stronger contacts between *β*-globin and its locus control region over time leads to the increase of *β*-globin gene transcriptional BS (17). If we use the erythroid maturation time after adding the hormone to reflect the E-P communication strength *k*_EP_, the BS of *β*-globin gene can well fit with a power-law function (Fig. 2E). In another example, transgene experiments examining the effects of differential enhancers on BF showed that a stronger enhancer such as *sna* shadow produces more bursts than a weak enhancer such as *rho* NEE in living *Drosophila* embryos (16). We find that the corresponding BF exhibits a power-law-like behavior if the enhancer’s strength increases in an equidistant interval on a logarithmic scale (Fig. 2F). *SI Appendix* Figs. S16A-C demonstrate more experimental data (16, 51) on the dependences of BS and BF as well as the mRNA expression level (approximately equals to the product of BS and the BF (91)) on E-P communication strengths, further verifying these power-law behaviors.

### E-P communication mainly modulates burst frequency rather than burst size

Eq. [4] indicates that BS and BF regulated by E-P communication strength *k*_EP_ follow their own power-law behaviors. Since *S*_BS_ and *S*_BF_ in Eq. [4] reflect the regulation ability of *k*_EP_, we next compare the sizes of *S*_BS_ and *S*_BF_, i.e., compute the ratio of *S*_BF_/*S*_BS_, to show which of BS and BF is primarily regulated by E-P communication. For this goal, we let E-P communication strength and all gene state switching rates change in broad ranges. In general, extracting insights from directly analyzing the effects of these changes in the high-dimensional parameter space consisting of E-P communication strength and all gene state switching rates on BS and BF is very difficult. Therefore, we resort to a dimension reduction method that maps the high dimensional parameter space into an experimentally measurable and theoretically computable two-dimensional space (Fig. 3A) in which two indices *ρ*_BS_. and *ρ*_BF_ are defined as the ratios of the maximum BS and BF over the minimum BS and BF, respectively.

**Fig. 3.**
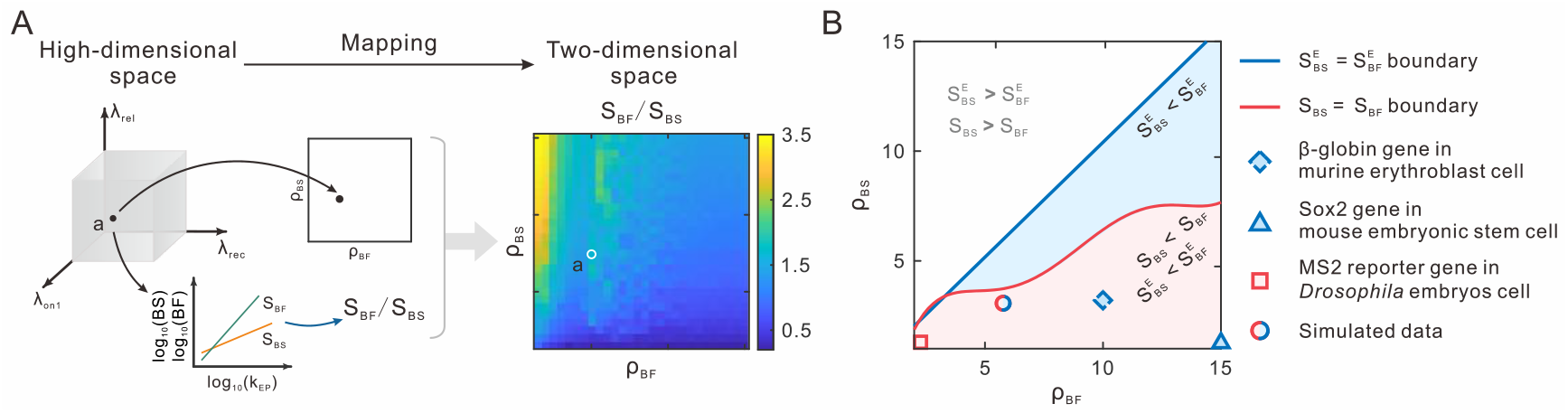
Separatrixes for power-law behaviors of BS and BF in the (*ρ*_BF_, *ρ*_BS_) plane. (*A*) Schematic illustration of the dimension reduction method, where a high-dimensional parameter space is mapped into the (*ρ*_BF_, *ρ*_BS_) space (from the parameter coordinate space to the square) and the calculated values of *S*_BF_/*S*_BS_ (the bottom inset) are plotted in the (*ρ*_BF_, *ρ*_BS_) plane (the heatmap). (*B*) The red line stands for the boundary obtained by smoothing the theoretical analysis, the blue line for the boundary after enhancer deletion. The colored regions indicate that E-P communication affects BF more than BS under different situations. The square at point (2.3,1.3) corresponds to the data in (16), the triangle at point (15,1.4) to the data in (92), the dashed diamond at point (o,3.3) to the data in (17), and the red blue mixed circle to the simulated data in this paper. Note that the BF in (92) defined as the reciprocal of OFF state dwell-time, which is approximately converted to the reciprocal of CT, and that the *ρ*_BF_ at the dashed diamond point related to burst fraction in (17) is approximate. By the blue region, we mean the whole areas below the blue line (some part is covered by red regions).

Next, we seek to calculate the value of *S*_BF_/*S*_BS_ in the (*ρ*_BF_, *ρ*_BS_) space. Note that one point in the low dimensional space possibly corresponds to multiple points in the high-dimensional space due to the irreversibility of the mapping. To overcome this multiple-to-one difficulty, We use the averaging method to estimate ratio *S*_BF_/*S*_BS_, finding that this ratio generally increases with the growth of *ρ*_BF_ or with the shrinking of *ρ*_BS_ (Fig. 3A, *SI Appendix*, Fig. S17A). In Fig. 3B, the separatrix *S*_BF_/*S*_BS_ = 1 (red line) separates the space into two regions: upper *S*_BF_ > *S*_BS_ and lower *S*_BF_ < *S*_BS_. In particular, the data shown in Figs. 2C and 2D correspond to the point (*ρ*_BF_, *p*_BS_) = (5.8,3.1), which apparently locates in the area of *S*_BS_ < *S*_BF_ (*S*_BF_/*S*_BS_ = 1.4, Fig. 3B, red-blue mixed circle). To show the reliability of this averaging method, we use the minimum (or maximum) value rather than the mean to compute *S*_BF_/*S*_BS_. As a result, the separatrix *S*_BF_/*S*_BS_ = 1 displays a similar change trend, i.e., the range of the *ρ*_BS_ satisfying *S*_BS_ < *S*_BF_ gradually becomes larger with the increase of *ρ*_BF_ (*SI Appendix*, Figs. S17A-B).

To complete the above analysis, we consider enhancer deletion where *k*_EP_ is exactly zero but out of the range 0.01 < *k*_EP_ < 1 (*SI Appendix*, Fig. S11A). For the same parameter values as in the above calculation, we find that the sizes of *ρ*_BS_ and *ρ*_BF_ with enhancer deletion are the same as those with enhancer regulation but the slope ratio 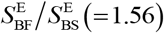 (representing the ratio after enhancer deletion, see *SI Appendix* G) is different from the value of *S*_BF_/*S*_BS_ (= 1.4) (Fig. 3B, red-blue mixed circle). Using the results obtained by a similar method of calculating *S*_BF_/*S*_BS_, we draw the separatrix 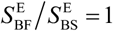 in Fig. 3B (blue line). Then we find that no matter whether enhancer deletion is considered, this does not influence *ρ*_BS_ and *ρ*_BF_, but impacts ratios *S*_BF_/*S*_BS_ and 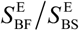 (referring to two different oblique lines in *SI Appendix*, Fig. S11F).

Finally, we locate the experimental data into the (*ρ*_BF_, *ρ*_BS_) space and determine which of BS and BF is mainly regulated by E-P communication. Although experimental data on gene state switching rates lack, we may use the maximum and minimum BS and BF experimentally available to compute *ρ*_BS_ and *ρ*_BF_. Based on analysis of data from transgene experiments, we find that the MS2 reporter gene displays different bursting kinetics for different enhancer strengths, e.g., for a stronger *sna* shadow enhancer and a weaker *Kr* CD2 enhancer, the result is (*ρ*_BF_, *ρ*_BS_) = (2.3,1.3) (Fig. 3B, red square point, (16)). The maximum BS of the gene *β*-globin measured in an experiment of erythroid maturation is over three times the minimum gained by the deletion of the LCR enhancer (Fig. 3B, dashed blue diamond point, (17)). In addition, based on the Sox2 gene expression data with and without enhancer deletion in normal ES cells, we can infer that the BS reduces 1.4 times and the BF reduces roughly 15 times (Fig. 3B, blue triangle point, (92)). Interestingly, all these analyzed experimental data are all located in a narrower region of *S*_BS_ < *S*_BF_ (the right lower region in Fig. 3B), further verifying that the E-P communication mainly modulates BF rather than BS.

### Threshold phenomena in modulation of bursting kinetics by E-P genomic distance

Having devled into the qualitative effect of E-P communication strength *k*_EP_ on bursting kinetics as described in Eq. [4], we next analyze how E-P genomic distance *d*_G_ quantitatively impacts transcriptional activities and further bursting kinetics. To focus on the effect of *d*_G_ on BS and BF, we freeze *k*_EP_ but let *d*_G_ increase. This can be achieved by arranging the enhancer and the promoter symmetrically on the chain and letting them move in the opposite direction along the chain (Fig. 4A).

**Fig. 4.**
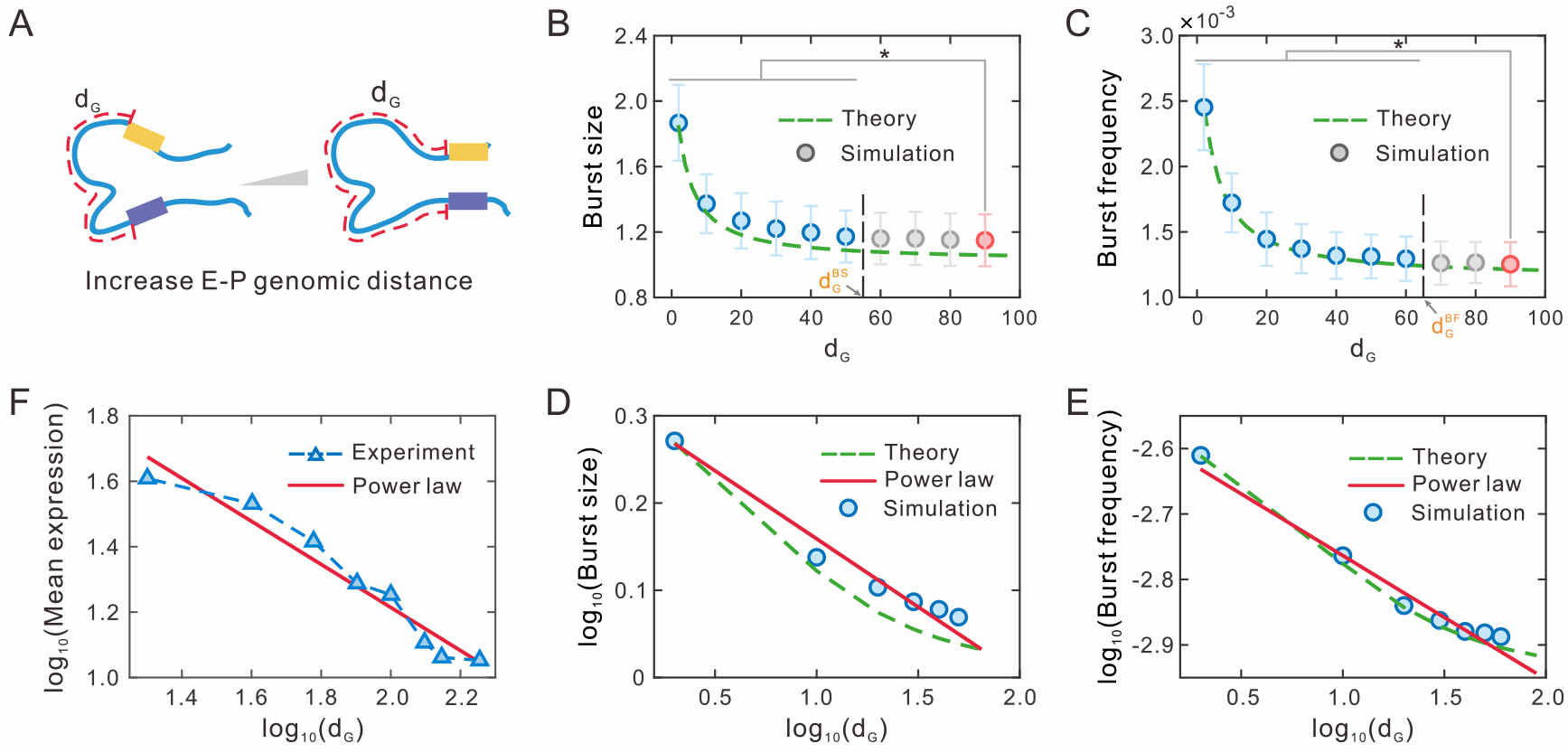
Effect of E-P genomic distance on transcriptional bursting kinetics. (*A*) A schematic diagram shows the increment of E-P genomic distance *d*_G_. (*B*) Influence of *d*_G_ on burst size. Dashed green line represents the theoretical result (Eq. [2]) whereas circles show numerical simulations. The blue circles indicate significant differences compared to the end data point (red) via Kruskal-Wallis nonparametric test, while the gray circles indicate no significance. (*C*) Influence of *d*_G_ on burst frequency, where the meanings of all symbols are the same as those in (*B*). (D) Log-log plot shows the relationship between mean burst size and *d*_G_. Blue circles are the data points with significant differences in (*B*). The red solid line indicates the power-law approximation (Eq. [5]) with the scaling exponent *S*_BS_ = −0.16. Meanings of the symbols are also similar to those in (*B*). (*E*) Log-log plot shows the relationship between burst frequency and *d*_G_. Meanings of the symbols are similar to those in (*D*) but the scaling exponent is *S*_BF_ =–0.19. (*F*) Log-log plot of experimental data from (53).

From Eq. [3], we see that *d*_G_ and *k*_EP_ have the just opposite impact on E-P spatial distance (*SI Appendix*, Fig. S10C). By the similar method of analyzing the effect of *k*_EP_ on BS and BF, we find that two logarithmic gains *∂*log_10_BS/*∂*log_10_ *d*_G_ and *∂*log_10_BF/*∂*log_10_ *d*_G_ are negative, implying that the increasing E-P genomic distance can reduce BS and BF (*SI Appendix*, Figs. S14F-H). With the further increase of *d*_G_, both BS and BF finally reduce to constants. These analyses indicate that the responses of BS and BF to the E-P genomic distance exhibit biphasic (first descent and then stable) change tendencies if other system parameters are fixed (Figs. 4B and 4C).

In order to confirm the above biphasic behaviors obtained by theoretical analysis, we first carry out numerical simulations and then perform the Kruskal-Wallis nonparametric test for simulated samples. This statistical test can identify the turning point of the biphasic responses of BS and BF to *d*_G_. Here the turning point (referring to black dashed lines in Figs. 4B and 4C) is defined as the value of *d*_G_ for which the Kruskal-Wallis test has no significant difference for the first time (referring to grey circles in Figs. 4B and 4C). We denote by 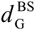 and 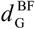 the *d*_G_ values corresponding to the turning points of BS and BF respectively. Then, we observe from Figs. 4B and 4C that 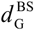 is smaller than 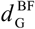. This difference highlights the importance of BF in the E-P communication-based regulation of transcriptional bursting kinetics.

Furthermore, we seek to use functions to characterize the above biphasic behaviors. Note that the logarithmic gains of BS and BF are almost constant for *d*_G_ in the first phase (i.e., 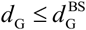 or 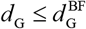), implying that in this phase, the logarithmic gains can be approximated as linear functions of *d*_G_ or equivalently, BS and BF can be approximated as power functions of *d*_G_ with exponents *S*_BS_ and *S*_BF_ respectively (see *SI Appendix* F2, Figs. S14I-J). For the second phase (i.e., 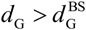 or 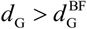), BS and BF are all constants. In summary, the change tendencies of BS and BF can be mathematically described as

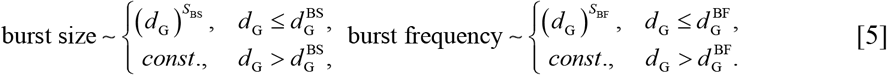

Here *S*_BS_ and *S*_BF_ are two negative scaling exponents that can be theoretically estimated. Note that these scaling exponents are in general different (e.g., *S*_BS_ = −0.16 in Fig. 4D, red) and BF (e.g., *S*_BF_ = −0.19 in Fig. 4E, red). An intuitive explanation for the asymptotic trend in large *d*_G_ is that the lumping parameter 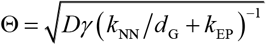 in Eq. [3] approaches to a constant with the enlargement of *d*_G_, indicating that for sufficiently large *d*_G_, the effect of E-P communication on transcriptional bursting is mainly determined by communication strength *k*_EP_ rather than by E-P genomic distance *d*_G_.

The above biphasic behaviors have been also verified by experimental results. First, quantitative living-imaging methods showed that large E-P genomic distance between *sna* shadow enhancer and its target gene significantly diminishes the levels of BS and BF in *Drosophila* (16, 54), in accordance with our prediction on what BS and BF are monotonically decreasing with increasing *d*_G_. Second, the mRNA expression level, which approximately equals to the product of BS and the BF (91), is an approximately linear function of smaller E-P genomic distance *d*_G_ in logarithmic scale (53) (Fig. 4F). Third, now that a monotonically decreasing transcription level does not drop to zero, the case that the level remains at a low level for large *d*_G_ could happen (53). In a word, the E-P genomic distance impacts the mRNA expression level in a biphasic and piecewise power manner as described in Eq. [5] (53).

### Mutual information reveals the dependence of bursting kinetics on E-P communication

In the above sections, we have used the average method, i.e., calculating the mean BS and BF (the reciprocal of mean CT) that only considers the mean information of BS’s and CT’s distribution, to study the impact of E-P communication on bursting kinetics. Here, we investigate bursting kinetics from a perspective of information transmission. The signaling pathway of transcriptional regulation transmits upstream E-P communication signal distribution *p_DS_*(*d*_S_) to downstream transcriptional output distribution *p_X_*(*x*) (Fig. 5A, *SI Appendix*, Fig. S13). The noisy “promoter channel” limits the fidelity of this information transduction. To better understand the ability of E-P communication to regulate BS and BF, here we use Shannon mutual information to measure how much information about the E-P communication is encoded in bursting kinetics (93, 94). Mutual information MI (*X, DS*) measured in bits is defined as

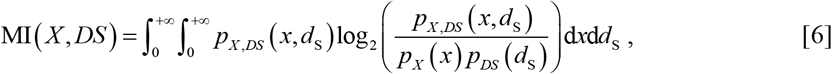

where *p_X_*(*x*) with *X* ∈ {*BS, CT, OFF, ON*} represent BS distribution, CT distribution, OFF and ON dwell-time distributions, respectively (*SI Appendix*, Fig. S13). The mutual information defined in such a manner can capture the contributions of all the aspects of transcriptional output distributions, not just the mean values.

**Fig. 5.**
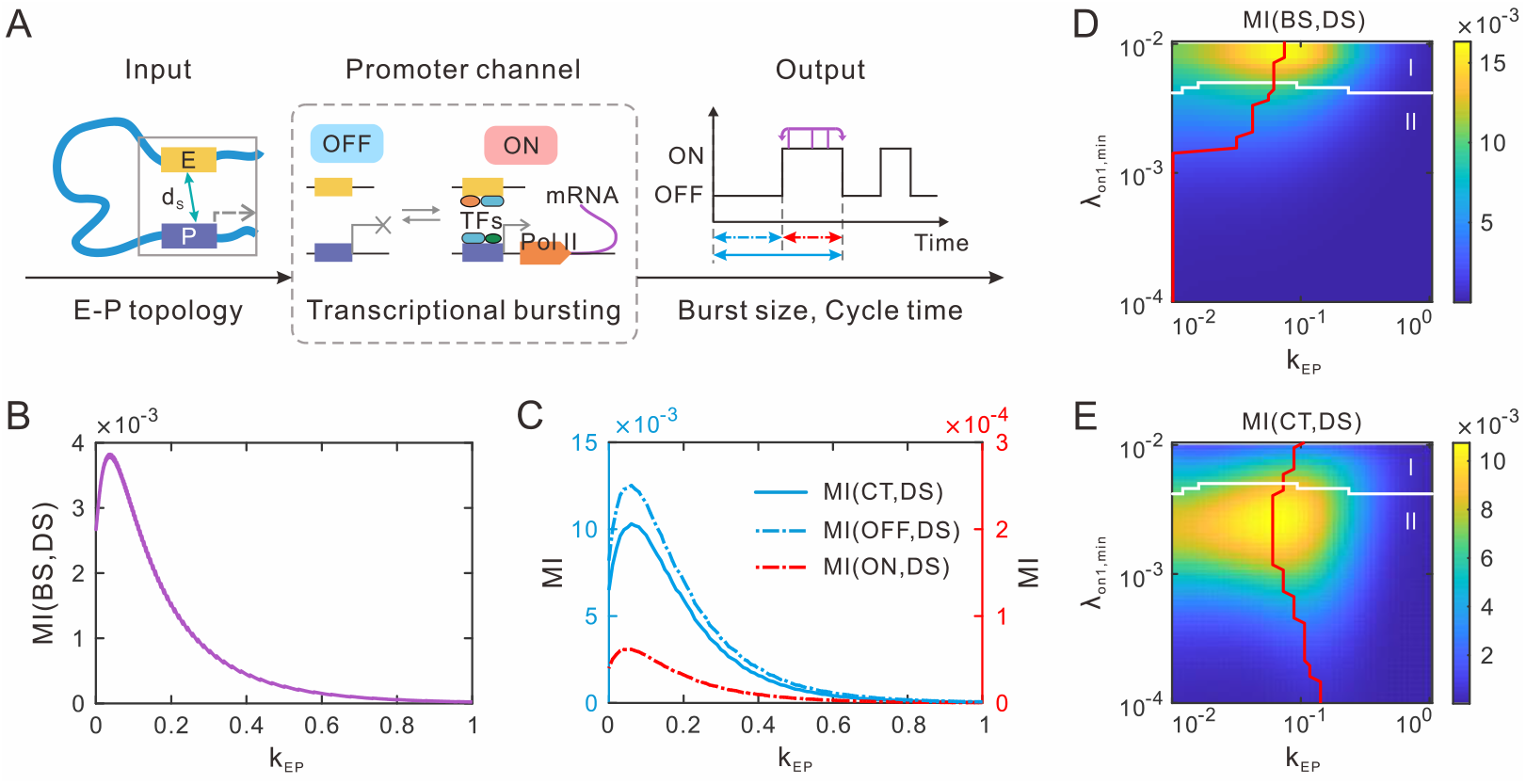
Mutual information reveals the effect of E-P communication on bursting kinetics. (*A*) An information theoretic framework used to study the effect of input E-P topology to bursting output. The promoter governing transcriptional bursting can be considered a noisy channel. (*B*) Mutual information between E-P spatial distance and burst size (MI(*DS*, *BS*)) as a function of E-P communication strength *k*_EP_. (*C*) Mutual information between E-P spatial distance and cycle time (or OFF time, ON time) as a function of *k*_EP_. (*D-E*) Heatmap shows the effect of *k*_EP_ and minimum rate *λ*_on1,min_ on MI(*DS*, *BS*) (*D*) and on MI(*DS*, *CT*) (*E*). The red line shows the values of *k*_EP_ obtained from the MMIs with different *λ*_on1,min_. The white line is the separatrix between the values of MI(*DS*, *BS*) and MI(*DS*, *CT*). Region I is the region of MI(*DS*, *BS*)>MI(*DS*, *CT*) and region II is the region of MI(*DS*, *BS*) < MI(*DS*, *CT*).

Based on the computation of MI(*X*, *DS*) in a wide range of *k*_EP_, we find that MI(*BS*, *DS*) is much smaller than MI(*CT*, *DS*) (Figs. 5B and 5C), implying that BF transmits more information than BS. Moreover, MI (*ON*, *DS*) is two-order smaller than MI (*OFF, DS*), indicating that ON dwell-time is insensitive when responding to the transduced information (Fig. 5C, red dashed line). Additionally, all the MI(*X*, *DS*) in Figs. 5B and 5C exhibit first an increase and then a decrease, indicating that the information transduction capacity is tunable and there exists an optimal *k*_EP_ that maximizes the MI (*X*, *DS*).

Note that the maximum mutual information (MMI) measures the maximum information transmission capacity over all possible signal distributions. Obviously, the MMI exists in Figs. 5B and 5C, where *k*_EP_ = 0.05 for MI(*BS*, *DS*) and *k*_EP_ = 0.07 for MI(*CT*, *DS*). To ascertain whether or not such an optimal *k*_EP_ always exists, we let key parameters change in wide ranges and find that there is indeed a critical *k*_EP_ maximizing the MI(*X*, *DS*) shown in Figs. 5D and 5E and *SI Appendix*, Fig. S18. In addition, the MMIs for BS and CT correspond to different *k*_EP_ values (see red lines in Figs. 5D and 5E). This asynchronous maximization indicates that the signal transduction pathway is BS- or BF-specific. In particular, the observation that MI(*DS*, *BS*) is smaller than MI(*DS,CT*) in region II (Fig. 5D and 5E, the region below the white line) indicates that BF transmits more E-P communication information than BS, and consequently, E-P communication mainly regulates BF rather than BS.

## Discussion

Imaging studies, high-resolution chromatin conformation maps, and genome-wide occupancy profile of architectural proteins have revealed that genome topology encoding E-P communication information is tightly correlated with gene expression through complex regulatory layers, creating various possible transcriptional burst phenotypes. The prediction of these phenotypes requires a quantitative understanding of transcriptional bursting mechanisms underlying the influence of genomic architectures. In this paper, we have proposed a multiscale model, which integrates 3D the information on chromatin dynamics into transcription bursting, to shed light on the pivotal role of E-P spatial communication in the control of transcriptional bursting kinetics.

We have established a general modeling framework that captures salient features of intra-nuclear transcriptional bursting processes. First, the generalized Rouse model, which can well describe chromatin motion, including E-P communication, was used to model chromatin dynamics (64–67). Second, the four-state model of gene transcription adopted here captured important events occurring in transcriptional processes. Two ON states in our model can describe not only the production of nascent mRNA accompanied by the change of promoter states but also the traveling ratio property reflecting the Pol II pause release rate (see *SI Appendix* E3, Fig. S18), two essential characteristics distinguishing from previous models. Third, an input-output relation was proposed to link upstream chromatin configurations to downstream gene transcription where the necessity of E-P proximity (18) is well described in our model.

Our analysis has given a clear answer to the question of which of BS and BF is modulated more than the other by E-P communication. In contrast to the transcriptome-wide inference that showed the important role of E-P communication in controlling BF (92, 95, 96), we have theoretically confirmed that enhancer mainly modulates BF rather than BS (Fig. 3B), which is in agreement with observations of most experiments (16, 17, 53, 90, 97, 98). Moreover, we have given a theoretical estimate on the effect of the enhancer on BF and BS, which is based on directly calculating the ratio of the maximum and minimum values of BS and BF experimental available. In addition, from the view of mutual information, we have shown that transcriptional bursting kinetics are regulated by transmitting more information to BF, which further verifies the importance of enhancers in modulating BF.

Our results allow us to make important predictions about how upstream chromatin dynamics affect downstream transcriptional bursting kinetics. First, we have shown that E-P communication strength *k*_EP_, a key parameter in our model, modulates BS and BF in power-law manners. This power-law behavior was also verified by experimental data (16, 17, 51). Second, we have demonstrated that E-P genomic distance *d*_G_, another vital parameter in our model, modulates BS and BF in biphasic fashions. This biphasic phenomenon was also verified by recent experimental observations (53). The opposite change trends for the effects of *k*_EP_ and *d*_G_ on bursting kinetics showed that the combination of *k*_EP_ and *d*_G_ would be a flexible regulation strategy that can provide insights into complex mechanisms of biological processes in organisms. We have shown that different gene state transition rates lead to different scaling exponents in power-law behaviors of BS and BF, indicating that the regulatory logics shaping differential signal transduction pathways are cell-specific and even gene-specific (99). Third, using mutual information to measure how upstream E-P communication dynamically and stochastically regulates downstream transcriptional bursting kinetics would be a more reasonable way. We have shown that the promoter information transduction capacity is tunable, and the MMI can be obtained by adjusting E-P communication strength *k*_EP_.

Our results are not limited to specific genes in *Drosophila* but may be applied to other organisms. The experimental data from different organisms such as a mouse model can be also well fitted (Figs. 3E-F and 4F) (17, 53), indicating the extensibility of our results. Importantly, our modeling framework can also be extended to more complex situations. For example, some experimental studies reported that transcription can affect chromatin structure (68, 69), and we can incorporate this feedback to our model by modifying the gene-state-dependent drift function ***V***(***r***, ***s***; *t*) in Eq. [1]. In addition, our modeling of chromatin motion is not limited to one pair E-P communication. The potential applications may include the cases of multiple enhancers to one promoter (70), one enhancer to multiple promoters (16), or super-enhancers (100). Finally, using multi-state models of gene expression to extract insights from enormous experimental data and complex biological phenomena is impressing. Our four-state model is not the default option, and we may adjust the form of the downstream transcription model to include more complex biological processes such as mRNA splices (101) and cell cycle (102).

Finally, we point out that our theoretical model, which aims to develop a general modeling framework to study 4D transcriptional bursting kinetics, may provide an opportunity for a dialogue between theoretical studies and biological experiments. We envision that our modeling framework will be useful for biophysical analysis of broader in *vivo* cellular processes.

## Acknowledgments

This work was supported by grants 12171494, 11931019, and 11775314 from Natural Science Foundation of P. R. China, and by grants 019B110233002, 202007030004 from Key-Area Research and Development Program of Guangzhou, P.R. China.

## Supplementary Information

### A Introduction to background knowledge

#### A1 Three-dimensional genome organization and dynamics

##### 1. Multiscale spatial organization of genome

Genomes have complex three-dimensional (3D) architectures employed to store complete genetic information in an organism. To accommodate the genome within a confined space of cell nucleus, the genomes must be organized and compacted, with the assurance that genomic function is accurately achieved at any location and time. This multi-level 3D genomic organization is highly ordered but spatially multiscale (1, 2).

In spite of their architectures depending on organisms and with differently characteristic scales, genomes contain DNA, nucleosomes, fibers, loops, domains, and compartments as well as chromosomes. DNA is composed of 2 polynucleotide chains wrapped around each other to form a double helix structure. Nucleosomes made up of the octamers that are composed of core-histones are wrapped by DNA and further helices to form chromatin fibers. Chromatin fibers can be folded to form loops that typically link upstream regulatory elements such as enhancers close to the promoters to control downstream gene transcription. Also, chromatin fibers themselves can form topologically associating domains (TADs), which are genomic regions that preferentially interplay with each other. The TADs are assembled to form higher-order chromatin compartments. Chromosomes are the primary elements of genomic organization, and distinct chromosomes exist in the nucleus in the form of chromosomal territories.

##### 2. Stochasticity of genome organization

Genomic organization is highly stochastic within each single cell and this stochasticity is similar to cell-to-cell variability in transcriptional outcomes (3). It has been experimentally observed that chromosomes are highly variable in structure, position, and level of cross-chromosomal contacts between two homologs and even between individual cells (4). Also, there is heterogeneity in the location of TAD boundaries and the formation of chromatin loops (5–7). In particular, the stochasticity of chromatin loops propagates directly to enhancer-promoter (E-P) communication, implying that distal regulation is a source of transcriptional noise. A major challenge is to identify how an enhancer distally regulates transcription via a precise regulation mechanism under this highly stochastic genome organization. It is noted that 3D genomic organization simultaneously possesses high order and high variability properties that are in conflict with each other. This double property is a sign that genomic organization is a self-organization, a mechanism that is universal in nature (1).

#### A2 Enhancer-Promoter communication

The chromatin loop structure of 3D genomes, which serves as an intermediate regulation between regulatory elements, often contributes to enhancer-promoter (E-P) communication and further gene expression (8). This E-P communication is a long-range interaction that makes it possible to spatially contact the promoter of the target gene by the distal regulatory element--enhancer, rather than to be just influenced by the proximal regulatory element. These observations raise an open question: how enhancers and promoters construct regulatory landscapes and how E-P communication is sculpted in 3D chromatin structure. See Section C3 for explanations.

#### A3 Multistep transcription process

Transcription is a complex multistep process, involving chromatin remodeling, recruitment of transcription factors (TFs) and RNA polymerase II (Pol II), transcription initialization, transitions between transcriptionally active and inactive states, etc. Specifically, a transcription period roughly involves the following eight steps: the first step is chromatin opening, the second step is the formation of pre-initiation complex (i.e., PIC formation), the third step is transcription initialization, the fourth step is promoter escape/clearance, the fifth step is the escape from pausing, the sixth step is productive elongation, the seventh step is another cycle of transcription, and the eighth step is transcription termination (9). In Section A5 and C2, we have also given some detailed explanations.

Concretely, E-P communication can regulate almost every step of the expression process involved in transcription initiation, including PIC formation, Pol II activation transcription and Pol II pause release. Therefore, it is essential to consider the multi-step process of transcription in investigating the impact of E-P communication on transcriptional expression. In Section C3, we have given some detailed explanations about how E-P communication regulates transcriptional bursting.

#### A4 Transcriptional bursting

Bursty expression patterns, as the outputs of multi-step transcriptional processes, have been observed in multiple experiments in various systems and cell types (10–14). Many mathematical models used to describe burst phenotypes have been proposed, such as the random telegraph (ON-OFF) model (15) and three-state model (16) as well as the multi-state model (17). In fact, the random telegraph model has been widely adopted to both explain transcriptional dynamics and understand the mechanisms of whole genome transcriptional regulation (18). In addition, three-state or multi-state model have been theoretically used to explain experimental phenomena (19, 20).

Transcriptional bursting kinetics can be characterized by two main quantities: burst size and burst frequency. First, once the burst is initiated, the Pol II is able to load on the promoter that DNA is transcribed into mRNA. When the promoter changes from the active to the inactive states, the number of mRNAs produced in this burst can be calculated, i.e., burst size. Another quantity that determines how frequently a burst can be observed, is burst frequency. It depends on the process that precedes the initial transcription, such as E-P communication, recruitment of TFs. Understanding how genes encode molecular regulatory mechanisms of bursting (burst frequency and size) is a highly interesting matter.

#### A5 Different timescales between chromatin motion and transcriptional bursting

First, chromatin spatial folding and chromatin accessibility are essential for the contact of distal enhancers with their target promoters (21). The basic unit of chromatin is nucleosome and nucleosome rewrapping is the obstacle hindering the access of Pol II (22). The assembly or unwrapping time of nucleosome is about 1 second (1s) (23, 24). Meanwhile, the communication between enhancer and distal promoter can form on a timescale of seconds to minutes (25–27). Therefore, the timescale of chromatin motion is less than seconds. Some studies have used 1e-4 to 1e-2s (28–30) to represent the timescale of chromatin dynamic motion.

Second, transcription will be possible once the chromatin is open, and a cascade of biochemical processes will be triggered, such as TFs binding, Pol II recruitment, transcriptional elongation, etc. The transcription cycle might be more complex than the mentioned-above simple eight steps, and many questions about transcriptional bursting remain elusive. Imaging studies have shown that the TFs binding to DNA is a highly transient and dynamic process, occurring on the order of hundreds of milliseconds to seconds (31–34). TFs binding, especially TFIID, can distort the TATA box (35). This distortion is considered to be a marker of locating the promoter, allowing subsequent PIC assembly steps. Besides, Pol II can form clusters in vivo and its dynamics takes place in a few seconds (36). TFs recruitment and Pol II recruitment are related to E-P communication. Although TFs and Pol II can bind to enhancer and promoter and maintain the E-P communication, the E-P spatial distance is highly dynamic (37) and stochastically fluctuating on a timescale of seconds to minutes (25–27). In addition, Pol II pause is a significant event in bursting processes. Pol II pause release needs positive transcription elongation factor b (P-TEFb) to phosphorylate Ser2 and negative elongation factor (NELF) (9). Recent genome-wide studies have shown that the half-lives of Pol II pause is on a minute timescale (38). Of course, a transcriptional burst may include multiple transcription cycles (i.e., another round of transcription after Pol II pause release). Thus, transcription processes occur in discontinuous episodic bursts and can last minutes to hours (39–41). Meanwhile, multiple cycles and burst duration indicate that transcriptional bursting itself is a multiscale process (16).

#### A6 Question: how E-P communication regulates transcriptional bursting

Imaging studies of chromatin conformations and genome-wide occupancy data of architectural proteins have revealed that genome topology, especially E-P communication, is tightly intertwined with gene expression (42, 43). Here, a challenging question arises: how E-P communication regulates transcriptional bursting. The fact that the genomic structures are stochastic (3) and that transcription process and chromatin movement are on different timescales (44) brings difficulties to address this question. In this paper, we establish an analysis framework to reveal the mechanism of transcriptional bursting and predict possible dynamic behaviors compatible with experimental observations.

### B Differential Chapman-Kolmogorov Equation (dCKE)

#### B1 Derivation of dCKE

As mentioned in Section A, gene transcription is a multistep and multiscale process, involving genomic organization (including chromatin folding and E-P communication) and upstream-to-downstream regulation. To establish a general model of transcription, we assume that there are in total *N* nucleosomes on chromatin, and denote by 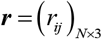 the spatial positions of these nucleosomes in 3D space. In addition, we assume that the gene has in total *K* transcriptionally active and inactive states, among which transitions occur on a slow timescale compared with chromatin motion occurring on a fast timescale. Denote by ***s*** the vector of all the gene states.

Let ***p***(***r***, ***s***;*t*) be a vector of the joint probability density functions (PDFs) that nucleosomes are in position ***r*** = [***r***_1_, ⋯, ***r**_N_*]^T^ and the gene is in state ***s*** = [*s*_1_, ⋯, *s_K_*]^T^ at time *t*, where T is transpose. Note that ***p***(***r***, ***s***;*t*) = [***p***(***r***, *s*_1_;*t*), ⋯, ***p***(***r***, *s_K_*;*t*)]^T^ can be regarded as a mapping: 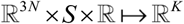 where *S* = {*s*_1_, ⋯, *s_K_*}. Based on the principle of total probability, ***p***(***r***, ***s***;*t*) can be written as the product of the PDF 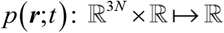 and the conditional PDF (cPDF) 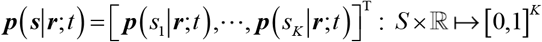, that is,

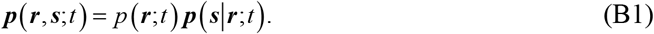

Differentiating Eq. (B1) with respect to time yields

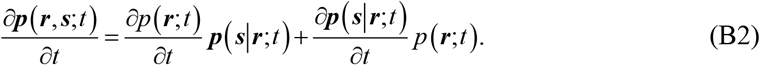

Considering that 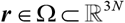 is a “fast” variable for a continuous trajectory, where Ω is a connected and bounded domain. Then, derivative 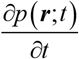 in Eq. (B2) can be formally written as

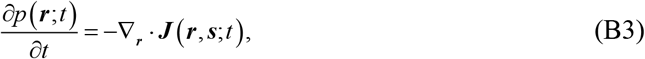

where ***J***(***r***, ***s***; *t*) = ***F***(***r***, ***s***; *t*)*p* (***r***; *t*) is the probability flux and ***F*** (***r***, ***s***; *t*) is a velocity field. If we only consider isotropic diffusion and friction across the region of nucleosome positions, then ***F*** (***r***, ***s***; *t*) can take the following generalized Fokker–Planck approximation:

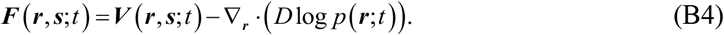

The first term on the right hand side of Eq. (B4) represents the deterministic part of the velocity field. The second term is a stochastic ingredient of the velocity following Fickian diffusion with a diffusion matrix that is assumed as the form of ***D*** = *D**I***, where *D* is a diffusion coefficient and ***I*** is the identity matrix. If we assume that changes in gene state do not contribute to chromatin motion, the velocity field ***V***(***r***, ***s***;*t*) can be approximated by ***V***(***r***;*t*) and 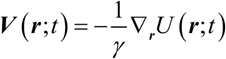, where *U*(***r***;*t*) is the energy field and *γ* is a drag coefficient.

Substituting Eq. (B4) into Eq. (B3) yields

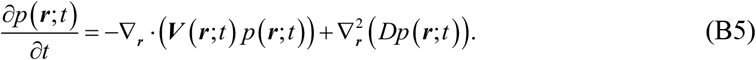

Since the diffusion of a chromatin chain within the cell nucleus is bounded and is often restricted to a subcellular compartment (Ω) that would have complex geometry, it is necessary to impose constraints of probability flux ***J***(***r***;*t*) on the boundary *∂*Ω of Ω. Here we choose a homogeneous Neumann boundary condition (no-flux), i.e., ***J***(***r***;*t*) · ***n***(***r***) = **0** for all ***r*** ∈ *∂*Ω, where ***n***(***r***) is the unit outward normal to the boundary *∂*Ω. The Langevin equation or stochastic differential equation corresponding to Eq. (B5) is

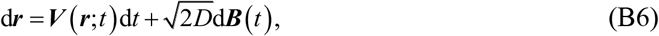

where ***B***(*t*) are independent Brownian motions or Wiener processes (representing Gaussian white noise).

On the other hand, consider that gene state ***s*** is a “slow” variable in a discrete pathway. Then, 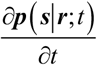 features the process of gene-state switching, and can be written by a master equation:

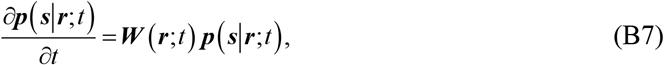

where 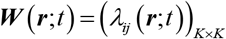 is a nucleosome position-dependent state transition matrix, satisfying the conservative condition: 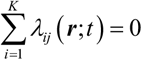.

Substituting Eq. (B5) and Eq. (B7) into Eq. (B2), Eq. (B1) yields the following equation

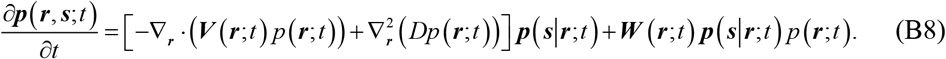

It is reasonable to assume that tiny changes in nucleosome coordinates do not alter gene state, implying that the derivative of conditional probability, ∇***_r_p***(***s|r***;*t*), approximately equals zero partly because of the time interval of downstream gene state transition is generally longer than that of upstream chromatin motion (44). Thus, according to Eq. (B1), we can further obtain the following equation

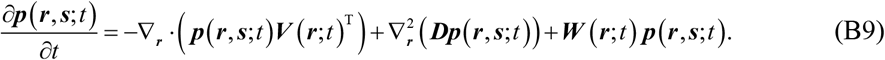

Summing all states ***s*** on both sides of Eq. (B9), we can in turn derive Eq. (B5). Also, integrating all positions ***r*** over Ω on both sides of Eq. (B9) yields the following integral equation for ***p*** (***r***, ***s***; *t*)

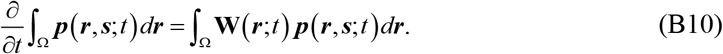

#### B2 Simulation algorithm for dCKE

Since Eq. (B9) involves the spatial derivative ∇_*r*_, the traditional Gillespie stochastic simulation algorithm (45) cannot be directly applied. Here we propose an improved version of this algorithm to solve Eq. (B9).

Assume that there are *L* reactions involved in the gene-state switching process. The *l* th reaction propensity is denoted by *a_l_* (***r***, ***s***;*t*), *l* = 1, 2, ⋯, *L*, and the total reaction propensity is calculated according to 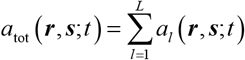. Note that ***s*** is the vector of all gene states, but in the following algorithm, we set *K*-dimensional state vector 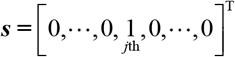 when the current state is *j* in numerical simulation. Element *v_ij_* of stoichiometric matrix (*v_ij_*)_*K×L*_ denotes the net change in gene state *j* due to each reaction *i*, and ***v**_μ_* is the *μ* th column of (*v_ij_*). Let *H*(***r***, ***s***; *t*) is the survival probability that chromatin position is ***r*** and gene state is ***s*** state at time *t*. The main steps for solving Eq. (B9) are listed below:

1. Set the initial state as ***r***_0_ = ***r***(*t*_0_), ***s***_0_ = ***s***(*t*_0_).
2. Generate two random variables *u*_1_ and *u*_2_ distributed uniformly in interval (0,1).
3. Integrate the system of stochastic differential equations

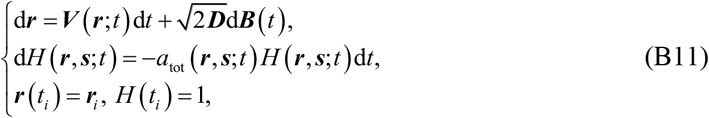

from time points *t_i_* to *t_i_* + *τ_i_*, and with the stopping condition *H*(***r**_i_*, ***s**_i_*;*t_i_* + *τ_i_*) = *u*_1_.
4. Update time and position: *t*_*i*+1_ = *t_i_* + *τ_i_*, ***r***_*i*+1_ = ***r***(*t*_*i*+1_).
5. Choose *μ* such that

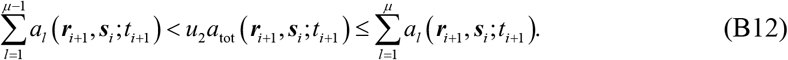
6. Update promoter state: ***s***_*i*+1_ = ***s**_i_* + ***v**_μ_*.
7. Reiterate the system from step (2) with a new state until a given largest time *t*_max_ is reached.

By the above algorithm steps, we can generate sample trajectories of the system described by Eq. (B9).

### C Modelling transcriptional bursting by dCKE

#### C1 Stochastic model of upstream chromatin motion

We model chromatin as a folded polymer chain, a collection of monomers connected by springs. We assume that a monomer represents a nucleosome, and a nucleosome represents an enhancer whereas another nucleosome represents a promoter. Also, we posit that there are only one enhancer and one promoter on the chromatin (i.e., polymer) although multiple enhancers may exist, and neglect attraction or repulsion between other monomers in the chain except for the communication between the enhancer and the promoter. Then, the chromatin motion can be described by the Langevin equation (i.e., (B6))

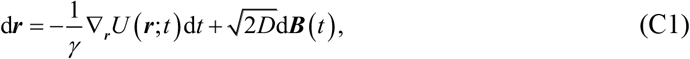

where *U*(***r***;*t*) is the total potential of the polymer chain at time *t*. We make decomposition 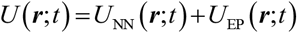, where *U*_NN_ (***r***;*t*) is the potential of the successive monomers and *U*_EP_ (***r***;*t*) is that of the E-P communication.

To simplify but without loss of generality, harmonic springs with stiffness *k*_NN_ are used between the monomers of nearest-neighbors along the chain. Thus, the potential of the successive monomers is

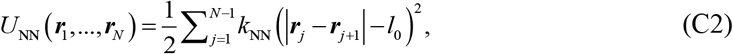

where ***r***_*j*_ =(*r*_*j*1_, *r*_*j*2_, *r*_*j*3_) is the position of the *j*th bead in space and *l*_0_ is the mean distance between neighboring monomers.

For the E-P communication part, one may consider several forms of the potential. In the following, we introduce two most commonly used forms.

##### 1. Harmonic potential

In the main text, the E-P communication is characterized by the harmonic spring, and the corresponding potential is assumed to take the form

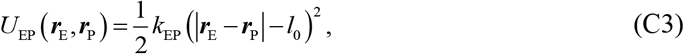

where *k*_EP_ is the spring coefficient between the enhancer and the promoter (also called communication strength), and E, P ∈{1, ⋯, *N*} represent the enhancer and the promoter with E < P. Let *d*_G_ = P – E represents the E-P genomic distance (SI Fig. 1).

For analysis simplicity but without loss of generality, we set *l*_0_ = 0 in Eq. (C2) and (C3). Since all the interactions we consider in the main text are represented by harmonic potentials, Eq. (C1) can be rewritten as

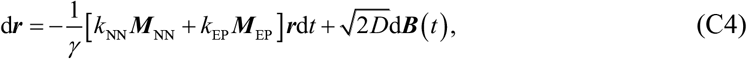

where ***M***_NN_ is a *N*×*N* matrix showing the connectivity between adjacent beads, given by

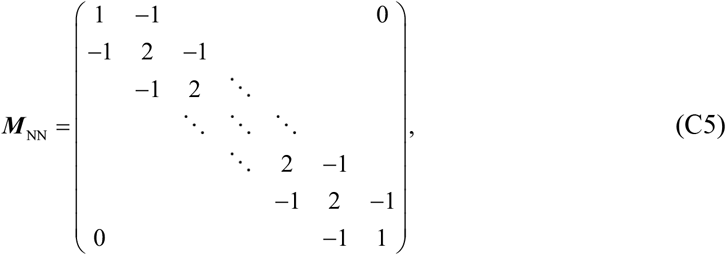

and *M*_EP_ is also a *N*×*N* matrix representing the E-P interaction, given by

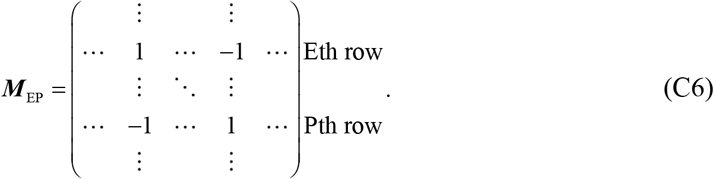

Eq. (C4) with Eq. (C5) and (C6) characterize the movement of chromatin.

**SI Figure 1.**
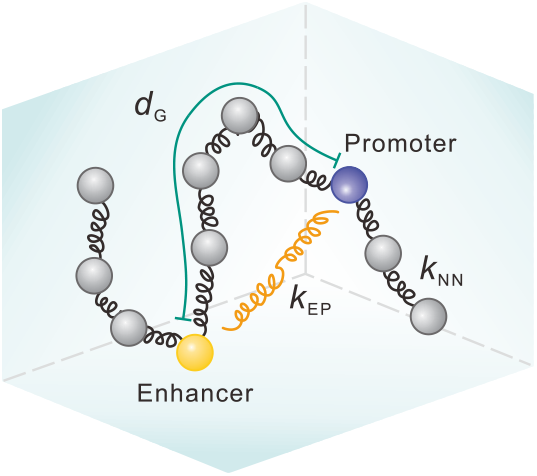
Schematic depiction of symbols related to chromatin structure.

##### 2. Lennard-Jones potential

Besides the harmonic potential, the Lennard-Jones (LJ) potential is also commonly used in simulating molecular dynamics. Here, we modify the LJ potential function to fit our model. That is

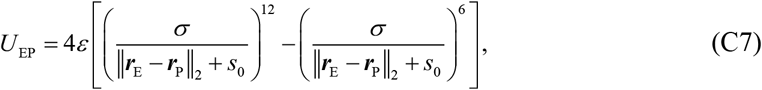

where *σ* is a distance parameter, *ε* is the depth of the potential well representing the strength of attraction between two particles, and *s*_0_ is a modified distance. Based on Eq. (C7), derivatives ∇_***r***_E__ *U*_EP_, and ∇_***r***_P__ *U*_EP_ can be easily obtained. Then, Eq. (C1) can be rewritten as

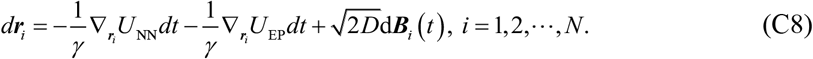

Eq. (C8) describes chromatin motion. Note that the second term on the right hand side of Eq. (C8) is zero for ***r**_i_* except ***r***_E_ and ***r***_P_. Apparently, Eq. (C8) that considers the LJ potential of complex form is different from Eq. (C4) that considers the potential of comparatively simpler form. In general, the former is more difficult to analyze and solve than the latter.

Finally, we point out that in subsection C1, communication strength *k*_EP_ depends, in general, on the inherent property of regulatory elements and the microenvironment around them. Different enhancers correspond to different communication strengths, so E-P communication strength is gene-specific. The E-P communication activating transcription (i.e., enhancer-dependent transcription activation) often requires the presence of multiple TFs, such as general TFs and sequence-dependent effectors of signaling pathways, to ensure the integration of internal and external environmental cues via the E-P communication. Therefore, communication strength *k*_EP_ is taken as a key parameter of our model.

#### C2 Multi-state model of downstream transcriptional bursting

##### 1. Four-state model for transcription cycle

In eukaryotic cells, mRNAs are usually produced in a burst mode that a high activity state follows a long refractory period. Such a biological phenomenon is frequently described by a telegraph model of gene switching, which consists of two states: an OFF state where the gene is not expressed and an ON state where the gene is transcribed. The two-state model provides a simple quantitative analysis framework, but transcription bursting is a complex biochemical process. Some gene-expression processes, e.g., those involving complex promoter (or E-P) dynamics, TFs and cofactors (COFs) dynamics and Pol II dynamics, would not be described by simple two-state models. Recent studies have extended two-state models to multistate models. In our main text, we used a four-state model to simulate transcriptional bursting dynamics with biological details stated below for better understanding.

**SI Figure 2.**
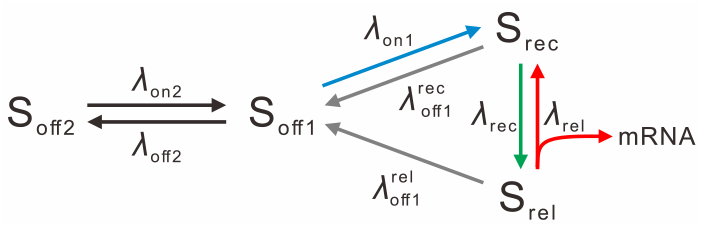
Kinetic scheme for a four-state model of transcriptional bursting.

###### OFF states

In the case that chromatin is silent, the mechanism of how a suitable transcriptional state for DNA transcription is created is unclear, but constantly increasing data have shown that the non-permissive period follows a non-exponential distribution (13, 46), indicating that the process from non-permissive to permissive transcription states is memorial, and is more complex rather than a single-step Markov process that is memoryless. This memory implies that a single OFF state in the traditional telegraph model is not enough to characterize the non-permissive state of chromatin. In addition, chromatin remodeling and nucleosome unwrapping due to the accessibility of chromatin are necessary for gene transcription. Pioneer TFs have unique properties that can open closed chromatin (47) to expose the promoter motif. Subsequently, the promoter, which is in a state of primed burst, employs general TFs, Mediator and Pol II for transcription. For this purpose, a deep inactive state (S_off2_, closed chromatin) and a primed burst state (S_off1_) are considered as the OFF state (S_OFF_, non-permissive period).

###### ON states

Besides the non-permissive period (inactive state S_OFF_), two most important processes in the transcription cycle are Pol II recruitment to the promoter and Pol II pause release from promoter-proximal (9, 48). Pol II is necessary for transcribing DNA into mRNA and paused Pol II is a common step in gene transcription and regulation (49, 50). Recent studies have indicated that paused Pol II is more stable than Pol II in the PIC stage (38, 51), and promoters with Pol II are more sensitive in response to stresses than those lacking Pol II (52). Meanwhile, paused Pol II can prevent new Pol II initiation to reduce transcriptional noise (38). After the release of paused Pol II, the next Pol II recruitment can be carried out. Since Pol II recruitment occurs after burst initiation (53), we employ Pol II recruitment state (S_rec_) and Pol II pause release state (S_rel_) as state ON (S_ON_, permissive period) in the bursting stage. These two states, S_rec_ and S_rel_, can be switched to each other. And we assume that the process from S_rel_ back to S_rec_ generates an mRNA.

##### 2. State switching in four-state model

The bursting system to be studied is described by a set of biochemical reactions on a slow timescale in contrast to the introduced-above system of chromatin motion on a fast timescale (see Section A and SI Fig. 2)

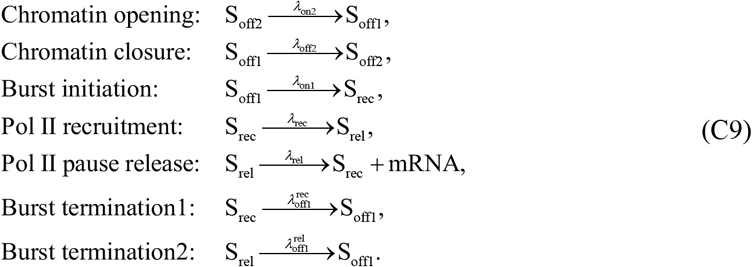

Note that rates *λ_c_*, *c* ∈{off2,off1,on2,on1,rec,rel} in Eq. (C9) are effective rates that summarize a series of sub-processes. Since burst termination is relatively dynamic (54), we hypothesize that both S_rec_ and S_rel_ states can return to S_OFF_. Also because S_off2_ is a deep inactive state, it is unlikely for S_rec_ and S_rel_ states to return directly to S_off2_ state in one step. Thus, we posit that only S_off1_ can be switched to S_off2_. Since the model has two burst termination channels, we add a superscript to distinguish these two termination rates (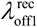 and 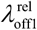). In order to help the readers’ understanding and memory, we state the following simple facts. The toggle between states S_off2_ and S_off1_ is regulated by the chromatin or nucleosome opening and rewrapping with rates *λ*_on2_ and *λ*_off2_, respectively. The transition from S_off1_ to S_rec_ is associated with TFs binding with rate *λ*_on_. The state switching from S_rec_ to S_rel_ is mediated by Pol II recruitment with rate *λ*_rec_. The transition from S_rel_ to S_rec_ is linked to Pol II pause release with rate *λ*_rel_ (which is also the rate of mRNA production). The transition from S_rec_ to S_off1_ is related to the TFs unbinding, the disruption of TFs binding site or other processes with rate *λ*_off1_, and the switching from S_rel_ to S_off1_ is involved in TFs and Pol II unbinding from the target gene, the collision of Pol II with exoribonucleases, or the disruption of Pol II active site (54) with rate *λ*_off3_. Note that before the gene transitions to state S_OFF_, there would be many switches between S_rel_ and S_rec_, thus producing bursts.

##### 3. Transcriptional bursting process

For the above biochemical reaction network, the characteristic of burst initiation is that S_off1_ switches to S_rec_ whereas that of the burst termination is that the S_ON_ state returns back to the S_off1_. If switching between S_off2_ and S_off1_ is relatively slow, this will result in periods of gene complete refractory, interspersed with sporadic bursty transcription.

Since the degradation of mRNA is not considered in the model, we assume that when the Pol II is released from the promoter-proximal pause, the transcription elongates along the genome for a fixed time *τ_E_*, and when the Pol II reaches the end of the gene, the Pol II falls off and the nascent mRNA is no longer detected by snapshots.

Note that although our model is a multi-state one, it is different from multi-state models in previous studies (55, 56). Previous models assumed that gene state does not change when mRNA is produced (i.e., ON →ON+mRNA), implying that the detailed processes of transcriptional bursting are ignored. In contrast, our model assumes, based on biological phenomena, that mRNA is produced in the process of switching from one state to another (i.e., S_rel_ →S_rec_ + mRNA). This is crucial for capturing the feature that only one Pol II is permitted to bind to promoter and the second Pol II recruitment must occur after the first Pol II pause release. Also, our model can capture some characteristics (such as traveling ratio, the effect of altering Pol II pause release rate) that cannot be obtained by the previous models.

##### 4. Transition probability matrix *W*

In Eq. (C9), each state switching is regarded as a reaction process. However, in *vivo*, each state transition is generally not a single reaction but would be the synergism of multiple reactions (SI Table 1). Although there would be many sub-reactions behind each reaction, we suppose that the reaction events in the four-state model, i.e., Eq. (C9), are Markovian, i.e., the probabilities of reaction events only depend on the current state of the system, independent of the prior history (this hypothesis was made in almost previous theoretical studies). With this hypothesis, the waiting time of each reaction follows an exponential distribution. If a process occurs via multiple sub-reactions (or multiple steps) but only one of these steps is rate-limiting, then this process can be effectively described by one step. The number of rate-limiting steps of a reaction process is usually not large in gene transcription, so our Markov model can work well (15).

**SI Table 1:** Molecular processes of transcriptional bursting

| Gene state |  | Reaction | Biological processes |
| --- | --- | --- | --- |
| OFF | Deep inactive state ($S_{\text{off}2}$) | $S_{\text{off}2} \rightarrow S_{\text{off}1}$ | Pioneer TFs binding (57), **opening chromatin** (58) |
| | Primed burst state ($S_{\text{off}1}$) | $S_{\text{off}1} \rightarrow S_{\text{rec}}$ | **TFs recruitment** (59), the binding of activators on enhancer (60), the distortion of TATA box (35) and alteration of chromatin structure (60). |
| ON | Pol II recruitment state ($S_{\text{rec}}$) | $S_{\text{rec}} \rightarrow S_{\text{off}1}$ | TFs unbinding, disruption of TFs binding site |
| | | $S_{\text{rec}} \rightarrow S_{\text{rel}}$ | **Pol II recruitment** (9), the formation of PIC (61), promoter escape (62) and Pol II pausing (63). |
| | Pol II pause release state ($S_{\text{rel}}$) | $S_{\text{rel}} \rightarrow S_{\text{off}1}$ | TFs and Pol II unbinding from the target gene. |
| | | $S_{\text{rel}} \rightarrow S_{\text{rec}} + \text{mRNA}$ | **Pol II pause release** (64), productive elongation (65), Pol II termination (54) and another round of transcription (9). |

Based on the above description, the state transition matrix ***W***(***r***;*t*) is expressed as

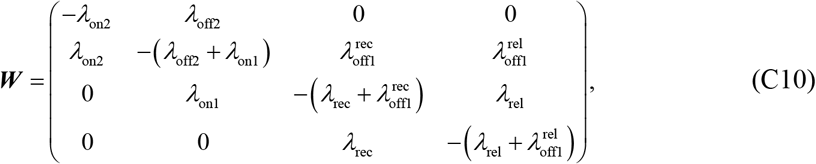

where all the *λ* – type parameters are dependent on the E-P spatial distance and will be discussed at Section C3.

#### C3 Information flow from chromatin conformation to transcriptional bursting

##### 1. Models of E-P communication

Regulatory information from distal enhancers to target genes is needed to execute accurate transcription. Although various mechanisms have been proposed (such as loops and hubs), how E-P communication is carried out is still a matter of debate (66, 67). The looping model assumes E-P direct contact through molecular complexes. This assumption was often made in previous studies with a reason that looping can directly increase transcription levels and reactivate developmental genes (40, 68, 69). However, in the work of Rao et al. (70) and Long et al. (71), active E-P pairs did not show an increased contact frequency, indicating that forming an E-P loop is not needed. Another viewpoint is the occurrence of transcription hubs, mainly because E-P interactions and the assembly of PIC facilitate that transcription loci forms clusters or hubs (66). Hypothetical hubs can well interpret the large spatial distance of E-P when transcription is activated. Therefore, the enhancer may not be too close to the target promoter. Recent experimental studies on measuring E-P spatial distances suggested a large distance on the order of a few hundred nanometers (nm) (25, 37). For example, the distance between the Sox2 region and its essential enhancer in ESCs ranges from 200-400nm (72).

##### 2. Necessity of E-P proximity

E-P proximity has been believed to increase the likelihood of transcription bursting. However, proximity does not mean direct contact. The distance between PP7 reporter gene and the enhancer decreases in transcriptional activation in contrast to the inactive state case, but the E-P distance in ON state is ~340nm (37), observably not via the direct contact distance. Throughout this paper, we use the hub hypothesis to investigate the transcription mechanism.

##### 3. Modeling information flow from chromatin structure to transcriptional bursting

E-P communication, which plays a vital role in regulating stochastic gene expression, should be taken into account in transcriptional bursting. First, transcription activators binding to enhancers recruit TFs to alter chromatin structure and make it more accessible to active transcription (60). Second, E-P interaction co-regulates the recruitment of TFs and Pol II and the assembly of PIC (73, 74). Besides, enhancers recruit Mediator complex or histone acetyltransferase p300 to help the Pol II on promoters initiate transcription (75). Third, enhancers promote dissociation of NELF by recruiting COFs to affect Pol II pause-release on promoter-proximity (76). Correspondingly, gene state transition rates *λ_c_* of the biochemical reactions govern transcriptional bursting. According to the above experimental observations or evidences, we assume that the state transition rates are related to the E-P spatial distance denoted by *d*_S_. However, biological experiments did not tell us how these rates quantitatively depend on *d*_S_. In fact, this dependence relationship would be complex and in particular, it would be organism-specific. To simplify our analysis, we will, by making assumptions, set a special but common form of *λ_c_* to link the downstream transcription to the upstream chromatin configuration.

Note that the opening chromatin (from S_off2_ to S_off1_) would be mainly associated with pioneer TFs. Thus, we assume *λ*_off2_ is a constant independent of E-P spatial distance. Additionally, since the relationship between the *λ_c_*, *c* ∈ {on1,rec,rel} and E-P distance may be opposite to that between *λ_c_*, *c* ∈ {off1,off2} and the distance, we set rates *λ*_off2_, 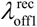 and 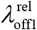 as constants for simplicity.

First, chromatin structure generally does not act as a binary switch but acts as a modulator of gene function (non-binary) (1). Second, cooperative and synergistic binding to DNA is a common way in organisms. For example, in transcription processes, the binding of TFs may affect the binding rate of other TFs, Mediators or Pol II. Thus, it seems more reasonable to assume that the E-P spatial distance affects transcriptional burst rates in a nonlinear manner. Here we use Hill functions, which are very successful in modeling biological phenomena (77), to describe the transcription rates. To sum up, we assume that the piecewise continuous nonlinear rate function related to the E-P topological structure is

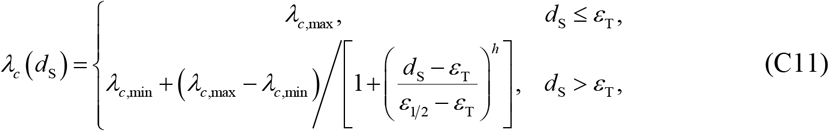

where *λ*_*c*,min_ and *λ*_*c*,max_ are the minimum (basic) and maximum reaction rates for reactions *c* ∈{on1,rec,rel}, *ε*_1/2_ is the spatial distance when *λ_c_* is equal to (*λ*_*c*,max_ – *λ*_*c*,min_)/2, *h* is a Hill coefficient that controls how steep the rate curve is, and *ε*_T_ is a distance threshold (SI Fig. 3A and 3B). When the E-P distance is less than *ε*_T_, the *ε*_T_ is also used to describe the encounter of E-P. Therefore, it is reasonable to assume that the state transition rates are maximum when the distance is less than *ε*_T_ in Eq. (C11). Besides, encounter, which is merely an assumption in physical statistics, does not mean direct contact. As the distance increases, state transition rates can reduce to the minimum. Eq. (C11) can illustrate how the transition rates vary with E-P topologies, and further indicate that transcription is regulated at any time.

In our numerical simulation and theoretical analysis, we sufficiently use the relationship described by Eq. (C11). However, it is needed to point out that parameters *λ_c_*, *c*∈{on1,rec,rel} may take other forms except for form (C11). In general, *λ_c_* may be set nonlinear decreasing functions of *d*_S_. Also, *λ_c_* can be set increasing functions to explain the special case that the enhancer activation of increasing E-P spatial distance *d*_S_ (78). Finally, *λ_c_*, *c* ∈{off1,off2} may take a form of functions that are not constants.

**SI Figure 3.**
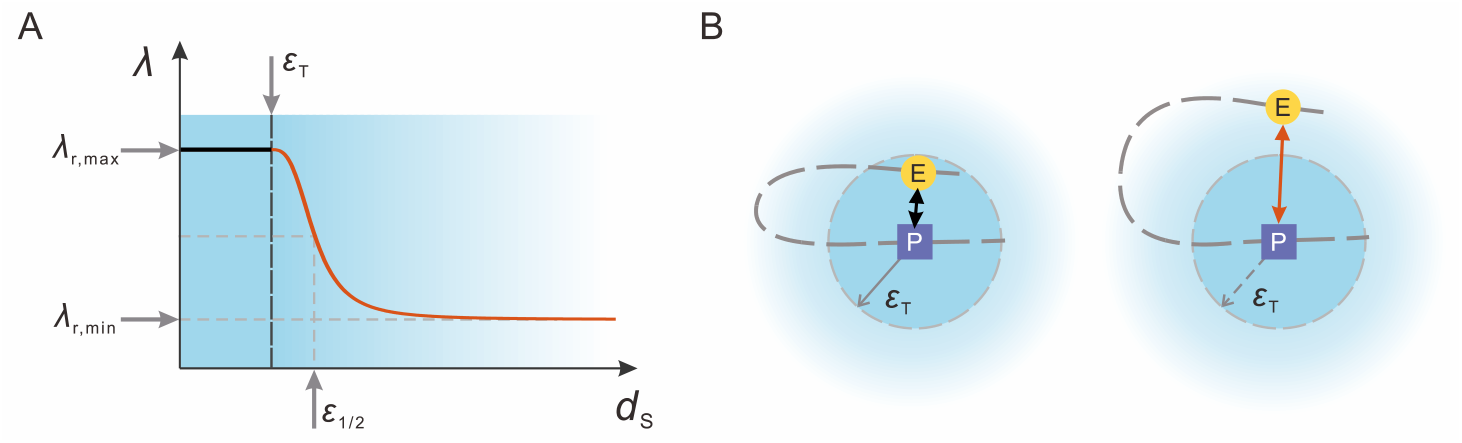
Schematic for the relationship between E-P spatial distance and gene state transition rates. (A) A piecewise nonbinary nonlinear Hill function describes the relationship between E-P spatial distance and burst rates. (B) E-P spatial distance is less (more) than *ε*_T_ in the left (right) panel. For the convenience of display, we take the promoter as the center position.

### D Simulation and statistical analysis of the whole model

#### 1. Numerical simulations of the whole model

The entire model, which considers both chromatin architecture and transcriptional bursting, is simulated using MATLAB (79). A numerical method has been described in the above Section B2. We examine a system consisting of *N* = 100 monomers in simulations. To simplify, we consider a spatial region isolated from neighboring DNA by boundary insulator elements (80, 81). Biologically reasonable values of all the model parameters are listed in Section H.

For a given set of parameters, we simulate 10^3^ gene copies, proceeding through 10^7^ seconds in time. Each simulation starts in the S_off2_ state. Before taking samples from every simulation, we run 10^4^ seconds to ensure equilibration between conformation and transcription. After that, snapshots of the system are taken every 100 seconds.

#### 2. Statistical analysis of simulation results

##### Statistics analysis of upstream chromatin dynamics

Using the data obtained by simulation, we perform statistical analysis for the chromatin structure, especially for the E-P spatial distance. Specifically, we calculate the PDF of E-P spatial distance based on the produced time series data. Also, we calculate the E-P encounter probability (referring to SI Fig. 10E). Furthermore, we can alter the encounter distance to survey the E-P effective density or concentration. That is, we calculate the E-P encounter probability at some sphere with radius *R* (E-P encounter distance is 2*R*), and then normalize this probability by the sphere volume as an effective density or concentration (referring to SI Fig. 10F, (82)).

##### Statistics analysis of downstream bursting kinetics

According to the reaction time series, we calculate the probability mass function (PMF) of burst size and the PDFs of the dwell time in S_OFF_ and S_ON_ states and the cycle time (the time from S_OFF_ to S_ON_ and back to S_OFF_, referring to SI Figs. 13E-H). Furthermore, we calculate some statistical quantities such as mean burst size (MBS), mean dwell-time (MDT) in ON/OFF state, mean cycle time (MCT), burst frequency (BF). Here MBS is defined as the average number of mRNA molecules produced per burst whereas BF as the average number of bursts occurred in a given time interval. In other words, BF is the reciprocal of the MCT. In addition, our model can also be used to compute the mean traveling ratio (MTR), which is defined as the ratio of Pol II density in the gene body divided by the Pol II density at promoter proximity. Based on the data produced by simulation and by dividing the Pol II into two types upon its location: Pol II at promoter-proximal (5’ end of the strand, Pol II at S_rel_ state) and Pol II in the gene body (3’ end of strand, a fixed time interval after Pol II pause release back to recruitment state, SI Fig. 4B), we calculate the MTR by uniformly-spaced sampling data (referring SI Fig. 19).

### E Theoretical analysis of transcriptional bursting

#### E1 Timescale separation method decoupling upstream and downstream kinetics

Note that burst size, dwell time and cycle time are random variables, denoted by *BS*, *DT* and *CT*, respectively. We mainly calculate the PDFs or PMF of these random variables (SI Fig. 4) and their statistical quantities such as expectations. Also note that these variables are related to the state switching rates that however are linked to the E-P spatial distance due to the fact that the burst rates are regulated by E-P spatial distance (Section C3).

**SI Figure 4.**
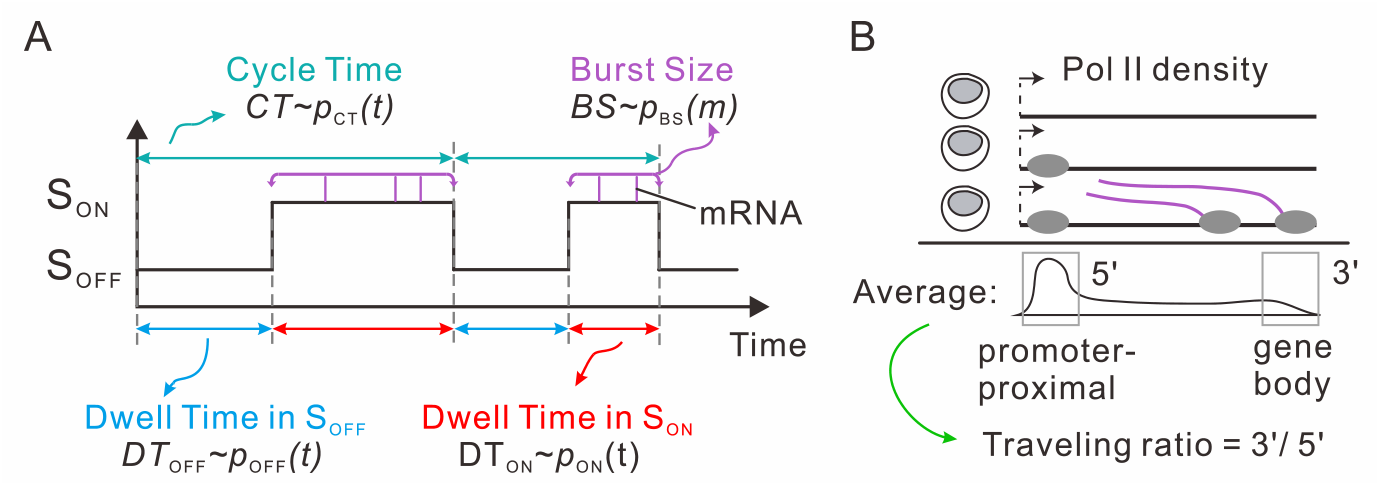
Meanings of symbols related to downstream transcriptional bursting. (A) Schematic diagram of burst size, dwell time in ON/OFF state and cycle time. (B) Schematic diagram of traveling ratio.

##### 1. Timescale separation parameter

Since the biochemical reactions for downstream transcriptional bursting occur on a slow timescale whereas the upstream chromatin motion takes place on a fast timescale, we can use a timescale separation method to carry out a theoretical analysis of bursting kinetics, although the timescale gap between the upstream and the downstream may be different in different cells or during different periods in the same cell. This method has been successfully used in the analysis of gene regulatory networks (83, 84) and the mutual information that disentangles interactions from changing environments (85). To quantify the timescale separation, we introduce a parameter *ω*, which is defined as the ratio between the propensities of the reactions on fast and slow timescales

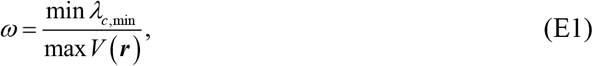

where *λ*_*c*,min_, *c* ∈{on1,rec,rel} are the gene state switching rates in downstream transcription processes and *V*(***r***) is the velocity field between the enhancer and the promoter in the upstream. In the following, we give detailed explanations.

##### 2. Upstream chromatin dynamics

The random motion of chromatin leads to fluctuations in the E-P spatial distance (*d*_S_). Let *DS* be the random variable representing the E-P spatial distance. In principle, *DS* should follows a distribution, i.e., *d*_S_ ~ *p_DS_*(*d*_S_). In Section E2, we will derive the analytical expression of E-P spatial distance distribution *p_DS_* (*d*_S_) (referring to SI Figs. 10A and 10C).

In Eq. (E1), the E-P velocity field equals 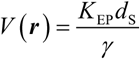, where *K*_EP_ represents the total spring coefficient between the enhancer and the promoter (see Section E2). We choose the value of *d*_S_ when cumulative density function (CDF) of *p_DS_* (*d*_S_) reaches 0.99, implying that the *d*_S_ reaches the maximum. This setting that corresponds to the maximum velocity is to include all possible cases of E-P spatial distance in simulations.

##### 3. Downstream transcriptional bursting

For downstream random variables *X* ∈ {*BS,DT|S,CT*}, we calculate the joint distribution *p_X,DS_* (*x, d*_S_), which accounts for fluctuations in *d*_S_ and their effect on fluctuations in burst.

In the case that the fluctuations in *d*_S_ are much faster compared with the rate of transcription, the *d*_S_ fluctuations can be averaged out, and the corresponding joint distribution denoted by 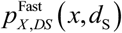 is given by

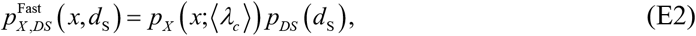

where 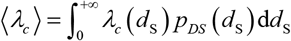. In this case, the marginal PDF (or PMF) of random variable *X* takes the form (referring to SI Figs. 12C-D)

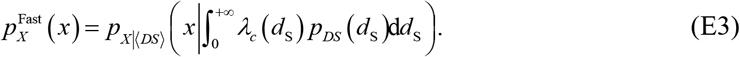

On the contrary, if fluctuations in *d*_S_ are much slower, the corresponding joint distribution denoted by 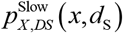 is given by

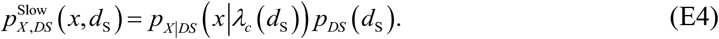

In this situation, the marginal PDF (or PMF) of random variable *X* takes the form (referring to SI Figs. 12E-F)

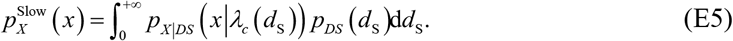

In general, the marginal PDF (or PMF) of *X* ∈{*BS,DT|S,CT*} is a mixing distribution obtained by weighting the marginal PDF (or PMF) given in Eq. (E3) and Eq. (E5) (referring to SI Figs. 13A-D)

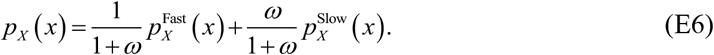

Based on Eq. (E6), the PMF of burst size *p_BS_*(*m*), the PDF of dwell time *p_S_* (*t*) (note: the complete expression is *p_DT|S_*(**t|s**) where *s* ∈{off2,off1,rec,rel} or *s* ∈{OFF,ON}) and the PDF of cycle time *p_CT_*(*t*) can be obtained (SI Fig. 4A). In particular, the expectation of *X* can be expressed as

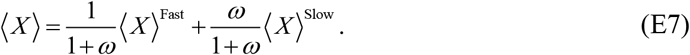

Finally, we point out that since the traveling ratio is the ratio of two quantities as defined in Section D, expectation *E*[*TR*] can be directly derived, where *TR* is a random variable representing traveling ratio (SI Fig. 4B).

In Section E3, we will derive the analytical expressions of the burst size, dwell time, cycle time and traveling ratio when we only consider downstream transcriptional bursts. Note that the PDF or PMF obtained in Section E3 is actually the cPDF. Then, the PDF or PMF of the whole transcriptional bursting can be obtained by Eq. (E6).

#### E2 Analytical results for upstream chromatin dynamics

In this subsection, we derive the explicit expression of the E-P spatial distance distribution. Term *k*_NN_***M***_NN_ + *k*_EP_***M***_EP_ in Eq. (C4) is a singular matrix. To eliminate the degrees of freedom, the position of the first monomer in the chain can be set as ***r**_1_* ≡ **0** based on the methods in (86). Thus, Eq. (B5) can be rewritten as

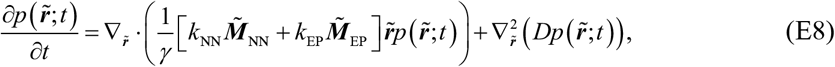

where 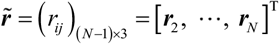 and

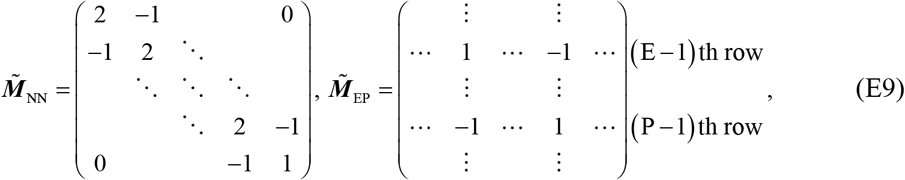

where .1 < E < P..

Owing to the fact that every monomer moves independently in each dimension, the PDF for chromatin conformation 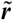 can be expressed as

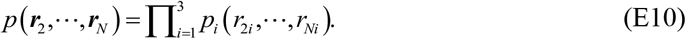

In fact, the monomer motion in Eq. (E8) is an Ornstein-Uhlenbeck process and the general solution to this equation is a Gaussian distribution. If we consider one-dimensional PDF *p_i_*, the *p_i_* can be analytically expressed as (87)

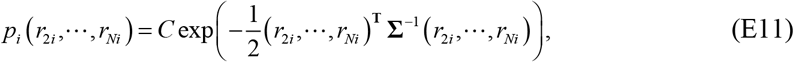

where *C* is a normalization constant and

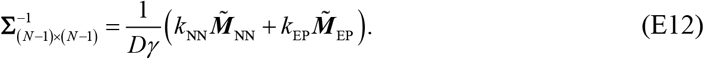

The *p_i_* is a multivariate normal distribution with the zero mean and the covariance matrix being **Σ**. The marginal distributions for the enhancer and promoter are calculated according to

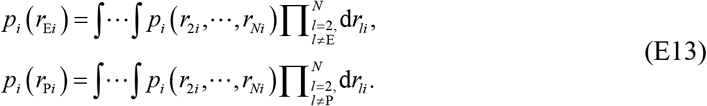

Based on the properties of Gaussian distribution, the marginal PDFs in Eq. (E13) are also Gaussian distributions.

By tedious calculating, we find that the analytical expression of the PDF *p_DS_* (*d*_S_) of the E-P spatial distance takes the form

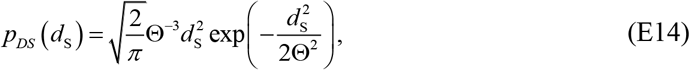

where

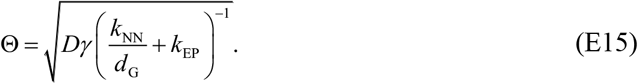

In Eq. (E15), *d*_G_ is the E-P genomic distance, i.e., *d*_G_ = P – E, which is different from the E-P spatial distance *d*_S_ (SI Fig. 1). Note that Eq. (E14) is a Maxwell-Boltzmann distribution (referring to SI Fig. 10A), which can be regarded as the positive square root of the sum of squares of three independent random variables with each following the same normal distribution. Equivalently, each normal distribution represents the E-P distance in a certain dimension. Thus, Eq. (E14) is the distribution of the E-P Euclidean distance in 3D. More precisely, Eq. (E14) is equivalent to the chi distribution with three degrees of freedom and a scale parameter Θ.

Except for the harmonic spring *k*_EP_ between the enhancer and the promoter in Eq. (E15), which accounts for the E-P communication, the enhancer and the promoter are also connected in series by *d*_G_ springs with the same elastic coefficient *k*_KK_. These connected springs are in effect equivalent to a spring with an elastic coefficient *k*_NN_/*d*_G_. Thus, the connection between the enhancer and the promoter can be viewed as two paralleling springs, which are further equivalent to a spring with an elastic coefficient 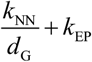. Due to the Einstein relation *Dγ* = *k_B_T*, the distribution is determined by the temperature *T* (or the product of *D* and *γ*, rather than *D* or *γ* alone) and the equivalent spring. Furthermore, the scale parameter Θ in Eq. (E15) measures the spatial distance in units proportional to the square root of the ratio of temperature *T* and spring coefficient 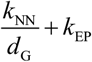. In Eq. (E1), 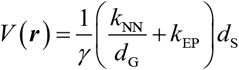.

Here we give details for deriving Eq. (E15). Making use of the fact that every monomer moves independently in each direction in 3-dimensional space, we can show that the lumping parameter Θ is expressed as 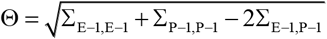 where 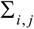 are the elements of matrix **Σ**_(*N*–1)×(*N*–1)_. For convenience, we define 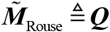 and 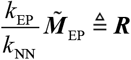. Then, Eq. (E12) can be rewritten as

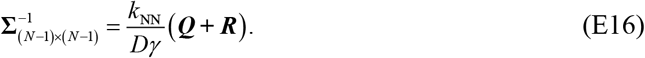

Obviously, the matrices ***Q*** and ***Q*+*R*** are nonsingular matrices, and ***R*** is a matrix of rank one. Thus, the inverse of ***Q*+*R*** is (88)

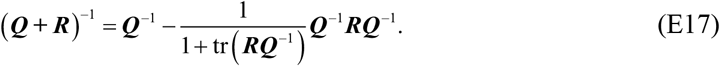

Because ***Q*** is a tridiagonal matrix, its inverse is given by (89)

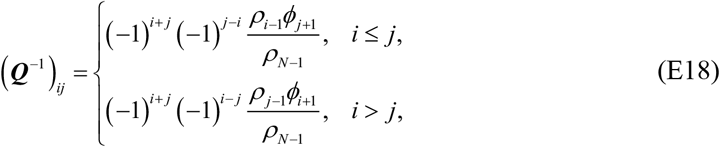

where *ρ_k_* = *k*+1 (*k* = 0,1, ⋯, *N*–2), *ρ*_*N*–1_ = det(***Q***) = 1, *ϕ_k_* = 1 (*k* = 1,2, ⋯, *N*). Then

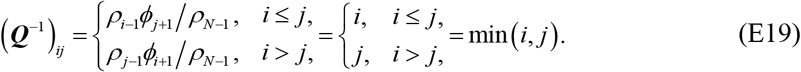

Hence,

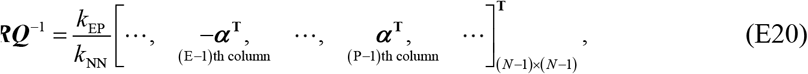

where 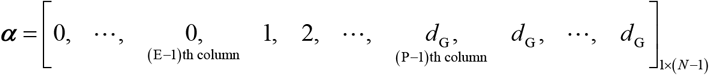. The trace of matrix ***RQ***^−1^ is given by

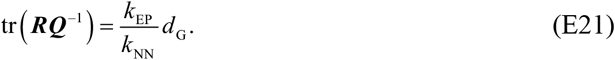

We then have

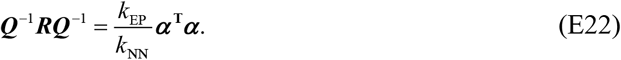

Therefore, based on Eq. (E16) – (E22), we can calculate matrix **∑**_(*N*–1)×(*N*–1)_. However, it is no need to calculate all the elements of **∑**_(*N*–1)×(*N*–1)_. Eq. (E15) can be thus derived.

Finally, we would like to point out: (1) the statement that *d*_G_ is the E-P genomic distance is not accurate, but *d*_G_ is actually the number of monomers between the enhancer and the promoter in simulation. If one monomer is assumed to be a nucleosome, the E-P genomic distance should be expressed as 200*d*_G_bp, where 200bp is the DNA length around the nucleosome; (2) the total genomic length is independent of the E-P spatial distance distribution *p_DS_*(*d*_S_). In numerical simulations, the number of simulated monomers is thus unimportant compared with that of monomers between the enhancer and the promoter, i.e., *d*_G_; (3) the form of modified LJ potential between the enhancer and the promoter in Eq. (C7) merely provides a strategy that replaces the harmonic spring potential. Although the LJ potential is more difficult to analyze, numerical simulations find that LJ potential can be also fitted with Maxwell-Boltzmann distribution (referring to SI Fig. 10G).

#### E3 Analytical results for downstream bursting kinetics

In this subsection, we consider the downstream transcriptional bursting. The PDF or PMF obtained in this subsection is actually the cPDF given the E-P spatial distance *d*_S_.

For our four-state model, an equivalent model with an absorbing boundary state S_abs_ is shown in SI Fig. 5. The following analysis is mainly based on SI Fig. 5. Burst size is unrelated to the state S_OFF_ since only when the gene is activated (S_ON_) can mRNA be produced. Besides, the residence time of the state S_OFF_ can be calculated independently from other states due to the particularity of S_OFF_. Thus, we first focus on the burst stage (i.e., S_ON_) and ignore the state S_OFF_ in the following derivation. Let the initial time of burst stage (*t* = 0) be set at the moment when the state S_off1_ switches to the state S_rec_.

**SI Figure 5.**
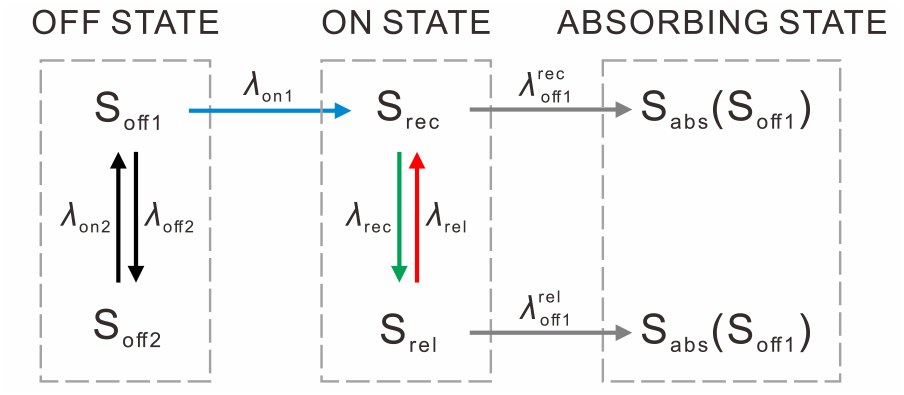
Kinetic scheme of an equivalent four-state model of transcriptional bursting.

Let *H_T|BS,S_* (*t|m,s*) be the survival probability that *m* mRNAs are produced during a burst (i.e., *BS* = *m*, *m* = 0,1,2, ⋯) in the state *S* (i.e., *S* = *s*, *s*∈{rec, rel}) at the time *t* (i.e., *T* = *t*).

Then,

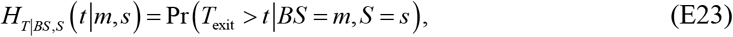

where *T*_exit_ is the exit time from the state *s*. Hereafter we define *H_T|BS,S_*(*t*|–1, *s*)≡0. Note that the expressions of the state *s* ∈ {rec, rel} in formulas and of the state S_rec_, S_rel_ in the main text may be different but their meanings are the same. In the following, we rewrite the survival probability *H_T|BS,S_*(*t|m,s*) as *H_m,s_*(*t*) for the sake of simplicity and beauty of the formulas.

Based on Eq. (E23), we consider an infinitesimal interval (*t,t* + Δ*t*). Then, *H_m,s_* (*t* + Δ*t*) can be expressed as

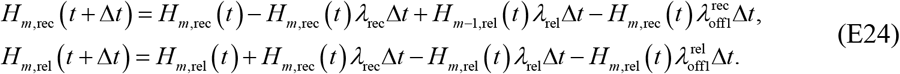

The master equations determining the probabilities *H_m,s_* (*t*) from Eq. (E24) are

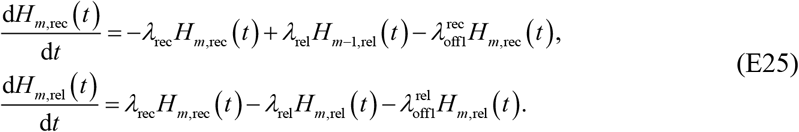

This equation group is valid for all *m* ≥ 0.

To solve Eq. (E25), we introduce the generating function *G_s_*(*z, t*) for *H_m,s_*(*t*)

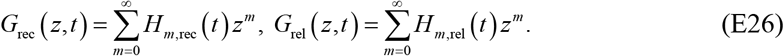

Then, we have the following partial differential equations

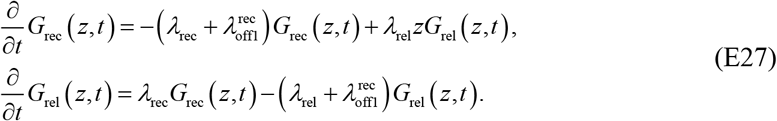

Eliminating *G*_rec_ (*z,t*) from Eq. (E27), we obtain the following second-order differential equation

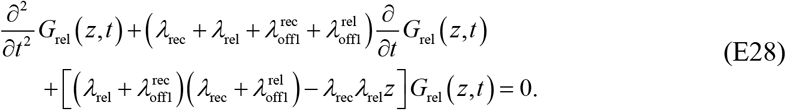

This equation can be viewed as an ordinary differential equation with constant coefficients. The corresponding characteristic equation is

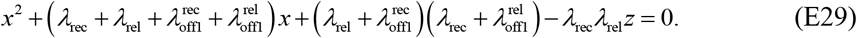

Solving this algebraic equation yields

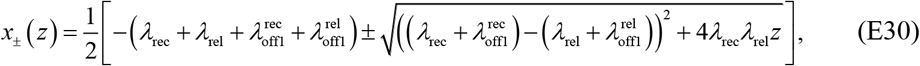

and we can show *x*_±_ < 0. Thus, a general solution to Eq. (E28) is

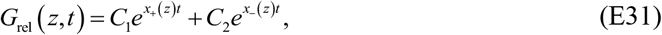

where *C*_1_ and *C*_2_ are constants determined by initial conditions. Substituting Eq. (E31) into the second equation of Eq. (E27), we have

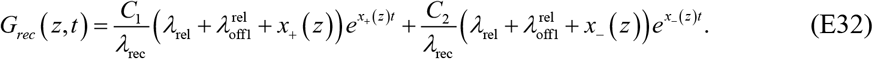

Note that *H*_0,rec_(0) is the initial state of a burst. If taking *H*_0,rec_(0) = 1, *H*_0,rel_(0) = 0 and *H_m,s_*(0) = 0 (*s* ∈ {rec, rel}, *m* = 1,2, ⋯) as the initial conditions, we have the following algebraic equation group determining two constants *C*_1_ and *C*_2_

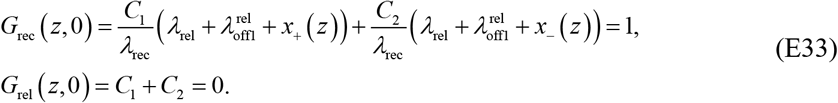

Solving Eq. (E33) gives

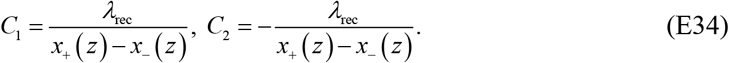

Substituting Eq. (E34) into Eq. (E31) and (E32), we know that the solution of Eq. (E27) is

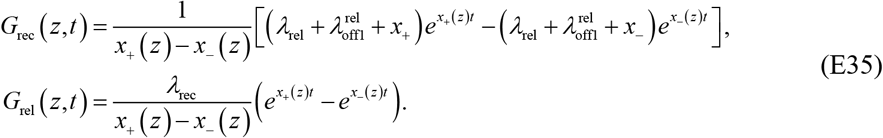

To that end, we have obtained the solution of the generating function corresponding to the survival probability.

##### 1. Burst size

In our four-state model, both S_rec_ and S_rel_ can return to S_off1_ so that the burst ends. And the transitions between S_rec_ and S_rel_ can be carried out many times before one burst ends, resulting in the bursty production of mRNAs.

The probability that burst termination time *T*_term_ falls within an infinitesimal interval (*t*, *t*+Δ*t*) equals

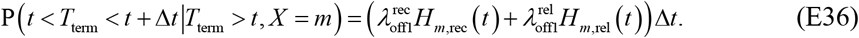

The joint probability density *p*_*BS,T*_term__, (*m,t*) for *BS* (discrete random variable) and burst termination time *T*_term_ (continuous random variable) include all incoming fluxes driving the system from state S_rec_ and S_rel_ to state S_abc_ (or S_off_) (SI Fig. 6). That is,

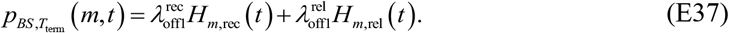

Then, the marginal probability distribution for BS is calculated according to

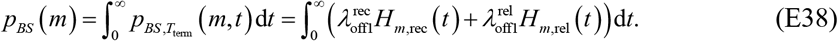

**SI Figure 6.**
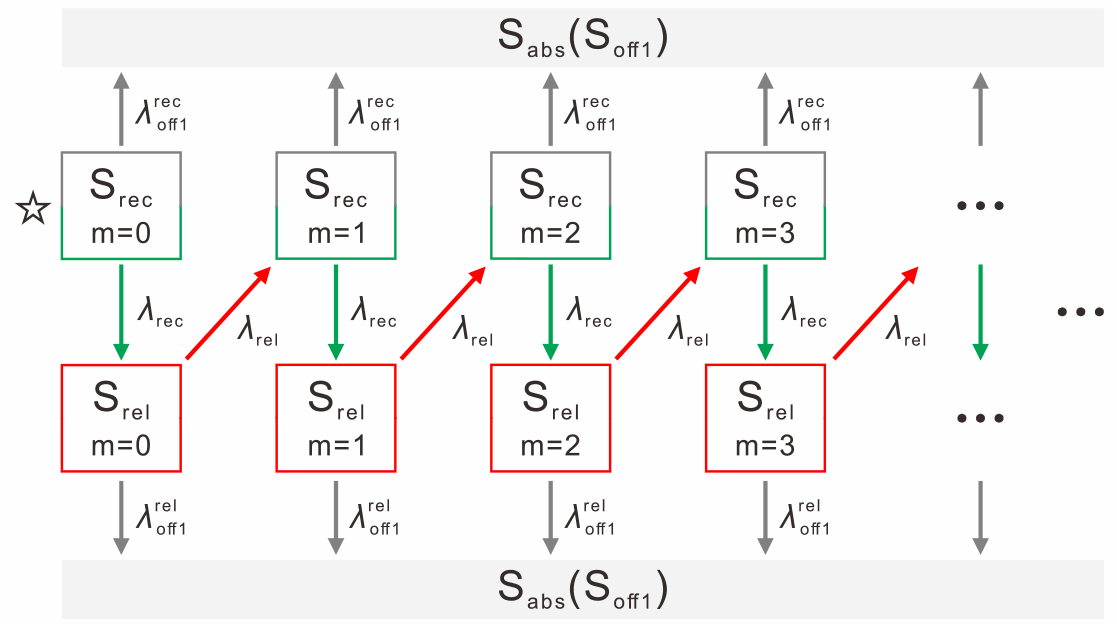
Schematic for the transitions between discrete states and the generation of mRNAs. S_rec_, S_rel_ and S_abs_ are three states in the model. *m* = 0,1,2, ⋯ is the number of mRNAs. *λ_c_*, *c* ∈ {rec, rel, off1} are burst rates. The star in the left top corner is the initial state of a burst.

Denote by *G*(*z*) the generating function for *p_BS_*(*m*). Then based on Eq. (E35) and Eq. (E38), we can obtain

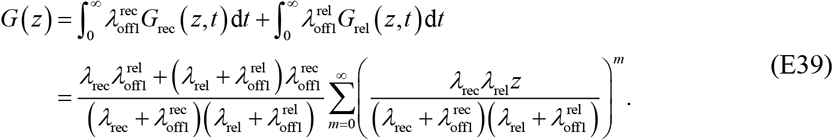

On the other hand, *G*(*z*) can be expanded as 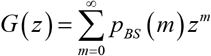. Therefore, by identifying the coefficients of the same powers of *z*, we can see that burst size BS follows the following geometric distribution (referring to SI Fig. 11B)

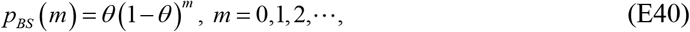

where

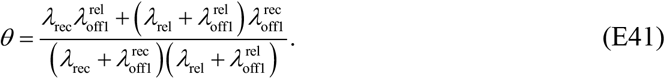

Furthermore, Eq. (E41) can be rewritten as

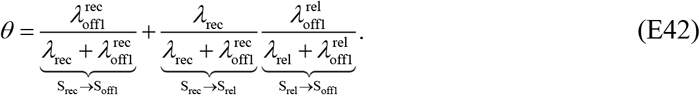

In our model, there are two termination channels whose probability fluxes are expressed perfectly in Eq. (E42). The first term on the right-hand side shows the probability from the state S_rec_ directly back to the burst termination state S_off1_ so that the burst ends. And the second term reflects that the probability flux is first from the state S_rec_ to state S_rel_ and then burst terminates at state S_rel_. Therefore, *θ* is the success probability of burst termination. Besides, 1 – *θ* can be expressed as

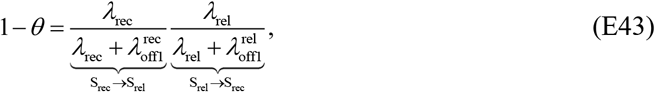

which shows the probability flux from state S_rec_ to state S_rel_ first and then back to the state S_rel_, implying a new mRNA is generated. Thus, 1 – *θ* is the failure probability of burst termination.

In addition, the above geometric distribution of burst size corresponds to the fact that the initial state of each newly generated mRNA is S_rec_ and the number of mRNAs accumulates before the burst termination.

And the MBS is given by

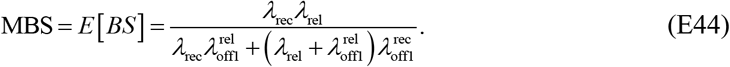

##### 2. Dwell time, cycle time and burst frequency

Next, we derive analytical expressions for dwell time and cycle time. First, we calculate the dwell time in each state. Note that the cycle time is equal to the sum of dwell time in all the states.

Following the above analysis process, we first neglect the dwell time in state S_OFF_ and then compute the dwell time in state S_ON_. Subsequently, the dwell time in the state S_OFF_ can be calculated separately (SI Fig. 7).

**SI Figure 7.**
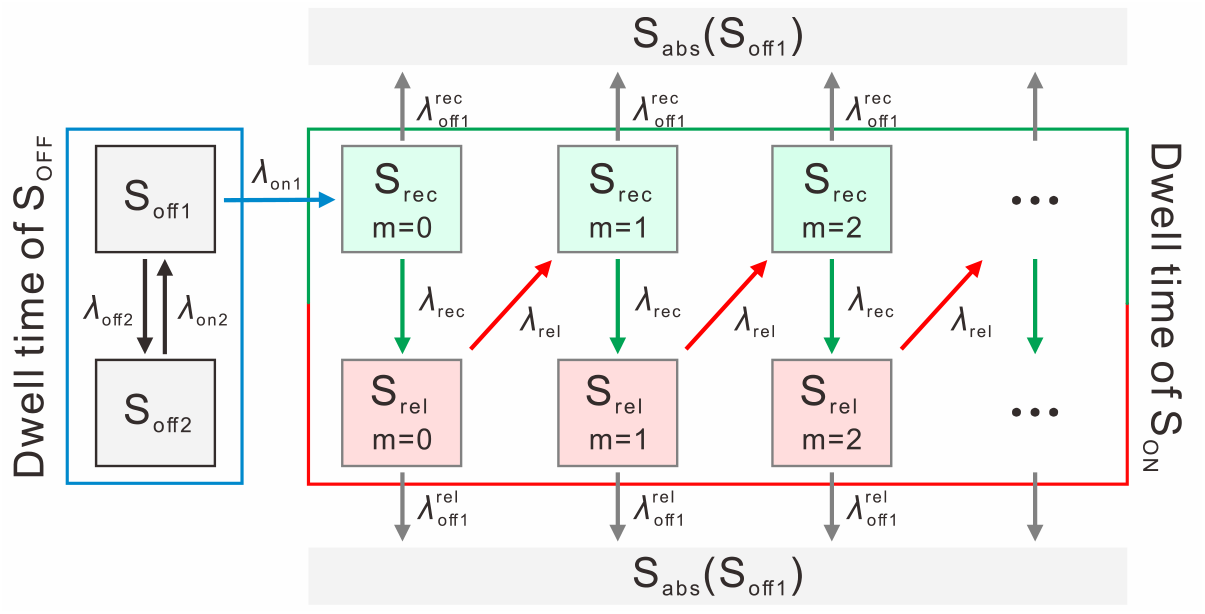
Schematic for the transition between discrete states. The notations have the same meaning as in SI Fig. 6.

Based on the discussion of burst size in the previous subsection, the marginal survival probability for time *T* in Eq. (E23) is

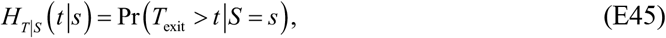

where *T*_exit_ is the exit time from state *s*. For simplicity, we denote *H_T|S_* (*t*|*s*) as *H_s_*(*t*).

By using Eq. (E26) and (E45), and by setting *z* = 1, we have

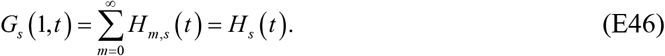

Thus, the survival probability functions at S_rec_ and S_rel_ states equal

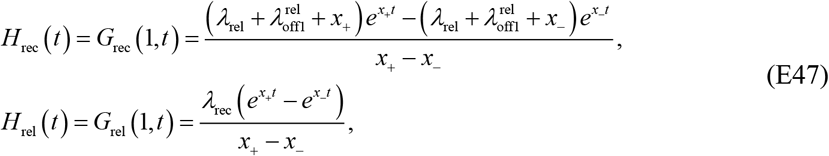

where

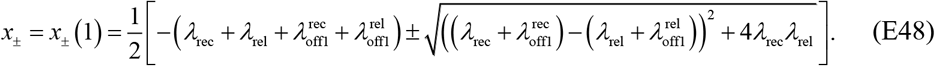

According to the relationship between PDF and survival probability and using Eq. (E47), the dwell time PDFs *p*_rec_(*t*) and *p*_rel_ (*t*) are

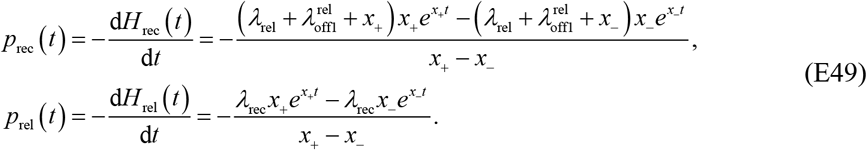

Since Eq. (E47) is the solution of Eq. (E25), the PDF of the total ON state dwell time, *p*_ON_(*t*) (note: the complete expression is *p*_DT|S_ (*t*|S_ON_)) is given by (referring to SI Fig. 11E)

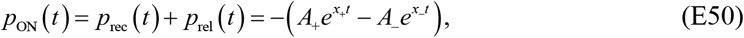

where

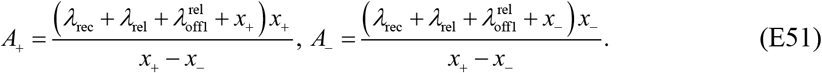

The mean ON dwell time *E*[*DT*_ON_] (note: the complete expression is *E*[*DT*|*S* = S_ON_]) can be obtained and the result is

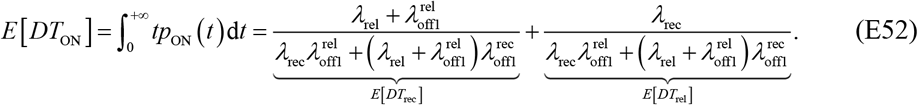

Next, we compute the PDF of dwell time in S_OFF_ state. Owing to the introduction of S_abs_, the S_OFF_ is a state of the flux that the probability only flows out but does not flow in. By using the same method (the derivation process is omitted here), the dwell time PDF *p*_off2_ (*t*) and *p*_off1_ (*t*) are found to be

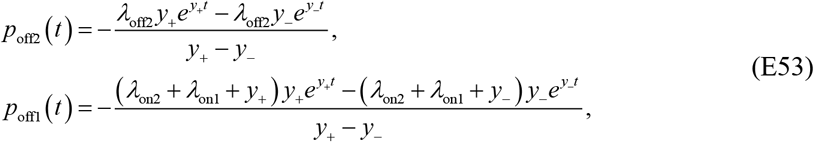

where

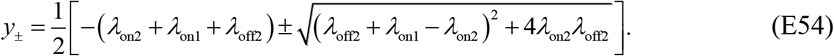

The total OFF state dwell time PDF *p*_OFF_ (*t*) (note: the complete expression is *p_DT|S_* (*t*|S_OFF_)) is given by (referring to SI Fig. 11D)

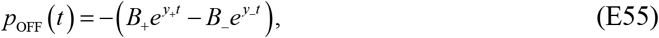

where

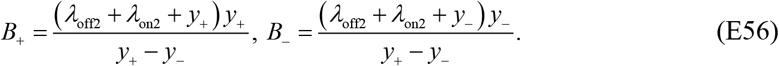

The mean OFF state dwell time *E*[*DT*_OFF_] (note: the complete expression is *E*[*DT|S* = S_OFF_]) is

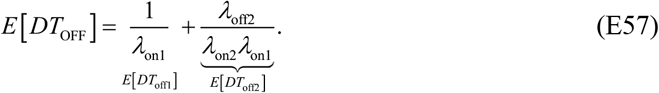

Note that the PDF of cycle time *p_CT_*(*t*) is the convolution of *p*_OFF_(*t*) in Eq. (E55) and *p*_ON_ (*t*) in Eq. (E50). As such, the *p*_CT_(*t*) equals (referring to SI Fig. 11C)

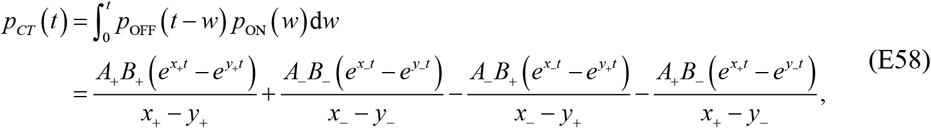

where *x*_±_, *y*_±_, *A*_±_ and *B*_±_ are shown in Eq. (E48), (E51), (E54) and (E56). By complex calculation, the MCT is given by

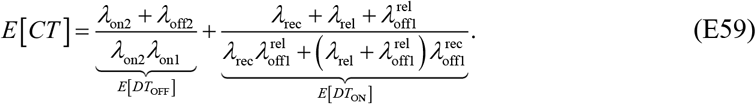

Finally, the BF is given by

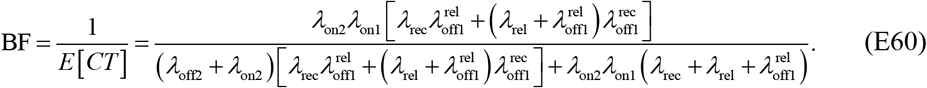

Note that the BF in Eq. (E60) is measured in sec^−1^, meaning how many transcriptional bursts occur in one second.

##### 3. Traveling ratio

Traveling ratio is defined as the Pol II density ratio between gene body and promoter proximity.

**SI Figure 8.**
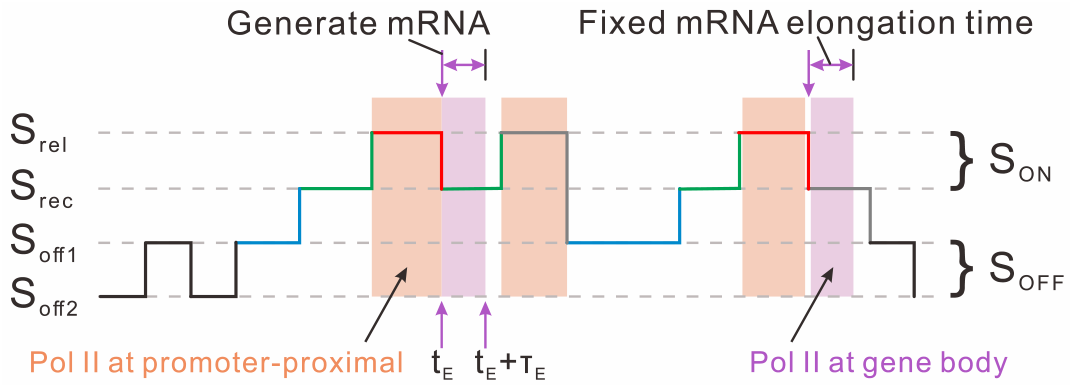
Schematic for describing the location and time of Pol II observable in once simulation. Orange block is the time interval of the Pol II at promoter proximity, and purple block is the time interval of the Pol II in gene body.

First, we calculate the Pol II density in gene body, which is the proportion of elongation time in the whole transcriptional burst or the probability that samples fall in the transcriptional elongation interval (SI Fig. 8, purple block). We can observe the generation of an mRNA at the time *t*_E_ when the state S_rel_ switches to state S_rec_ and the Pol II elongation at a time interval from *t*_E_ to *t*_E_ + *τ*_E_ (where *τ*_E_ is a fixed time lag for mRNA elongation). Obviously, the total elongation time equals the elongation time (*τ*_E_) of one mRNA times the MBS. Thus, the total elongation time of Pol II can be theoretically expressed as *E*[*BS*] · *τ*_E_. The fraction of Pol II in the gene body is the proportion of total elongation time in total cycle time. Based on Eq. (E44) and (E59), and this can be expressed as

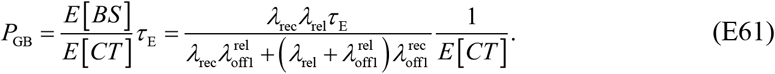

Next, we consider the Pol II density at promoter proximity, which can be viewed as the proportion of dwell-time of state S_rel_ in the whole transcriptional burst (SI Fig. 8, orange block). Hence, based on Eq. (E52) and (E59), the fraction of Pol II at promoter proximity can be expressed as

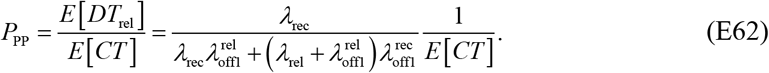

The fraction in Eq. (E62) indicates the probability that the samples are in the state S_rel_. This is the stationary probability in the state S_rel_. Let *P_s_* (*s* ∈{off2, off1, rec, rel}) be the stationary probability of different states. Then, *P_s_* satisfies the following equations

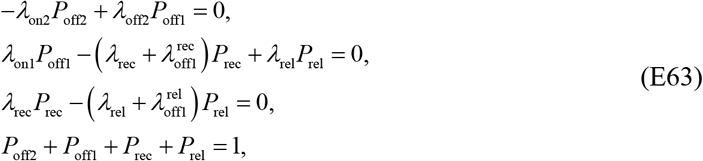

where the last equation holds due to the conservative condition of probability. Solving Eq. (E63), we can obtain *P*_rel_ as expressed in Eq. (E62). That is *P*_PP_ = *P*_rel_.

Finally, the MTR is

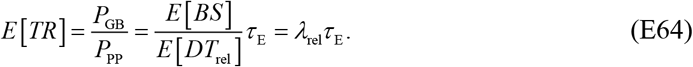

Note that Eq. (E64) is the ratio of two mean values. However, when calculating MTR, we first need to calculate the traveling ratio at each sampling time, and then take the mean value over all sample times. That is 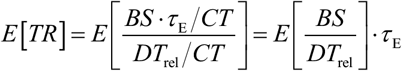. If there are enough samples, we may use approximation 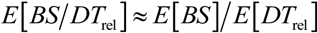. Therefore, the mean traveling ratio is given by Eq. (E64).

### F Power laws for transcriptional bursting kinetics

#### F1 Binary approximation

In the previous section, some rates in Eq. (E6) related to E-P spatial distance are described by a piecewise continuous non-binary Hill function in Eq. (C11). The power *h* of Hill function brings difficulties to the theoretical calculations of MBS, MDT, MCT, and MTR. For this reason, we consider using a simple function to approximate the Hill function. At the simplest level, we may use a deterministic binary rate for transcriptional burst for approximation (SI Fig. 9). That is,

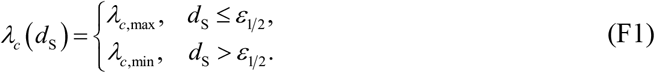

Note that we choose *ε*_1/2_ instead of *ε*_T_ in Eq. (C11) as the threshold.

**SI Figure 9.**
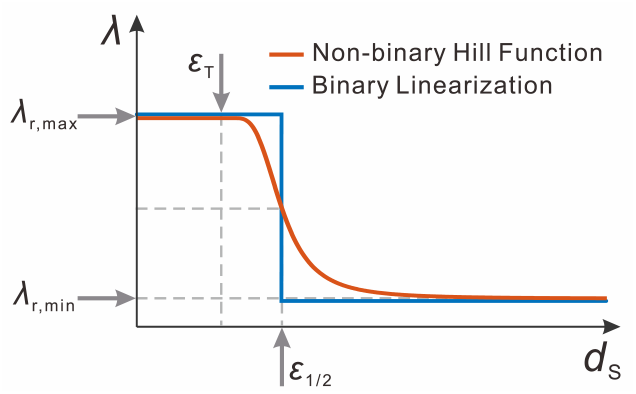
Schematic description of the non-binary Hill function and binary linearization.

For binary linearization, only the maximum rate *λ*_*c*,max_ and the minimum rate *λ*_*c*,min_ are needed. For requirement, we compute ∫*p_DS_* (*d*_S_)d*d*_S_ (referring to SI Fig. 10C), which is the CDF of *p_DS_*(*d*_S_) that can be expressed as

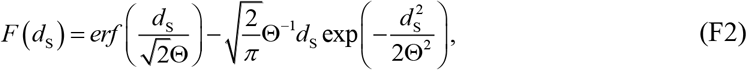

where *erf*(·) is the error function defined as 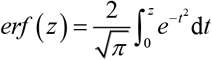.

In the case that fluctuations in *d*_S_ are much faster compared with the rate of transcription, 〈*λ_c_*〉 may be approximated as 〈*λ_c_*〉_*a*_

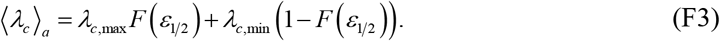

Then, MBS^Fast^ and MCT^Fast^ can be approximated as 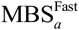 and 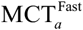, and

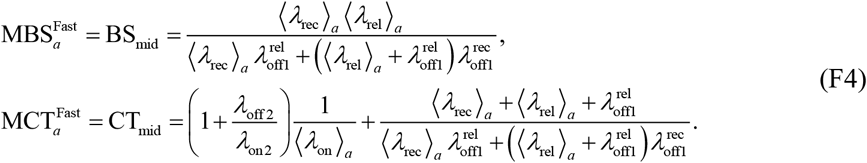

On contrary, if fluctuations in *d*_S_ are much slower, the MBS^Slow^ and MCT^Slow^ can be approximated as 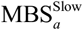 and 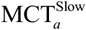, and

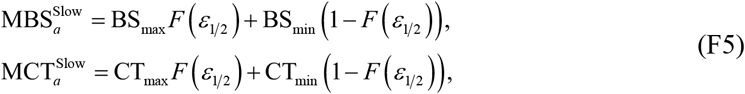

where

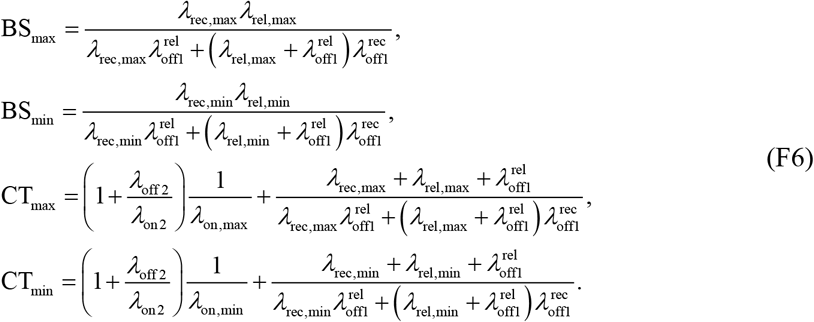

Therefore, based on Eq. (E7) and Eq. (F3) - (F6), the approximations of MBS_*a*_ and MCT_*a*_ can be expressed as (referring to SI Figs. 14D-E and I-J)

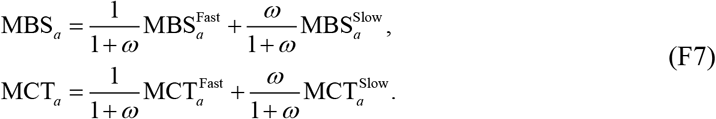

#### F2 Power laws for transcriptional bursting kinetics

E-P communication strength *k*_EP_ and E-P genomic distance *d*_G_ are two key parameters in our model. Note that *F* in Eq. (F2) can be regarded as a function of three independent variables *k*_EP_, *d*_G_ and *d*_S_, and may be written as *F*(*k*_EP_, *d*_G_, *d*_S_). Therefore, MBS and MCT in Eq. (E7) can be also treated as multivariate functions, denoted by MBS(*λ*_*c*,max_, *λ*_*c*,min_, *k*_EP_, *d*_G_, *d*_S_) and MCT (*λ*_*c*,max_, *λ*_*c*,min_, *k*_EP_, *d*_G_, *d*_S_), respectively. In order to show the effects of increasing *k*_EP_ or *d*_G_ on MBS and BF, we calculate derivatives: 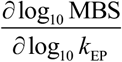 and 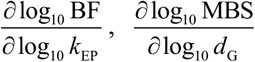 and 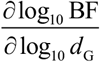.

Using Eq. (F2), we know that 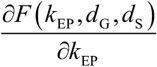 and 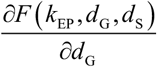 are

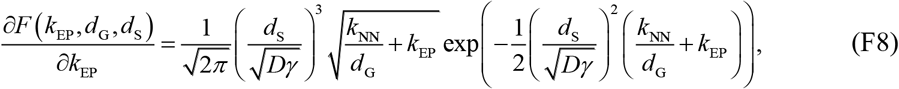

and

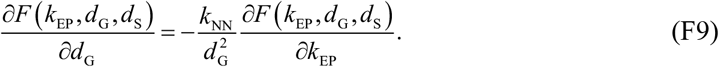

In the following, for the sake of simplicity, we rewrite *F*(*k*_EP_, *d*_G_, *d*_S_)|_*d*_S_=*ε*_1/2__ as *F*(*k*_EP_) (*F*(*d*_G_)) when analyzing the effect of *k*_EP_ (*d*_G_). Note that the following theoretical derivation is for the case of *d*_G_ = 50 (when we consider *k*_EP_) and *k*_EP_ = 0.1 (when we consider *d*_G_). Of course, the corresponding theoretical derivation method is still applicable to cases of other parameter values.

##### 1. Power law in terms of E-P communication strength

The effect of changing *k*_EP_ on MBS can be approximated as

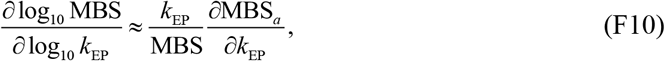

where

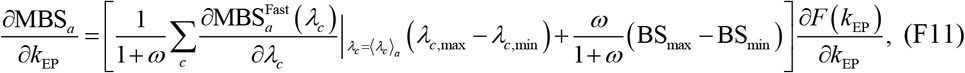

and *c* ∈ {on1,rec,rel}.

Similarly, the effect of changing *k*_EP_ on BF is

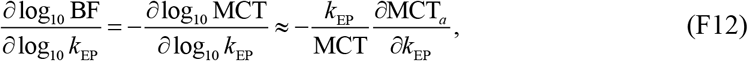

where

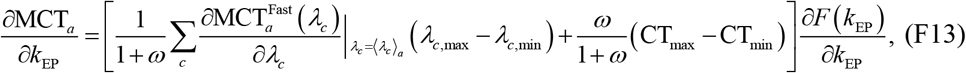

and *c* ∈ {on1,rec,rel}.

Note that both 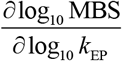 and 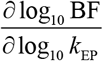 are positive when the parameters are not extreme. They increase for small *k*_EP_ and then quickly decrease to 0 with increasing *k*_EP_. Meanwhile, the value of the derivatives changes little (the maximum values are not more than 0.8, SI Figs. 14A-C), implying that linear approximation is appropriate for log_10_ MBS(log_10_ *k*_EP_) and log_10_BF(log_10_ *k*_EP_) within an appropriate range of *k*_EP_.

Assume that we can obtain distribution *F*(*k*_EP_1__)≈ 1 when *k*_EP_1__ is greater than a pre-given value, and 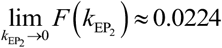 where *d*_G_ = 50. We define

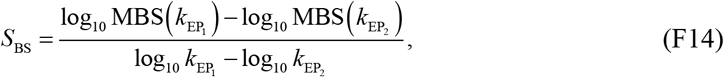

which represents the slope of the line of log_10_ MBS vs log_10_ *k*_EP_. Then, we obtain the following an approximate linear relation:

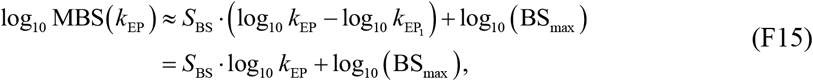

which implies that mean burst size obeys the following power law:

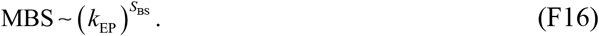

Similarly, if we define

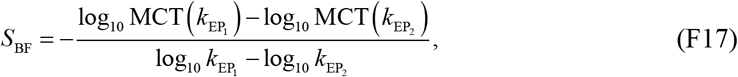

then

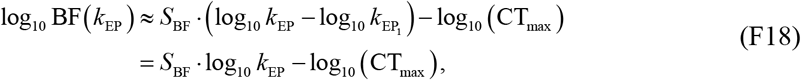

which implies that mean burst frequency obeys the following power law:

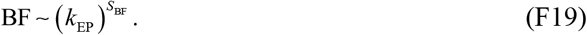

##### 2. Power law in terms of E-P genomic distance

Similarly, we can also compute 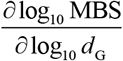 and 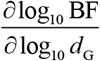, that is,

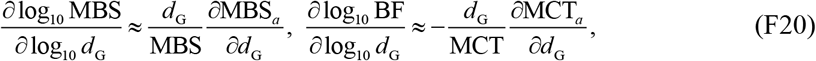

where

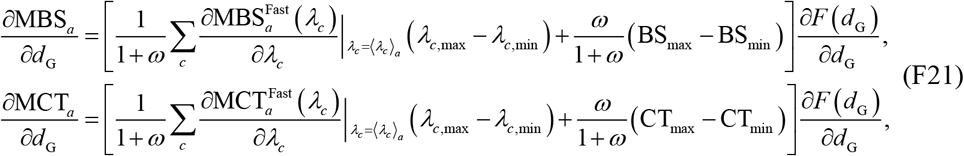

and *c* ∈ {on1,rec,rel}.

Based on Eq. (F9), we know that the monotonicity of 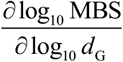 and 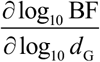 is
opposite that of 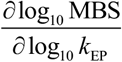 and 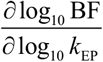, respectively. Moreover, 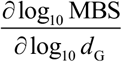 and 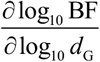 decrease for small *d*_G_ and then quickly increase to 0 with increasing *d*_G_. Meanwhile, the derivatives value changes little (the minimum values are not less than −0.6, SI Figs. 14F-H), implying that linear approximation is appropriate for log_10_ MBS(log_10_ *d*_G_) and log_10_ BF(log_10_ *d*_G_) within an appropriate range of *d*_G_.

Assume that we can obtain distribution *F*(*d*_G_1__ = 1) = 0.9883 where 1 is the minimum of *d*_G_ and *k*_EP_ = 0.1. Numerical results indicate that the turning point for the biphasic responses of burst size and burst frequency to *d*_G_ is different. We denote by 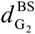 and 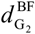 the *d*_G_ values corresponding to the turning points of burst size and burst frequency respectively. And we define

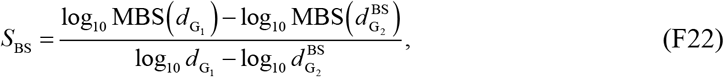

which represents the slope of the line of log_10_ MBS vs log_10_ *k*_EP_. Then, we obtain the following approximate linear relation when 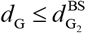

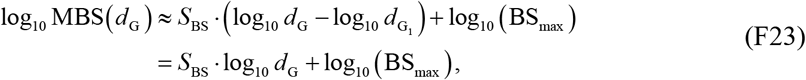

which implies that mean burst size obeys the following power law:

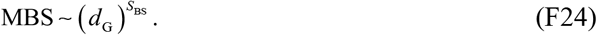

For log_10_ BF, if we define

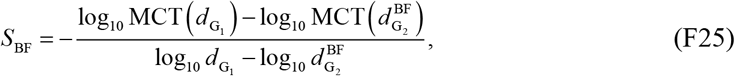

then when 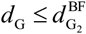, we have

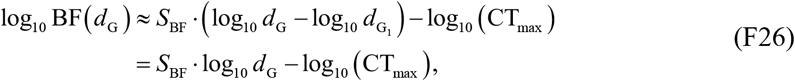

which implies that mean burst frequency obeys the following power law:

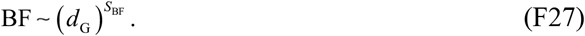

Numerical simulations including analysis of experimental data have verified these power-law behaviors, referring to SI Fig. 16.

### G E-P communication mainly modulates burst frequency

In order to theoretically investigate which of burst size and burst frequency is more affected than the other by E-P communication, we consider the ratio

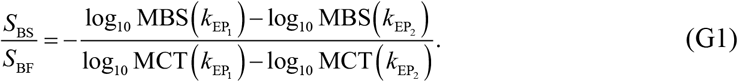

Note that for enhancer deletion, we have *k*_EP_2__ = 0. In this case, the transcriptional burst rates are always the minimums and are independent of E-P spatial distance *d*_S_. Then, Eq. (G1) can be written as

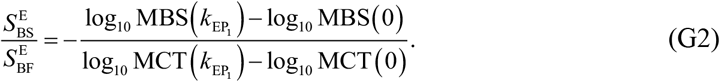

In the main text, we have mapped the high-dimensional parameter space into an experimentally measurable and theoretically computable two-dimensional space. The key is that we take ratios 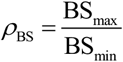 and 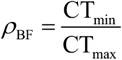 as the coordinates in two-dimensional space, where the involved BS_max_, BS_min_, CT_min_, CT_max_ are defined in Eq. (F6).

### H Calculation of mutual information

Many molecular regulations can impact chromatin conformation. For example, because the hypervariable genes are estrogen-responsive and estrogen induces changes in E-P contacts (90, 91), this leads to genome-wide chromosome conformation changes in response to external stimulations (92). A more detailed study is that by experimentally investigating the endogenous regulation of the estrogen-responsive TFF1 gene, J. Rodriguez et al observed that topologically associated domains appear denser in contacts overall, as reported recently (93). They also observed increases in off-diagonal contact frequency, characteristic of defined enhancer-promoter interactions. However, many estrogen response elements did not show such obvious enrichments between two specific regions, but increased the contact probability with multiple parts of the chromosome and a concomitant loss of contacts with many other regions.

In principle, we may use Shannon entropy to measure the effect of this information propagation on the resulting transcriptional bursting. However, to put this quantitative entropy change in a biological context, here we calculate the mutual information (MI) between downstream random variables *X* ∈{*BS,DT*|*S,CT*} and upstream E-P spatial distance *DS*, denoted by MI(*X*, *DS*), such as MI(*CT*, *DS*). Specifically, we define the mutual information as (referring to SI Fig. 18)

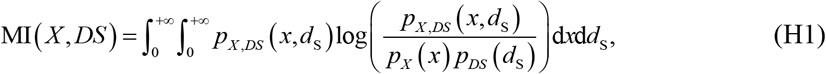

where *p_X_*(*x*) are given in Eq. (E6), *p_DS_*(*d*_S_) are shown in Eq. (E14), and *p_X,DS_*(*x*, *d*_S_) can be calculated according to 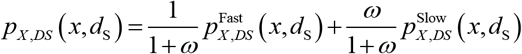, where 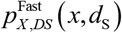 and 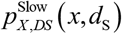 are given in Eq. (E2) and Eq. (E4), respectively. When we calculate MI(*BS*, *DS*), it is necessary to replace the integral symbol in Eq. (H1) with the summation symbol since *BS* is a discrete random variable.

We find the MI(*X*, *DS*) value is in general small. This may be because of the following three reasons. First, our model only considers one E-P pair, but in the real organisms, multiple pairs can be formed. And multiple genes or multipair E-P communication generally transduce more information than a single E-P communication (94). Second, multiple output distributions such as burst size distribution and cycle time distribution make it possible to separately measure the amount of the information transduced from the E-P communication signal to outcome distributions. This would lead to smaller MI(*X*, *DS*) than calculating the mutual information between the input distribution and the gene expression distribution that integrates BS’s and CT’s information. The third possible reason is that the timescale difference between the upstream and the downstream influences the size of mutual information, e.g., if the upstream motion is much faster than the downstream but the information arriving at the downstream is the average, the upstream and the downstream are independent and the mutual information between them is zero. However, the smaller mutual information does not mean that E-P communication is meaningless but may reflect the robustness of E-P communication.

### I Biologically reasonable setting of model parameter values

We set the values of our model parameters mainly based on experimental results on *Drosophila*.

**SI Table 2:** List of model parameter values

| Param | Description | Value | Ref |
| --- | --- | --- | --- |
| Chromatin motion |  |  |  |
| $\Delta t$ | Time step | 0.01s | (28, 29) |
| $k_B$ | Boltzmann constant | $1.38 \times 10^{-23} JK^{-1}$ | --- |
| $T$ | Room temperature | 300K | --- |
| $D$ | Monomer diffusion coefficient | $4 \times 10^{-4} \mu m^2 s^{-1}$ | (95) |
| $\gamma$ | Friction coefficient | $10 pNs \mu m^{-1}$ | $D\gamma = k_B T$ |
| $b$ | Standard deviation distance between monomers | $0.1 \mu m$ | (28) |
| $k_{NN}$ | Spring constant between adjacent monomers | $1 pN \mu m^{-1}$ | $k_{NN} = 3k_B T / b^2$ |
| $k_{EP}$ | E-P interaction coefficient | free | --- |
| Transcription |  |  |  |
| $\lambda_{on2}$ | Chromatin opening rate | $0.002 s^{-1}$ | (56) |
| $\lambda_{off2}$ | Chromatin closure rate | $0.006 s^{-1}$ | (56) |
| $\lambda_{on1}$ | Transcriptional initiation rate (from min to max) | $0.002 \sim 0.020 s^{-1}$ | (96) |
| $\lambda_{rec}$ | Pol II recruitment rate (from min to max) | $0.009 \sim 0.024 s^{-1}$ | (97) |
| $\lambda_{rel}$ | Pol II pause release rate (from min to max) | $0.008 \sim 0.025 s^{-1}$ | (97) |
| $\lambda_{off1}^{rec}$ | Transcriptional termination rate (channel 1) | $0.009 s^{-1}$ | (38, 96) |
| $\lambda_{off1}^{rel}$ | Transcriptional termination rate (channel 2) | $0.002 s^{-1}$ | (38, 96) |
| $h$ | Hill coefficient | 3 | (98) |
| $\varepsilon_{1/2}$ | Half-value distance of burst rates | $0.2 \mu m$ | (37) |
| $\varepsilon_T$ | E-P encounter distance (distance threshold) | $0.1 \mu m$ | $b$ |
| $r_E$ | Transcriptional elongation rate | $30 bps^{-1}$ | (99-102) |
| $L_E$ | Elongation length (gene body length) | $3000 bp$ | (103) |
| $\tau_E$ | Transcriptional elongation time | 100s | $\tau_E = L_E / r_E$ |
| Other parameters |  |  |  |
| $l$ | Nucleosome DNA length | $200 bp$ | (104) |
| $L$ | Genome length | $20 kbp$ | (105) |
| $N$ | Number of monomers | 100 | $N = L/l$ |

*Drosophila*, one of the intensively studied organisms in the biological field, serves as a test system to explore cellular processes including transcriptional bursting. To investigate the power of our model, it is needed to discuss how the model parameters to fit experimental conditions.

#### 1. Enhancer and promoter lengths

Enhancers and promoters are regulatory elements in transcriptional activation and bursting, and their lengths are highly variable in different organisms or genes.

The length of an enhancer can vary from about 50 bp to 1.5 kbp (106) and typically, the metazoan enhancer is ~500 bp (107). For example, in *Drosophila* embryo, the *rhomboid* (*rho*) enhancer is 300 bp in length, the *brinker* (*brk*) and *vein* (*vn*) enhancers are 500 bp in length (107) and the enhancer of *lab* gene contains a 550 bp fragment (108). Each enhancer contains specific motifs and multiple binding sites for TFs, and these motifs assemble within a ~300 bp core domain (107). Therefore, we stipulate that the enhancer length is 300 bp in simulations.

Promoters can be divided into peaked and broad promoters based on their widths. And promoter length is also highly variable. In *Drosophila*, the median width of broad promoters is 162 bp (109). An experimental program obtained 422 kbp promoter sequences from 2424 genes in *Drosophila* (110), and then we can calculate that one promoter is about 174 bp in length. Thus, we assume the promoter length is 170 bp in simulations.

The length of DNA around a nucleosome is about 167 bp, which is approximate to that of a promoter. In contrast, the length of the linker DNA is more variable, ranging from 10 bp to 90 bp (104, 111). Thus, the length of DNA in one nucleosome plus the length of the linker DNA at both sides of the nucleosome is approximately equal to that of the enhancer. Overall, we can use one nucleosome indicates an enhancer or a promoter.

#### 2. E-P genomic distance and the number of monomers

Along the linear genome, the distances between the enhancers and their cognate promoters vary greatly. In *Drosophila*, this distance can be more than 100 kbp. The median distance is 10 kbp and the most possible distance of an E-P pair is 5 kbp (105). The distances from more than 70% of enhancers to target transcriptional start sites do not exceed 20 kbp. Therefore, we set the total genome length *L* as 20 kbp in simulations. And the E-P genomic distance is set to be 10 kbp by default. When considering the effect of E-P genomic distance on transcriptional bursting, we let the E-P genomic distance increase from 1 kbp to 20 kbp. In simulations, we assume that one monomer represents a nucleosome (~200 bp). Thus, the number of monomers *N* is 100.

#### 3. Spring coefficient between adjacent monomers and monomer diffusion coefficient

The Boltzmann constant equals *k*_B_ = 1.38× 10^−23^ *J/K* and we set temperature *T* as 300*K* (room temperature). Thus, *k_B_T* ≈ 4×10^−3^ *pN μm*. The standard deviation of the distance between adjacent monomers is at the micron level (28). Therefore, we posit *b* = 0.1*μm*. According to the relation *k*_NN_ = *dk_B_T/b*^2^, we can compute the spring coefficient for successive monomers, i.e., *k*_NN_ = 1*pN/μm*.

Owing to the Einstein relation *Dγ* = *k_B_T* and the fact that the E-P spatial distance PDF is affected by the product of *D* and *γ* (see Eq. (E14)), we may not care about the respective values of *D* and *γ*, but set their product as *k_B_T* ≈ 4×10^−3^ *pN μm*. We set the monomer diffusion coefficient of DNA as *D* = 4×10^−4^ *μm*^2^/*s* and thus the friction coefficient is *γ* = 10*pNs/μm*. Of course, for *D* and *γ*, we can set different values, but *Dγ* = *k_B_T* must be guaranteed. It is worth noting that the size of *γ* affects the timescale of upstream chromatin motion.

#### 4. E-P communication strength

In *Drosophila* embryos (37), the difference between the E-P distances of different experiments can reach up to 1*μm*. The E-P interaction coefficient *k*_EP_ may change in the range of 0–1 *pN/μm*, so we can study the effects of distinct *k*_EP_ on bursting kinetics. Note that *k*_EP_ = 0 means that there is no communication between enhancer and promoter. Also, it is unnecessary to consider the case of larger *k*_EP_ (> 1*pN/μm*) because the bursting dynamics do not change with the increase *k*_EP_ when *k*_EP_ > 1*pN/μm*. The *k*_EP_ is set to be *k*_EP_ = 0.1 *pN/μm* by default in the main text.

#### 5. Hill function

The Hill coefficient *H* measures ultra-sensitivity. In biology, the *H* usually changes from 2 to 5 (98, 112). We fix *h* = 3 for the studies in the main text. The E-P encounter distance is *ε_T_*= 0.1*μm*, which equals to the standard deviation of the distance between adjacent monomers *b* and approximately equals to 0.12*μm* in (113).

E-P communication or E-P proximity is necessary for transcriptional bursting. The E-P distance is about 0.2 ~ 0.4*μm* in bursting (25, 37, 72, 82). When E-P spatial distance is larger than 0.4*μm*, the burst rates tend to the minimum. Considering this, we assume the half-value distance of burst rates is *ε*_1/2_ = 0.2*μm*, and thus the rates drop to the minimum around 0.4*μm*.

#### 6. Gene states switching rates

In *Drosophila*, the burst duration is about 10-20 minutes (14, 96, 114, 115). The sm-FISH has shown that in *Drosophila* gap genes, the mRNA is transcribed with the average residence time ≈ 3 min and the gene is not expressed with a relatively longer lifetime.

In Section E, we analytically showed that the OFF-state dwell time is 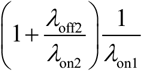. For simplicity, we assume that the relationship between *λ*_off2_ and *λ*_on2_ is multiple (56), e.g., *λ*_off2_ = 3*λ*_on2_. Thus, the OFF-state dwell time is 4/*λ*_on1_. We can assume that the fluctuating range of the transcriptional initial rate is from *λ*_on1,min_ = 0002*s*^−1^ to *λ*_on1,max_ = 0.020*s*^−1^. Due to the deep inactive state S_off2_, the transition rates between S_off2_ and S_off1_ are smaller than those between S_off1_ and S_rec_. We set *λ*_off2_ = 0.006*s*^−1^ and *λ*_on2_ = 0.002*s*^−1^.

If we set 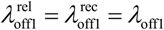, the ON-state dwell time can be theoretically reduced to 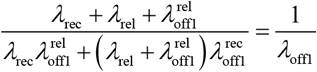. We set the value of *λ*_off1_ as 0.006*s*^−1^. Because the paused Pol II is more stable (38, 51), we suppose 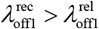 and set 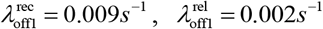.

The promoter-proximal pausing timescale is about a minute level (97). We assume that the fluctuating range of the Pol II pause release rate is from *λ*_rel,min_ = 0.008*s*^−1^ to *λ*_rel,max_ = 0.025*s*^−1^. In addition, we assume that the fluctuating range of the Pol II recruitment rate is from *λ*_rec,min_ = 0.009*s*^−1^ to *λ*_rec,max_ = 0.024*s*^−1^

#### 7. Transcriptional elongation

DNAs are transcribed at a speed of about 10–100 *bp/s* (116) across organisms and conditions. Many methods have measured that in *Drosophila*, the elongation rate is 1.1–1.5 kb/min, i.e., ~ 25 *bp/s* (99–102). In addition, recent studies have suggested that in the *Drosophila* embryo, the Pol II elongation rate is ~40*bp/s* (117), approximately twice as the previously estimated value. These facts indicate that the rate of elongation is variable in different developmental stages. For simplicity, we consider that the Pol II elongation rate *r*_E_ is about 30 *bp/s*.

In *Drosophila*, the size of the averagely predicted mRNA transcripts is 3058 bp (103). We let the gene body’s length *L*_E_ be roughly 3000 bp. Therefore, the transcriptional elongation time *τ*_E_ is about 100*s*. Note that the *L*_E_ here is different from genome length *L*. *L*_E_ is the length of the gene to be transcribed after the pause release of the Pol II, which is independent of *L* in simulations.

### J Experimental confirmations and model predictions

Our model studies verify multiple experimental results and make some experimentally testable predictions. Here we summarize and discuss a few of the predictions in details.

#### J1 Upstream E-P Maxwell-Boltzmann distribution

##### 1. E-P 3D spatial distance and its Maxwell-Boltzmann distribution

In the main text, we set *k*_EP_ = 0.1 and *d*_G_ = 50, corresponding to Θ = 0.183. Both theoretical derivations and numerical simulations verify that the E-P distance follows a Maxwell-Boltzmann distribution with the mean of −0.3*μm*, which nearly equals to 0.34*μm* of PP7 reporter gene in *Drosophila* (37) and Sox2 gene in ESCs (72). Nevertheless, the spatial distance between cis-regulatory elements and *eve* locus in *Drosophila* (82) is notably smaller (~0.2*μm*) than that in our model. This difference may come from the fact that *eve* locus contains multiple enhancers, resulting in a compact globular structure with a spatial scale ~ 0.2*μm*.

Importantly, the work of (72) clearly showed the histogram of 3D separation distance between Sox2 and its essential enhancer in living ESCs. Upon Maxwell-Boltzmann distribution fitting, the E-P distances fit well with the parameter Θ of 0.195, suggesting the validity of theoretical distribution and parameter selection (SI Fig. 10B).

##### 2. E-P encounter probability and effective density

Based on the theoretically-derived Maxwell-Boltzmann distribution, we can calculate the relationship between encounter probability and mean physical distance (〈*d*_S_〉). The log-log plot (SI Fig. 10E, left) shows that encounter probability is monotonically decreasing with the spatial distance following a power-law behavior, in accordance with the experimental results for consecutive and nonconsecutive TAD borders in *Drosophila* (113). In fact, in that reference, the interaction between TAD borders was built through some proteins (such as cohesin), similar to the E-P communication. Furthermore, the encounter probability is decreasing with E-P genomic distance and even unchanged for larger *d*_G_. For small *d*_G_, the data represent a power-law fitting (SI Fig. 10E, right). This finding is almost consistent to the result in (113).

On the other hand, we draw the effective density or concentration for variable radius *R* (SI Fig. 10F). There is a plateau below *R* ~ 0.2*μm*, which is supportive to the experimental finding of (82). Besides, the additional genomic separation between E5 and the promoter (82), corresponding to the increment of *d*_G_, results in an overall decrease in effective density (SI Fig. 10F, right).

#### J2 Power-laws for transcriptional bursting kinetics

##### 1. E-P communication strength

E-P communication strength *k*_EP_ is a key parameter in the regulation of transcriptional bursting. In physics, *k*_EP_ is merely a quantity used to measure the E-P communication intensity. The *k*_EP_ can represent distinctive enhancers or external stimuli that can influence E-P communication. For example, it was found that transcriptional burst size and traveling ratio increase during estradiol-mediated erythroid maturation (40, 53). These experimental phenomena are consistent with our model prediction that the increase *k*_EP_ can boost both burst size and burst frequency.

In fact, we have found that the log_10_MBS and log_10_BF respectively obey an approximately linear relation with respect to log_10_*k*_EP_. In the erythroid maturation experiment (40), if we use the erythroid maturation time after adding the hormone to reflect the E-P communication strength *k*_EP_, burst size of *β*-globin gene can fit well with a power-law function. In the work of (39), the researchers studied the effect of five distinctive enhancers on transcriptional bursting in living *Drosophila* embryos. We find that the burst size and burst frequency as well as transcriptional output (SI Figs. 16A-B) exhibit power-law-like behaviors when the enhancer’s strength increases in an equidistant interval on a logarithmic scale. In another example, higher average c-Fos mRNA levels were induced with increasing zinc concentration (118). If taking the experimental data in the logarithmic sense, we find a linear relation between logarithmic mean mRNA expression and logarithmic *k*_EP_ representing zinc concentration (SI Fig. 16C).

##### 2. E-P genomic distance

The E-P genomic distance *d*_G_ is also a key parameter in controlling transcriptional bursting. A support for this is clearly shown in the pivotal work of (39, 119), wherein the authors monitored the expression level of distinctive E-P distance. The burst size and frequency reduce when the E-P genomic distance becomes larger in *sna* shadow enhancer and synthetic MS2 reporter gene in *Drosophila*. These experimental phenomena are in accord with the results predicted by our model.

It was theoretically shown that for a smaller E-P genomic distance, the log_10_MBS and log_10_BF respectively obey an approximately linear relation with respect to log_10_*d*_G_. In the work of (120), the investigators plotted the average mRNA expression level in MSCs, which approximately equals to the burst size multiplied by the burst frequency, as a function of the contact probability, which can be regarded as negative linearity over a small distance. Interestingly, the logarithmic mean mRNA expression level follows a linear relation with the logarithmic contact probability, which is consist with the prediction of our model (SI Fig. S16D). Also, the mean mRNA expression level monotonically decreased with increasing E-P genomic distance in MSCs, implying a power-law behavior (120). Importantly, the transcription level may not drop to zero but remain at a low level for larger *d_G_* (120).

#### J3 E-P communication mainly regulates burst frequency

In Section G, by theoretically calculating the values of *S*_BS_/*S*_BF_ and 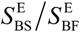, we have shown that E-P communication mainly regulates burst frequency instead of burst size.

The early studies about cell-to-cell variability indicated that the enhancer regulates the probability of transcriptional bursting, rather than the transcriptional level (121, 122). The core promoter determines burst size but enhancers increase bursting frequency (75). Recent experiments such as *β*-globin enhancer in the murine erythroblast cell during erythroid maturation (40), different developmental enhancers in *Drosophila* embryos (39), and transgene experiment in mouse embryonic stem cells (120), also verified this phenomenon.

Besides, transcriptome-wide inference in mouse embryonic stem cells and primary mouse fibroblasts has shown the important role of E-P communication in controlling burst frequency (18, 123).

### K Supplementary figures

**SI Figure 10.**
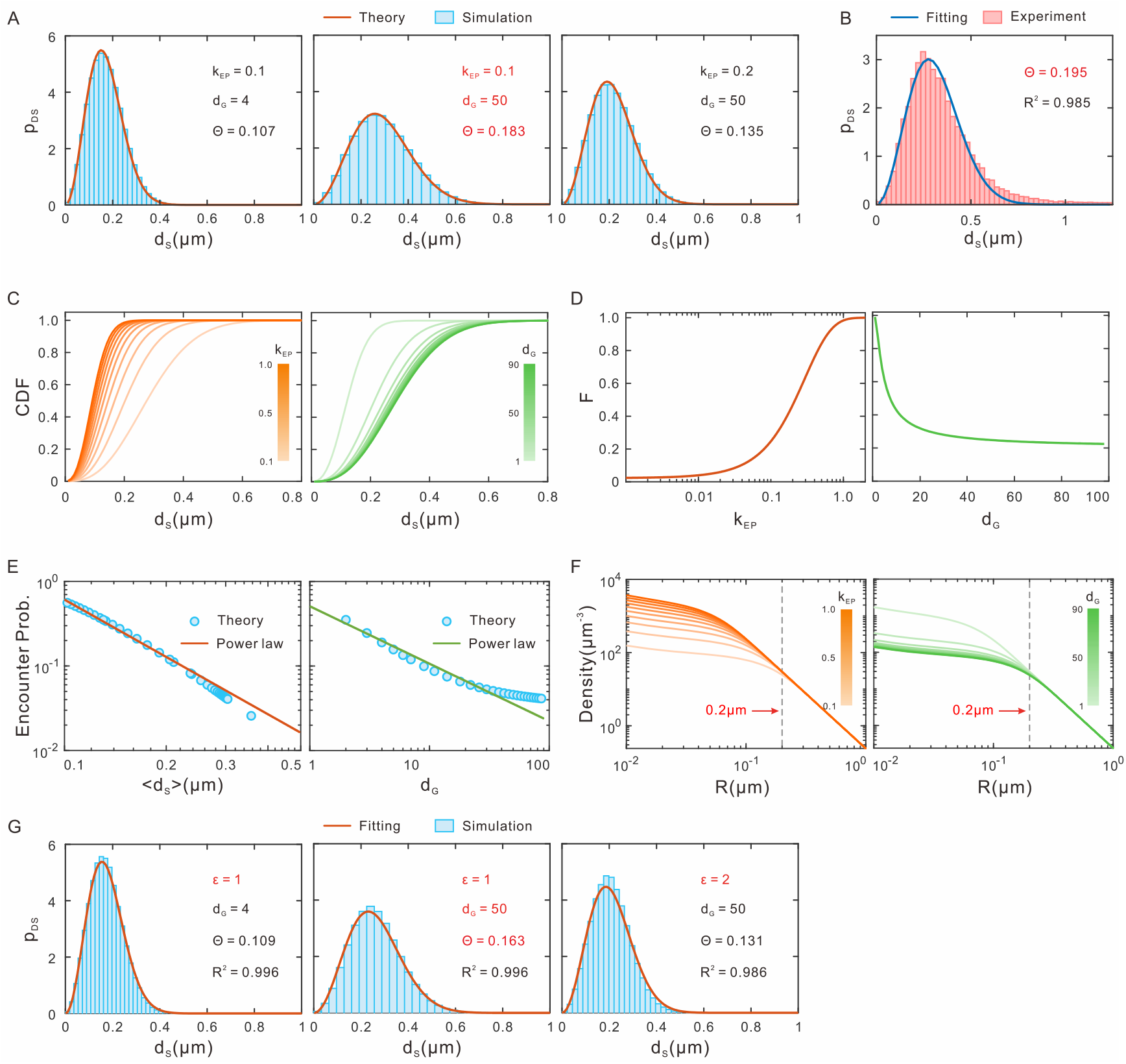
Distributions and properties of upstream E-P spatial distance. (A) Influence of *k*_EP_ and *d*_G_ on the distribution of E-P spatial distance (Maxwell-Boltzmann distribution, Eq. (E14) and (E15)) where red lines represent theoretical results, histograms represent numerical results. (B) Experimental measurement of 3D spatial distance between Sox2 and the SCR enhancer in living ESCs (72). The blue line is obtained by fitting the Maxwell-Boltzmann distribution. (C) Cumulative density function (CDF) of E-P spatial distance (Eq. (F2)) for different *k*_EP_ (left, fixed *d*_G_ = 50) and *d*_G_ (right, fixed *k*_EP_ = 0.1). (D) *F*(*k*_EP_, *d*_G_, *d*_S_) as a function of *k*_EP_ (left, fixed *d*_G_ = 50 and *d*_S_ =*ε*_1/2_) and *d*_G_ (right, fixed *k*_EP_ = 0.1 and *d*_S_ = *ε*_1/2_). (E) Encounter probability as a function of mean spatial distance 〈*d*_S_〉 (left) and *d*_G_ (right). Blue points are obtained by theoretical calculation, whereas solid lines represent power-law fitting. Only the points with *d*_G_ < 60 are fitted by power-law. (F) Effective density or concentration of E-P communication as a function of the averaging volume (the sphere of radius *R*) for different *k*_EP_ (left, fixed *d*_G_ = 50) and *d*_G_ (right, fixed *k*_EP_ = 0.1). (G) Influence of Lennard-Jones (LJ) potential well depth *ε* and *d*_G_ on probability density function of *d*_S_ where histograms represent numerical results and red lines represent fitting results by Maxwell-Boltzmann distribution (parameter is Θ). Other parameters are the default values as in SI Table 2.

**SI Figure 11.**
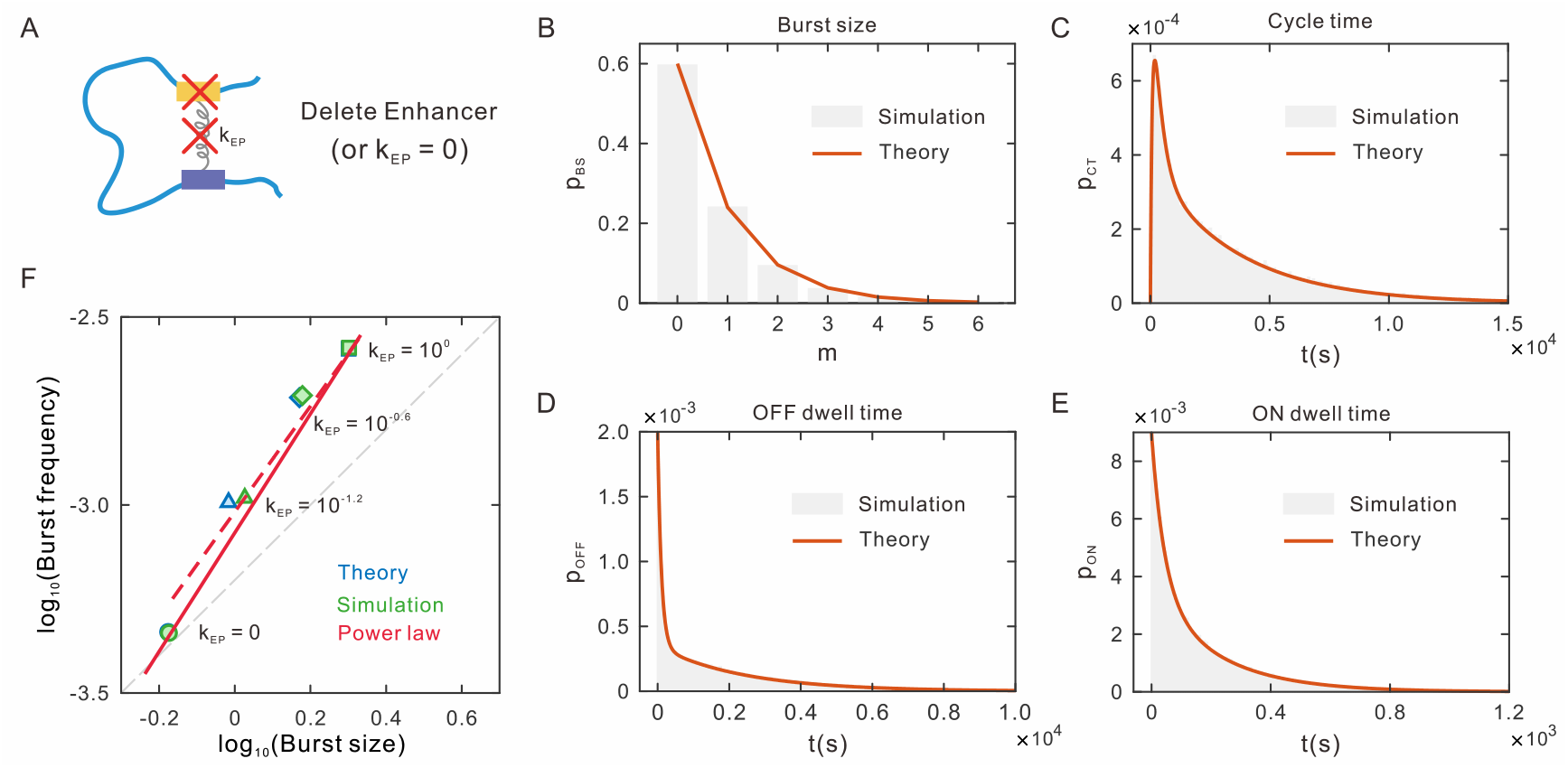
Effect of enhancer deletion on transcriptional bursting, corresponding to the case where only downstream transcriptional bursting is considered. (A) Schematic for enhancer deletion (i.e., *k*_EP_ = 0). In this case, the transcriptional burst rates are always minimums. (B-E) Probability distributions of burst size (B), cycle time (C), OFF dwell time (D) and ON dwell time (E) when *k*_EP_ = 0. The red solid lines are drawn based on Eq. (E40) for (B), Eq. (E58) for (C), Eq. (E55) for (D) and Eq. (E50) for (E), representing theoretical results. The histograms represent the results obtained by numerical simulations. (F) Influences of different values of *k*_EP_ on log_10_ (MBS) and log_10_ (BF). The red dashed line represents the approximate linear relationship between log_10_ (MBS) and log_10_ (BF) without considering enhancer deletion. And this line’s slope, obtained by calculating the ratio of slope (Eq. (G1)) between log_10_ (MBS) and log_10_ (*k*_EP_) (Eq. (F14), *S*_BS_ = 0.23) over the slope between log_10_(BF) and log_10_(*k*_EP_) (Eq. (F17), *S*_BS_ = 0.33), is 1.40. The red solid line represents the approximate linear relationship between log_10_(MBS) and log_10_(BF) when *k*_EP_ = 0 and the line slope is 1.56. The dashed gray line is the indicator of the slope equaling 1. Other parameters are the default values as in SI Table 2.

**SI Figure 12.**
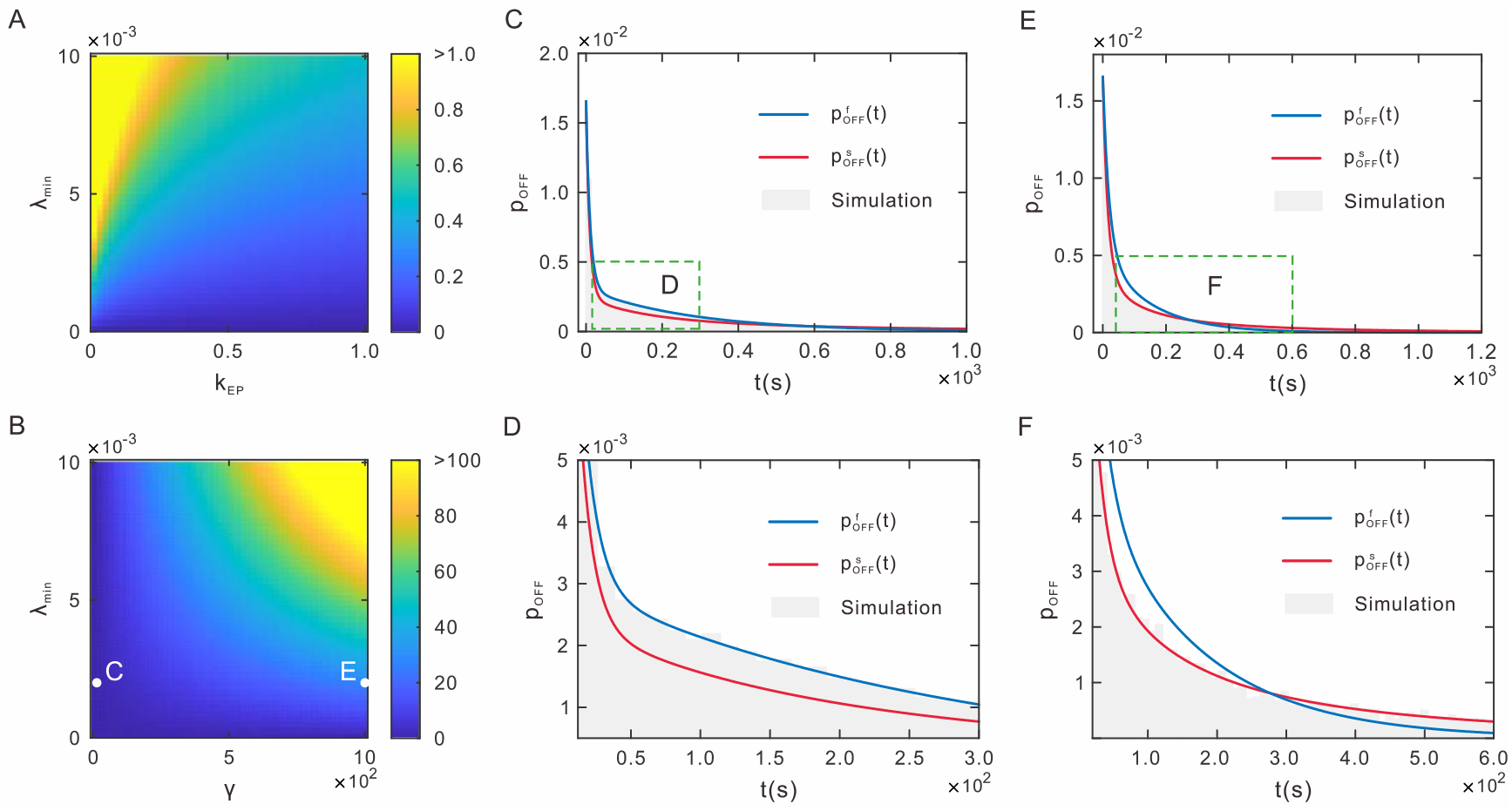
The effect of upstream and downstream timescales on downstream distributions, where OFF distribution is taken as an example for demonstration. (A) The maximum achievable ratio of upstream and downstream timescales *ω* in Eq. (E1) is a function of minimum state switching rates *λ*_min_ and E-P communication strength *k*_EP_. (B) The *ω* is a function of minimum transcriptional rate *λ_min_* and drag coefficient *γ*. (C-F) Blue solid lines representing the OFF PDFs are drawn by Eq. (E3), whereas red solid lines by Eq. (E5). The histograms represent the results obtained by numerical simulations. (C-D) Upstream chromatin fluctuations are much faster compared with the downstream state switching rates. Parameter values are set as *γ* = 0.1, *γ*_on1,min_ = 0.002, *γ*_on1,max_ = 0.05, *λ*_off2_ = 0.06, *γ*_on2_ = 0.02. (E-F) Upstream chromatin fluctuations are slow. Parameter values are set as *γ* = 1000, *λ*_on1,min_ = 0.002, *λ*_on1,max_ = 0.05, *λ*_off2_ = 0.02, *λ*_on2_ = 0.02. Other parameters are the default values as in SI Table 2.

**SI Figure 13.**
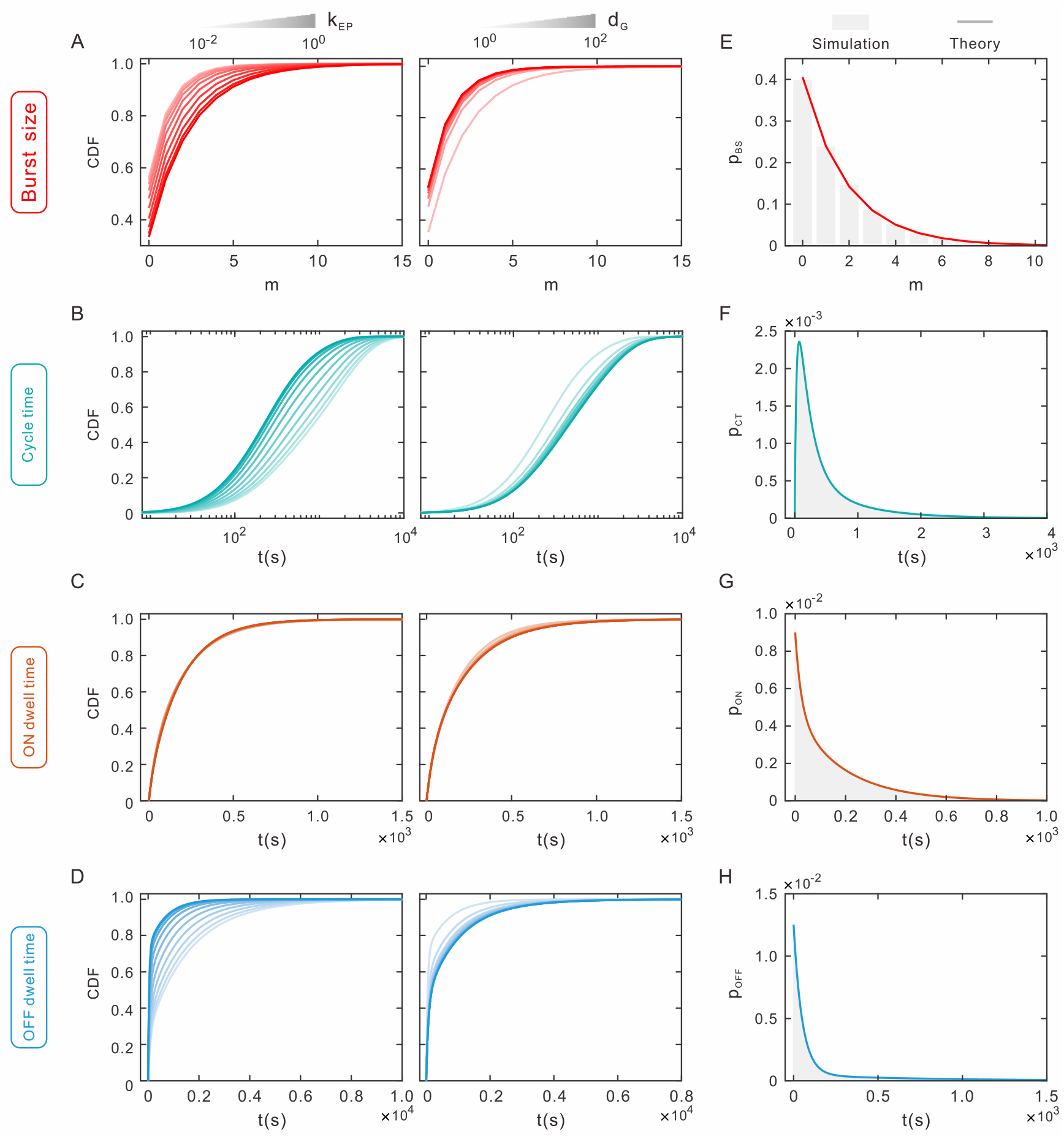
Distributions of burst size, cycle time, ON time and OFF time. (A-D) CDF (The corresponding PDFs or PMF are shown in Eq. (E6)) of burst size (A), cycle time (B), ON time (C) and OFF time (D) for different *k*_EP_ (left, fixed *d*_G_ = 50) and *d*_G_ (right, fixed *k*_EP_ = 0.1). (E-H) Probability distributions of burst size (E), cycle time (F), OFF dwell time (G) and ON dwell time (H) where numerical simulation results correspond to grey histograms whereas analytical results to solid lines. Parameter values in (E-H) are set as *k*_EP_ = 10^−0.6^, *d*_G_ = 50. Other parameters are the default values as in SI Table 2.

**SI Figure 14.**
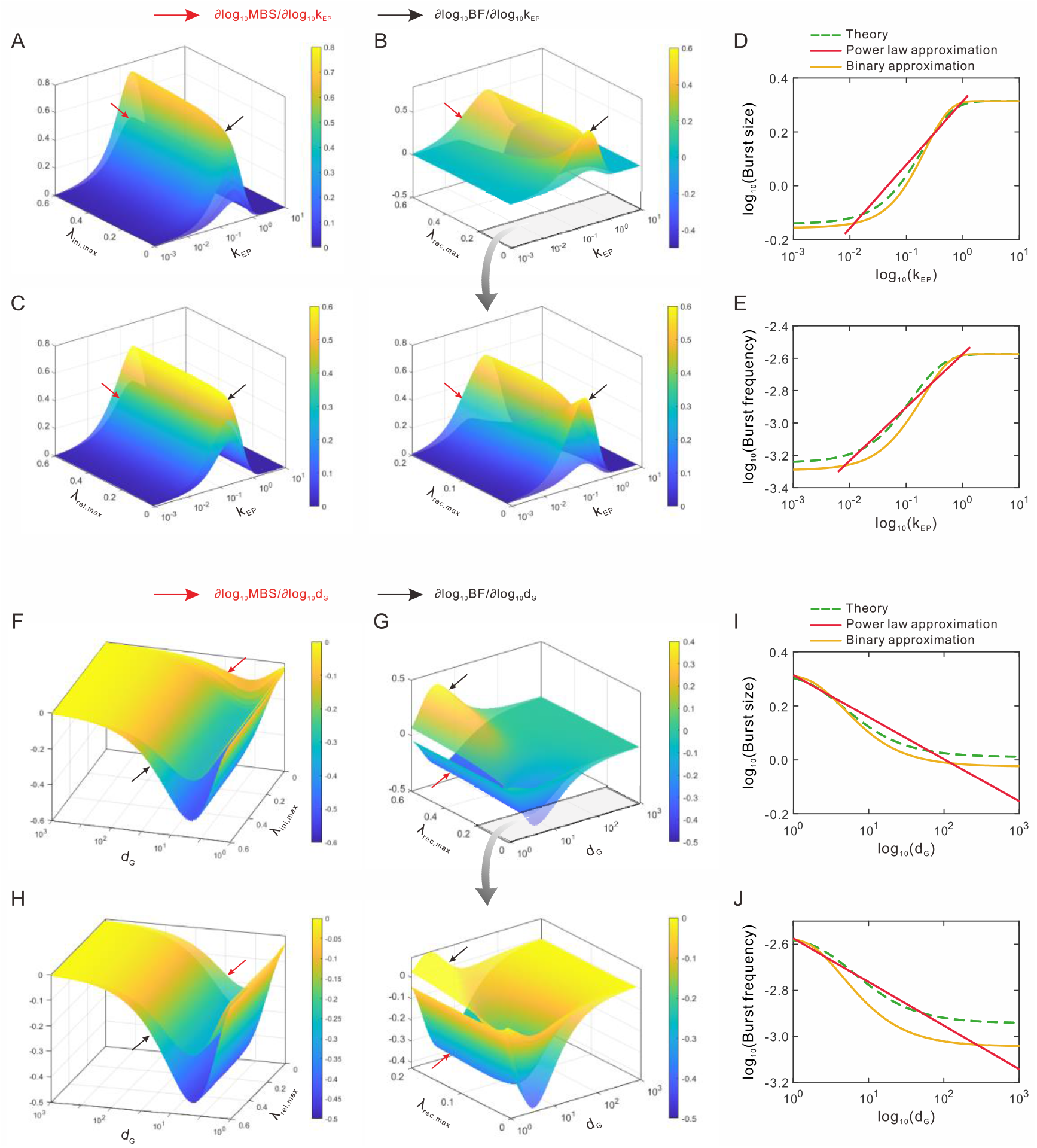
Rationality and validity analysis of power-law approximations for transcriptional bursting kinetics. (A-C) Derivatives 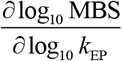 (Eq. (F10)) and 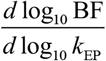 (Eq. (F12)) as the functions of parameters *k*_EP_ and *λ*_ini,max_ (A), *λ*_rec,max_ (B), *λ*_rel,max_ (C). The bottom of (B) is a magnified display of a portion of the top of (B). (D-E) log_10_ MBS and log_10_BF as the functions of log_10_(*k*_EP_), where green dashed lines represent theoretical prediction (Eq. (E7)), orange solid lines represent binary approximation (Eq. (E7)), and red solid lines represent power-law approximation (Eq. (F15)). (F-H) Derivatives 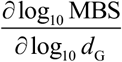 and 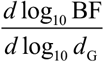 (Eq. (F20)) as the functions of parameter *d*_G_ and *λ*_ini,max_ (F), *λ*_rec,max_ (G), *λ*_rel,max_ (H). The bottom of (G) is a magnified display of a portion of the top of (G). (I-J) log_10_ MBS and log_10_BF as the functions of log_10_ (*d*_G_), where green dashed lines represent theoretical prediction (Eq. (E7)), orange solid lines represent binary approximation (Eq. (E7)), and red solid lines represent power-law approximation (Eq. (F18)). Other parameters are set as the default values in SI Table 2.

**SI Figure 15.**
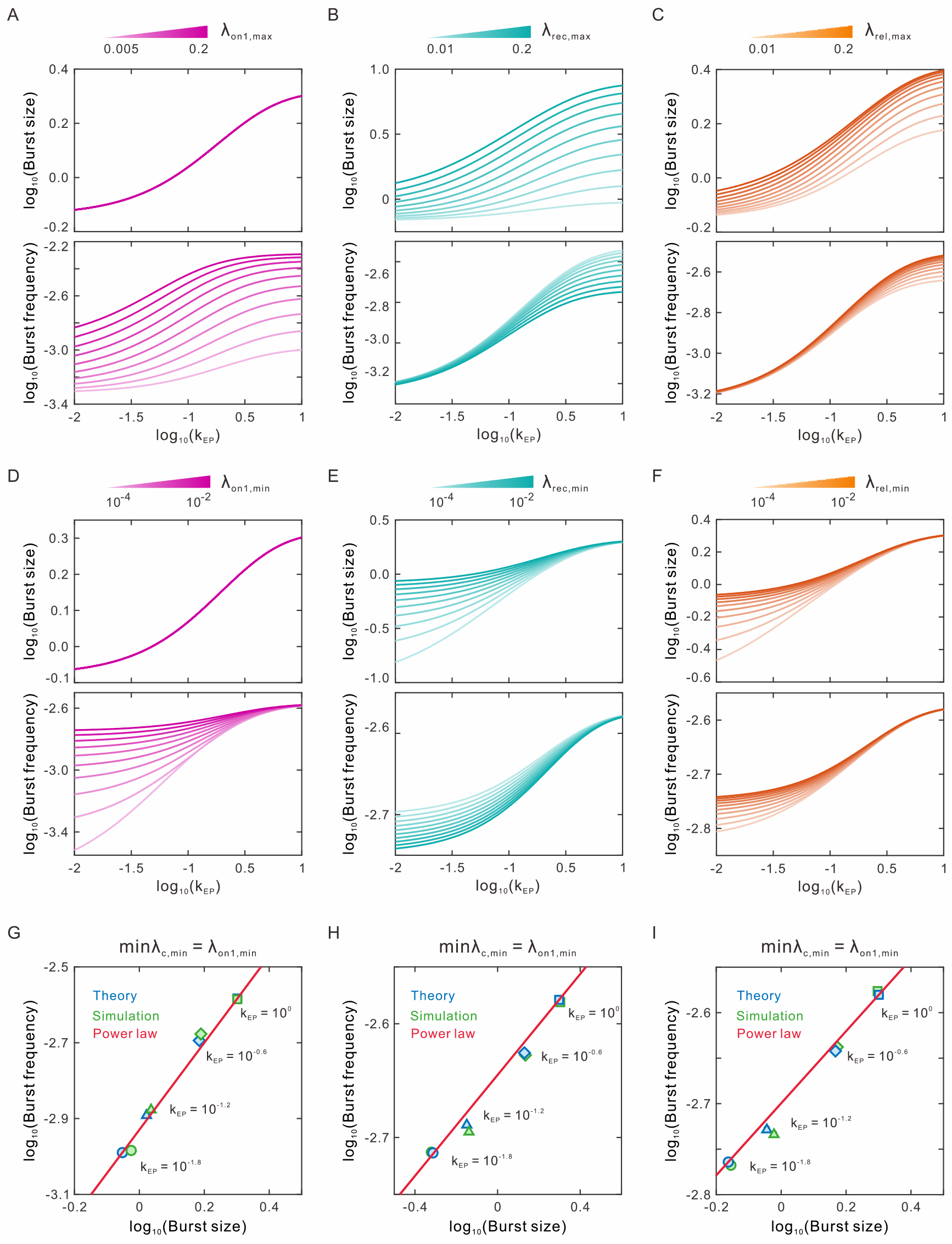
Effects of model parameters and upstream and downstream scales on burst size and burst frequency on logarithmic scale. (A-C) Influences of model parameters *λ*_on1,max_ (A),*λ*_rec,max_ (B), *λ*_rel,max_ (C) on burst size (top) and burst frequency (bottom), where other parameter values are the same as the default settings in SI Table 3. (D-F) Influences of min *λ*_*c*,min_ on burst size (top) and burst frequency (bottom). 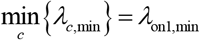 in (D) with fixed *λ*_rec,min_ = 0.01, 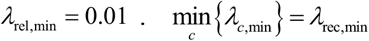 in (E) with fixed *λ*_on1,min_ = 0.01, 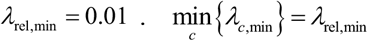 (F) with fixed *λ*_on1,min_ = 0.01, *λ*_rec,min_ = 0.01. Other parameter values are the same as the default settings in SI Table 3. (G-I) Influences of different values of *k*_EP_ on log_10_(MBS) and log_10_(BF). The red solid lines represent the approximate linear relationship between log_10_(MBS) and log_10_(BF). Parameter values are set as *λ*_on1,min_ = 0.004, *λ*_rec,min_ = 0.01, *λ*_rel,min_ = 0.01 in (G), *λ*_on1,min_ = 0.01, *λ*_rec,min_ = 0.004, *λ*_rel,min_ = 0.01 in (H) and *λ*_on1,min_ = 0.01, *λ*_rec,min_ = 0.01, *λ*_rel,min_ = 0.04, in (I). Other parameter values are the default values as in SI Table 2.

**SI Figure 16.**
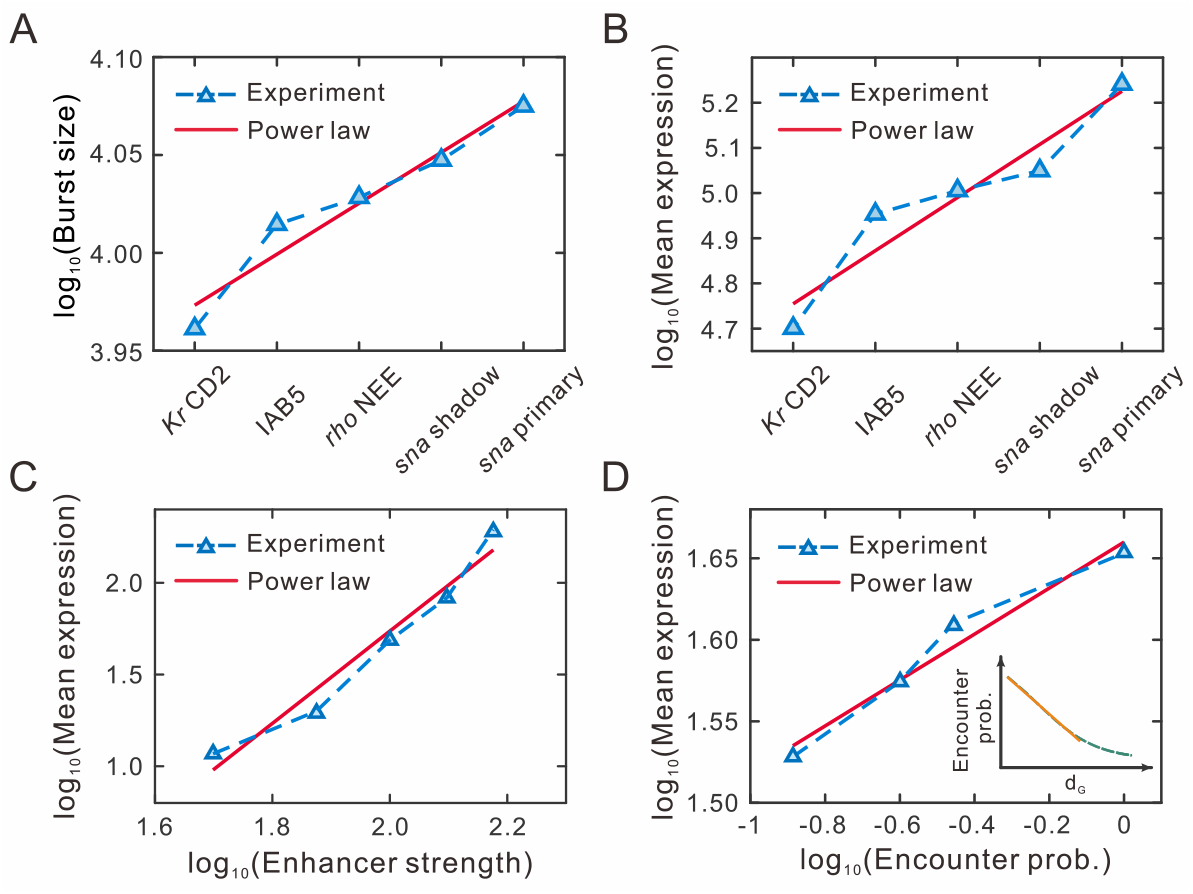
Experimental data verify the validity of power-law behaviors on burst size and burst frequency as well as mean expression level. (A-B) Plot of experimental data from (39). The labels of the *x*-axis represent the *sna* shadow, *sna* primary, *rho* NEE, IAB5, and *Kr* CD2 enhancers, respectively. (C) Log-log plot of experimental data from (118). (D) Log-log plot of experimental data from (120). Insert figure shows the log-log relationship between *d*_G_ and encounter probability. The yellow line indicates an approximate power-law relationship for smaller *d*_G_.

**SI Figure 17.**
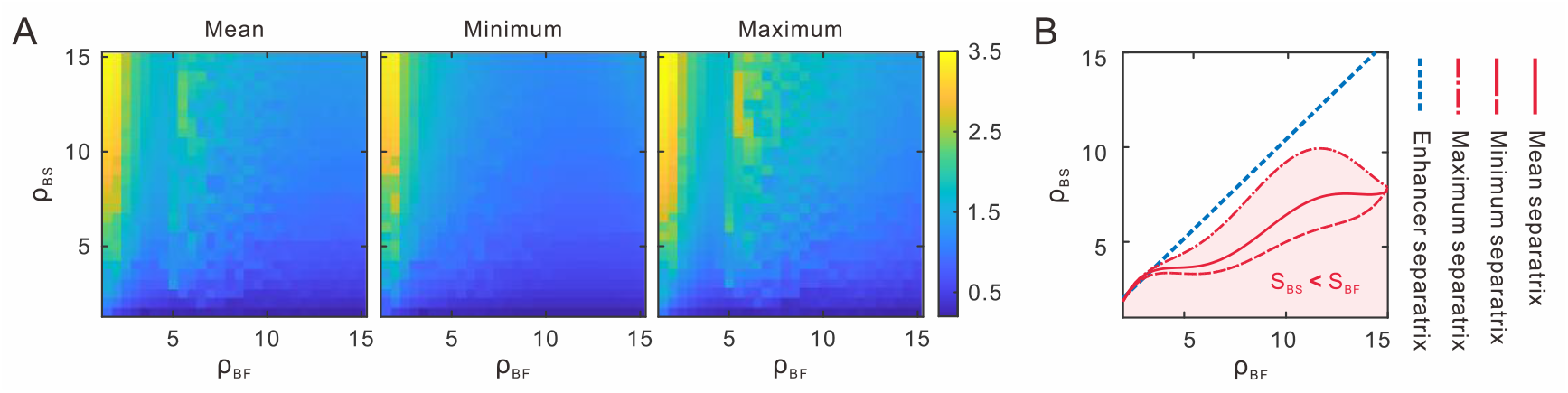
Effects of model parameters and upstream and downstream scales on ratio *S*_BF_/*S*_BS_ in the (*ρ*_BF_, *ρ*_BS_) plane. (A-C) Mean slope ratio for (A), minimum slope ratio for (B) and maximum slope ratio for (C) are obtained after the *S*_BF_/*S*_BS_ is calculated according to Eq. (F17) and Eq. (F14). (D) A phase diagram illustrates the size boundaries of the slopes of burst size and burst frequency in the (*ρ*_BF_, *ρ*_BS_) plane. The red lines are the separatrixes of data in (A). The blue dashed line is the separatrix in the case of enhancer deletion. Other parameter values are the default values in SI Table 2.

**SI Figure 18.**
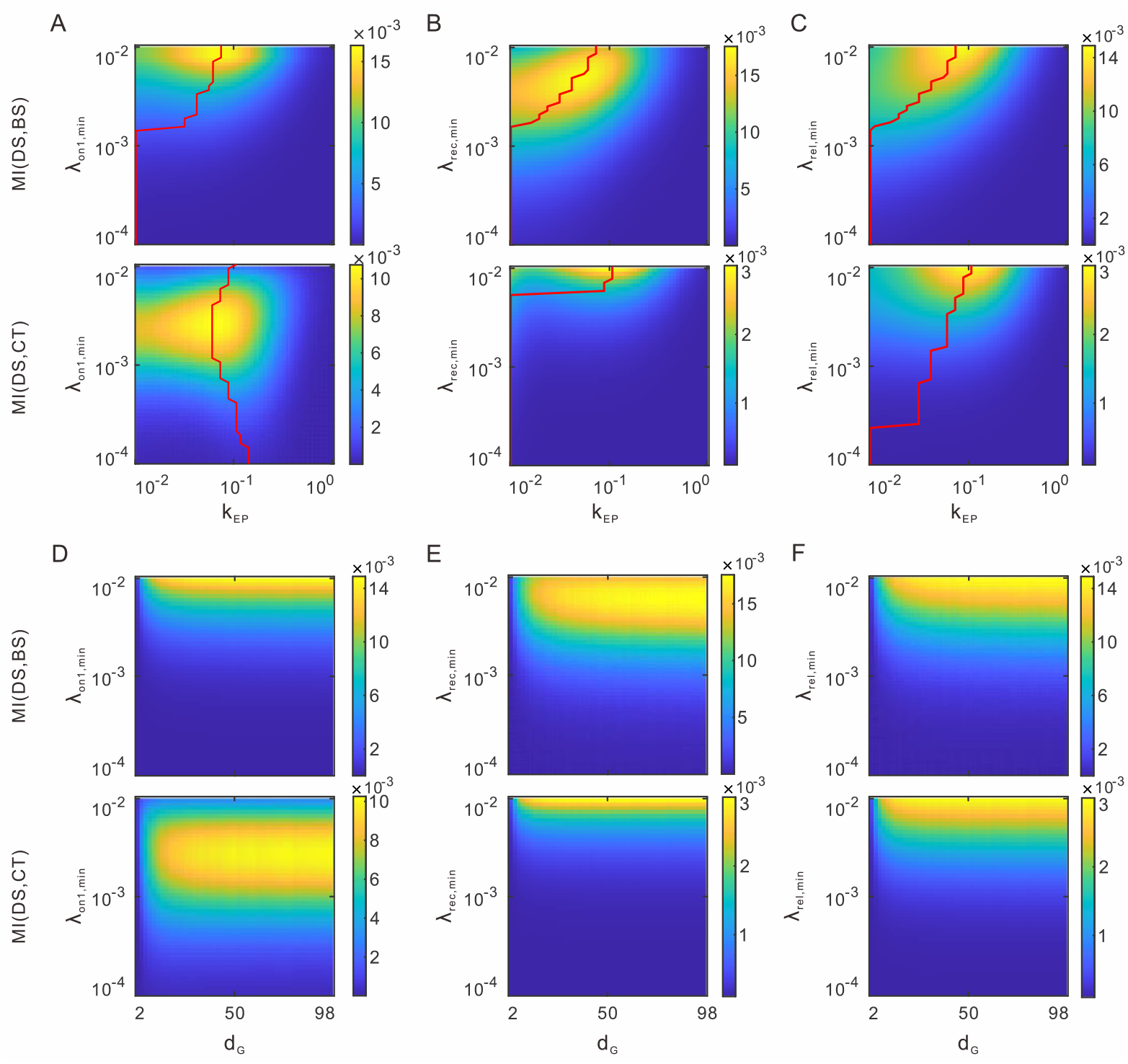
Influence of model parameters on mutual information. (A-C) Heatmap shows the joint effects of communication strength *k*_EP_ and minimum rate *λ*_on1,min_ for fixed *λ*_rec,min_ = 0.01, *λ*_rel,min_ = 0.01 in (A), *λ*_rec,min_ for fixed *λ*_on1,min_ = 0.01, *λ*_rel,min_ = 0.01 in (B), *λ*_rel,min_ for fixed *λ*_on1,min_ = 0.01, *λ*_rec,min_ = 0.01 in (C) on MI(*DS,BS*) (top) and MI(*DS,CT*) (bottom). Parameter values in (A) are the same as the default settings in SI Table 3. The red line shows the values of *k*_EP_ corresponding to the maximum mutual information. (D-F) Heatmap shows the joint effects of genomic distance *d*_G_ and minimum rate *λ*_on1,min_ for fixed *λ*_rec,min_ = 0.01, *λ*_rel,min_ = 0.01 in (D), *λ*_rec,min_ for fixed *λ*_on1,min_ = 0.01, *λ*_rel,min_ = 0.01 in (E) *λ*_rel,min_ for fixed *λ*_on1,min_ = 0.01, *λ*_rec,min_ = 0.01 in (F) on MI(*DS,BS*) (top) and MI(*DS,CT*) (bottom). Other parameter values are the same as the default settings in SI Table 2.

**SI Figure 19.**
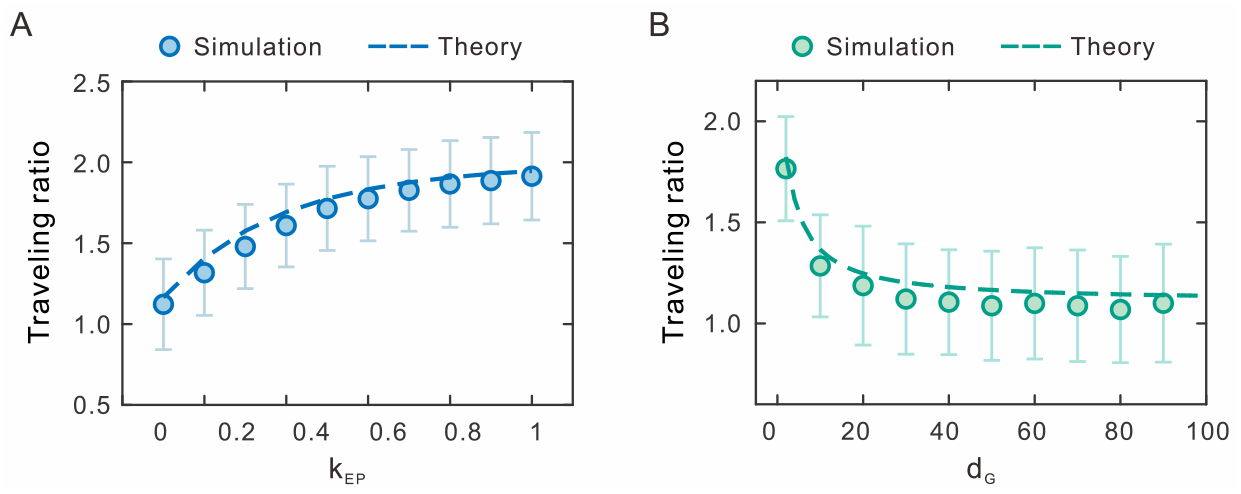
Influence of parameters *k*_EP_ (A) and *d*_G_ (B) on traveling ratio. The parameter values are set as *λ*_on2_ = 0.003, *λ*_off2_ = 0.006, *λ*_on1,min_ = 0.003, *λ*_on1,max_ = 0.020, *λ*_rec,min_ = 0.010, *λ*_rec,max_ = 0.030, *λ*_rel,min_ = 0.008, *λ*_rel,min_ = 0.020, 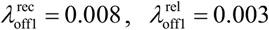. Other parameter values are the default values as in SI Table 2.

## Notes

### Competing Interest Statement

The authors have declared no competing interest.

